# North American Breeding Bird Survey status and trend estimates to inform a wide-range of conservation needs, using a flexible Bayesian hierarchical generalized additive model

**DOI:** 10.1101/2020.03.26.010215

**Authors:** Adam C. Smith, Brandon P.M. Edwards

**Author notes:** Author Contributions: ACS conceived the ideas and designed methodology; BPME and ACS analyzed the data; ACS led the writing of the manuscript. ACS and BPME contributed critically to the drafts and gave final approval for publication.

## Abstract

The status and trend estimates derived from the North American Breeding Bird Survey (BBS), are critical sources of information for bird conservation. However, the estimates are partly dependent on the statistical model used. Therefore, multiple models are useful because not all of the varied uses of these estimates (e.g. inferences about long-term change, annual fluctuations, population cycles, recovery of once declining populations) are supported equally well by a single statistical model. Here we describe Bayesian hierarchical generalized additive models (GAM) for the BBS, which share information on the pattern of population change across a species’ range. We demonstrate the models and their benefits using data a selection of species; and we run a full cross-validation of the GAMs against two other models to compare predictive fit. The GAMs have better predictive fit than the standard model for all species studied here, and comparable predictive fit to an alternative first difference model. In addition, one version of the GAM described here (GAMYE) estimates a population trajectory that can be decomposed into a smooth component and the annual fluctuations around that smooth. This decomposition allows trend estimates based only on the smooth component, which are more stable between years and are therefore particularly useful for trend-based status assessments, such as those by the IUCN. It also allows for the easy customization of the model to incorporate covariates that influence the smooth component separately from those that influence annual fluctuations (e.g., climate cycles vs annual precipitation). For these reasons and more, this GAMYE model is a particularly useful model for the BBS-based status and trend estimates.

**LAY SUMMARY:**

- The status and trend estimates derived from the North American Breeding Bird Survey are critical sources of information for bird conservation, but they are partly dependent on the statistical model used.
- We describe a model to estimate population status and trends from the North American Breeding Bird Survey data, using a Bayesian hierarchical generalized additive mixed-model that allows for flexible population trajectories and shares information on population change across a species’ range.
- The model generates estimates that are broadly useful for a wide range of common conservation applications, such as IUCN status assessments based on trends or changes in the rates of decline for species of concern; and the estimates have better or similar predictive accuracy to other models., and

## INTRODUCTION

Estimates of population change derived from the North American Breeding Bird Survey are a keystone of avian conservation in North America. Using these data, the Canadian Wildlife Service (CWS, a branch of Environment and Climate Change Canada) and the United States Geological Survey (USGS) produce national and regional status and trend estimates (estimates of annual relative abundance and rates of change in abundance, respectively) for 300-500 species of birds (Smith et al. 2019, Sauer et al. 2014). These estimates are derived from models designed to account for some of the sampling imperfections inherent to an international, long-term field survey, such as which sites or routes are surveyed in a given year and variability among observers (Sauer and Link 2011, Smith et al. 2014). Producing these estimates requires significant analytical expertise, time, and computing resources, but they are used by many conservation organizations and researchers to visualize, analyze, and assess the population status of many North American bird species (e.g., Rosenberg et al. 2017, NABCI Canada 2019, Rosenberg et al. 2019).

While the estimates of status and trend from the BBS serve many different purposes, not all uses are equally well supported by the standard models, and so there is a need for alternative models and for a continual evolution of the modeling. Different conservation-based uses of the BBS status and trend estimates relate to different aspects of population change, including long-term trends for overall status (Partners in Flight, 2019), short-term trends to assess extinction-risk (IUCN 2019), changes in population trends to assess species recovery (Environment Climate Change Canada, 2016), or annual fluctuations (Wilson et al., 2018). Each one of these uses relies on different parameters, or spatial and temporal variations in those parameters, and no single model can estimate all parameters equally well. This is not a criticism; it is true of any single model. For example, the standard model used between 2011 and 2017 in the United States and 2011 and 2016 in Canada, is essentially a Poisson regression model, which estimates population change using random year-effects around a continuous slope in a Bayesian hierarchical framework (Sauer and Link 2011, Smith et al. 2014). These slope and year-effects are well suited to estimating annual fluctuations around a continuous long-term change, but the model tends to be conservative when it comes to estimating changes in a species’ population trend (e.g., population recovery after decline), or population cycles (Fewster et al. 2000, Smith et al. 2015). Similarly, short-term trends (e.g., the last 10-years of the time-series) derived from this standard model incorporate information from the entire time-series (i.e., the slope component of the model). For many purposes, this is a reasonable and useful assumption, which guards against extreme and imprecise fluctuations in short-term trends. However, for assessing changes in trends of a once-declining species, such as the recovery of a species at risk (Environment and Climate Change Canada, 2016), this feature of the model is problematic.

Generalized Additive Models (GAM, Wood 2017) provide a flexible framework for tracking changes in populations over time, without any assumptions about a particular temporal pattern in population change (Fewster et al., 2000, Knape 2016). The semi-parametric smooths can fit almost any shape of population trajectory, including stable populations, constant rates of increase or decrease, cycles of varying frequency and amplitude, or change points in population trends (Wood 2017). Furthermore, the addition of new data in subsequent years has relatively little influence on estimates of population change in the earlier portions of the time-series. By contrast, the slope parameter in the standard models effectively assumes that there is some consistent rate of change. As a result, to the extent that the slope parameter influences the estimated trajectory, estimates of the rate of a species population change in the early portion of the time series (e.g., during the 1970s or 80s) can change in response to the addition of contemporary data and recent rates of population change.

GAMs also provide a useful framework for sharing information on the shape and rate of population change across a species’ range. The GAM smoothing parameters can be estimated as random effects within geographic strata, thus allowing the model to share information on the shape of the population trajectory across a species range. In the terminology of Pedersen et al. 2019, this hierarchical structure on the GAM parameters would make our model a “HGAM” (Hierarchical Generalized Additive Model). However, it also includes random effects for parameters not included in the smooth and could therefore be referred to as a GAMM (Generalized Additive Mixed Model), in the terminology of Wood 2017. Similarly in the standard model, the slope parameters can be estimated as random effects and share information among strata, which improves estimates of trend for relatively data-sparse regions (Link et al. 2017, Smith et al. 2019). Although recent work has shown that the standard model is, for many species, out-performed by a first-difference model (Link et al. 2020), the population change components of the first-difference model (Link et al. 2017), include no way to share information on population change in space and so population trajectories are estimated independently among strata. Of course, for some conservation uses, this independent estimation of population trajectories might be critical (e.g., if one were interested specifically in estimating the differences in trends among provinces or states), and in these situations the sharing of information could be problematic.

Trend estimates (interval-specific rates of mean annual population change, Sauer and Link 2011, Link et al. 2020) derived from the inherently smooth temporal patterns generated by GAMs are well suited to particularly common conservation uses, such as assessments of trends in populations from any portion of a time-series, as well as assessments of the change in the trends over time. For example, the population trend criteria of the IUCN (IUCN 2019) or Canada’s national assessments by the Committee on the Status of Endangered Wildlife in Canada (COSEWIC) are based on rates of change over 3 generations. For most bird species monitored by the BBS, this 3-generation time is approximately the same as the 10-year, short-term trends produced by the CWS and USGS analyses. Because of the inclusion of year-effects in the standard model, these short-term trends fluctuate from year to year, complicating the quantitative assessment of a species trend in comparison to the thresholds. Species trends may surpass the threshold in one year, but not in the next. The same end-point comparisons on estimates from a GAM will change much more gradually over time, and be much less dependent on the particular year in which a species is assessed.

In this paper, we describe a status and trend model that uses a hierarchical GAM to estimate the relative abundance trajectory of bird populations, using data from the BBS. This model allows for the sharing of information about a species’ population trajectory among geographic strata and for the decomposition of long- and medium-term population changes from annual fluctuations. We also compare the fit of the GAM, and a GAM-version that includes random year-effects (conceptually similar to Knape et al. 2016), to the fit of two alternative models for the BBS (Sauer and Link 2011, Smith et al. 2015, Link et al. 2020).

## METHODS

### Overview

We designed a Bayesian hierarchical model for estimating status and trends from the North American Breeding Bird Survey (BBS) that uses a Generalized Additive Model (GAM) smooth to estimate the medium- and long-term temporal components of a species population trajectory (i.e., changes occurring over time-periods ranging from 3-53 years). In the model, the parameters of the GAM smooths are treated as random-effects within the geographic strata (the spatial units of the predictions, intersections of Bird Conservation Regions and province/state/territory boundaries), so that information is shared on the shape of the population trajectory across the species’ range. In comparison to the non-Bayesian hierarchical GAMs (HGAM) in Pedersen et al. 2019, our model is most similar to the “GS” model, which has a global smooth in addition to group-level smooths with a similar degree of flexibility. We applied two versions of the GAM: one in which the GAM smooth was the only component modeling changes in abundance over time (GAM), and another in which random year effects were also estimated to allow for single-year departures from the GAM smooth (GAMYE, which is conceptually similar to the model described in Knape 2016).

For a selection of species, we compared estimates and predictive accuracy of our two models using the GAM smooth, against two alternative models that have been used to analyze the BBS data. We chose the main comparison species (Barn Swallow) because of the striking differences between trajectories from the SLOPE model and a number of non-linear models (Sauer and Link 2017, Smith et al. 2015). We added a selection of other species to represent a range of anticipated patterns of population change, including species with known change points in their population trajectories (Chimney Swift, Smith et al. 2015), and species with relatively more data and known large and long-term trends (Wood Thrush, Ruby-throated Hummingbird) and species with relatively fewer data and long-term changes (Canada Warbler, Cooper’s Hawk, and Chestnut-collared Longspur). Finally, we also added a few species with strong annual fluctuations and/or abrupt step-changes in abundance (Pine Siskin, Carolina Wren).

The BBS data are collected along roadside survey-routes that include 50 stops at which a 3-minute point count is conducted, once annually, during the peak of the breeding season (Robbins et al. 1986, Sauer et al. 2017, Hudson et al. 2017). All of the models here use the count of individual birds observed on each BBS route (summed across all 50 stops) in a given year by a particular observer. The four statistical models differed only in the parameters used to model changes in species relative abundance over time. We used 15-fold cross validation (Burman 1983) to estimate the observation-level, out-of-sample predictive accuracy of all four models (Link et al. 2020, Vehtari et al. 2017). We chose this 15-fold cross-validation approach because although full leave-one-out (loo) cross-validation minimizes bias and variance of the estimate of predictive accuracy (Zhang and Yang 2015), the size of the BBS dataset makes this impractical (Link et al. 2017), and cross-validation with k > 10 is a reasonable approximation of loo (Kohavi 1995, Vehtari et al. 2017). We compared the overall predictive accuracy among the models, and we explored the spatial and temporal variation in predictive accuracy in depth.

Using the cross-validation, we have compared four alternative BBS models, all of which have the same basic structure:

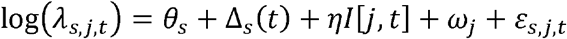

The models treat the observed BBS counts as overdispersed Poisson random variables, with mean *λ*_*s,j,t*_ (i.e., geographic stratum *s*, observer and route combination *j*, and year *t*). The means are log-linear functions of stratum-specific intercepts (*θ*_*s*_, estimated as fixed effects and with the same priors following Smith et al. 2014), observer-route effects (*ω* _*j*_, estimated as random effects and with the same priors following Sauer and Link 2011), first-year startup effects for a observer (*η*, estimated as fixed effects and with the same priors following Sauer and Link 2011), a count-level random effect to model overdispersion (*ε*_*s,j,t*_, estimated using heavy-tailed, t-distribution and with the same priors following Link et al. 2020), and a temporal component estimated using a function of year, which varies across the four models (Δ_*s*_(*t*)). The models here only varied in their temporal components (Δ_*s*_(*t*)).

### Bayesian hierarchical GAMs

#### GAM

The main temporal component Δ_*s*_ (*t*) in the GAM was modeled with a semi-parametric smooth, estimated following Crainiceanu et al (2005) as

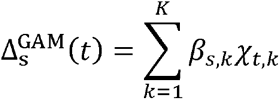

where *K* is the number of knots, *χ*_*t,k*_ is the year *t* and *k*th entry in the design matrix X (defined below), and *β*_*s*,_.is the -length vector of parameters that control the shape of the trajectory in stratum *s*. Each *β*_*s,k*_ is estimated as a random effect, centered on a mean across all strata (a hyperparameter B_*k*_)

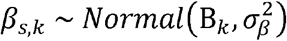

and

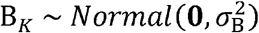

where the variance 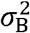 acts as the complexity penalty, shrinking the complexity and the overall change of the mean trajectory towards a flat line). It would be possible to add an additional slope parameter, as was done in Crainiceanu et al. 2005, but we have found that the BBS data for most species are insufficient to allow for the separate estimation of the linear component to population change and the additive smooth. In addition, we see little benefit to including a linear component because the assumptions required to include a constant linear slope for a 53 year time-series are unlikely to be met for any continental-scale population. In combination, these variance parameters 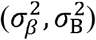 control the complexity penalty of the species trajectories and the variation in pattern and complexity among strata and were given the following priors, following advice in Crainiceanu et al (2005):

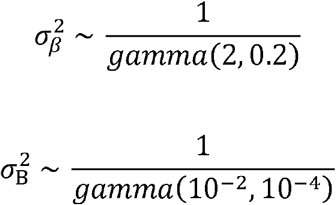

These prior parameters were chosen to ensure that the priors are sufficiently vague that they are overwhelmed by the data, particularly for 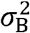 that controls the shape of the survey-wide trajectory (Crainiceanu et al 2005). We have so far had good results across a wide range of species using these priors, and in tests of alternative priors there is no effect on posterior estimates (Supplemental Figure S9). For example, estimates of B_*K*_ and *σ*_***B***_ for Chestnut-collared Longspur (a relatively data-poor species) are unchanged even if using a much more restrictive prior on *σ*_***B***_ that places 99% of the prior density for *σ*_***B***_ below 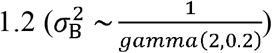. However, these variance priors are an area of ongoing research, aimed at improving the efficiency of the MCMC sampling.

The design matrix for the smoothing function (X) has a row for each year, and a column for each of *K* knots. The GAM smooth represented a 3^rd^-degree polynomial spline: *χ*_*t,k*_ = | *t*′ *– t*′_*k*_ |^3^, and was calculated in R, following Crainiceanu et al (2005). We centered and re-scaled the year-values to improve convergence, so that *t*′ = (*t* – *midyear*)/*T*, where midyear is the middle year of the time-series, and *T* is the number of years in the time-series. Here, we have used 13 knots (*K*= 13), across the 53-year time-series of the BBS (1966-2018), which results in approximately one knot for every 4 years in the time-series. With this number of knots, we have found that the 53-year trajectories are sufficiently flexible to capture all but the shortest-term variation (i.e., variation on the scale of 3-53 years, but not annual fluctuations). Models with more knots are possible but in the case of a penalized smooth, the overall patterns in the trajectory will be very similar, as long as a sufficient number of knots is allowed (Wood 2017). The number of knots could be customized in a species-specific approach, however because we are looking for a general model structure that can be applied similarly across the >500 species in the BBS, we have fixed the number of knots at 13. Our approach relies on the shrinkage of the smoothing parameters (B,β) to ensure that the trajectories are only as complex as the data support, and the limited number of knots constrains the complexity of the additive function (Wood 2017, Fewster et al. 2000).

### GAMYE

The GAMYE was identical to the GAM, with the addition of random year effects (*γ*_*t,s*_) estimated independently among strata, following Sauer and Link (2011) and Smith et al. (2015), as

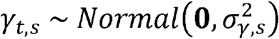

Where 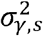 are stratum-specific variances. Thus, the temporal component for the GAMYE is given by

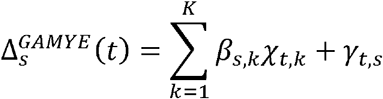

The GAMYE trajectories are therefore an additive combination of the smooth and random annual fluctuations. The smooth components of the trajectory in the GAMYE are generally very similar to those from the GAM, but tend to be slightly less variable (i.e., less complex) because the year-effects components can account for single-year deviations from the longer-term patterns of population change. The full trajectories from the GAMYE (smooth plus the year-effects) generally follow the same overall pattern as the GAM estimates, and include abrupt single-year changes in abundance, which increases the capacity to model step-changes in abundance.

### Alternative models

For a selection of species, we compared the predictions and predictive accuracy of the two GAMs against two alternative models previously used for the BBS.

#### SLOPE

The SLOPE model includes a slope parameter and random year-effects to model species trajectories. It is a linear year-effects model currently used by both the CWS (Smith et al. 2014) and the USGS (Sauer et al. 2017) as an omnibus model to supply status and trend estimates from the BBS (essentially the same as model SH, the Slope model with Heavy-tailed error in Link et al 2017). The temporal component in the SLOPE model is

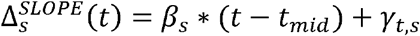

#### DIFFERENCE

The first-difference model (DIFFERENCE) is based on a model described in Link and Sauer (2015) and models the temporal component as

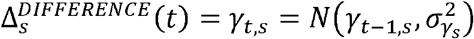

The DIFFERENCE model includes year-effects that follow a random walk prior from the first year of the time-series, by modeling the first-order differences between years as random effects with mean zero and an estimated variance.

All analyses in this paper were conducted in R (R Core Team, 2019), using JAGS to implement the Bayesian analyses (Plummer 2003), and an R-package *bbsBayes* (Edwards and Smith 2020) to access the BBS data and run all of the models used here. We used the same number of burn-in iterations (10 000), thinning-rate (1/10), chains (3), and number of saved samples from the posterior (3000) to estimate trends and trajectories for all models. We examined trace plots and the Rhat statistic to assess convergence. The graphs relied heavily on the package *ggplot2* (Wickham 2016). BUGS-language descriptions of the GAM and GAMYE, as well as all the code and data used to produce the analyses in this study, are archived online (see Data Depository in Acknowledgements).

### Cross-validation

We used a temporally and spatially stratified v-fold cross-validation (Burman 1983, often termed “k-fold”, but here we use Berman’s original “v-fold” to distinguish it from “k” which is often used to describe the number of knots in a GAM) with v = 15, where we held-out random sets of counts, stratified across all years and strata so that each of the v-folds included some observations from almost every combination of strata and years. We did this by randomly allocating each count within a given stratum and year to one of the 15 folds. We chose this approach over a leave-one-out cross-validation (loo) approach using a random subset of counts (e.g., Link et al. 2017) because we wanted to assess the predictive success across all counts in the dataset, explore the temporal and spatial patterns in predictive success, and a full loo is not practical for computational reasons (see Link et al. 2017). We could also have chosen to conduct a structured cross-validation (Roberts et al. 2017), but cross-validation in a Bayesian context has particularly large computational requirements; there are multiple levels of dependencies in the BBS data (dependences in time, space, and across observers); and models being compared vary in the way they treat some of those dependencies (models that share information differently in space and/or time). Therefore, we chose a relatively simple non-structured approach where the folds are balanced in time and space, and for a given species were identical across all models compared. We followed a similar procedure to that outlined in Link et al. (2017) to implement the cross-validation in a parallel computing environment, using the R-package foreach (Microsoft and Weston 2019). We used the end-values from the model-run using the full dataset as initial values in each of the 15 cross-validation runs, ran a short burn-in of 1000 samples, then used a draw of 3000 samples of the posterior with a thinning rate of 1/10 spread across 3 chains. We did not calculate WAIC because previous work has shown that WAIC does not approximate loo well for the BBS data (Link et al. 2017, Link et al. 2020).

We used the estimated log predictive density (elpd_*i,M*_) to compare the observation-level, out-of-sample predictive success of all four models (Link et al. 2020, Vehtari et al. 2017). For a given model *M*, elpd is the estimated log posterior density for each observation *i*, for the model *M* fit to all data except those in the set *ν* that includes *i* (*Y* _-*ν,i*∈ *ν*_). That is,

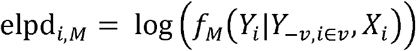

Larger values of elpd indicate better predictive success, that is a higher probability of the observed data given the model *M*, the estimated parameters, the vector of covariates for observation i, such as the year, observer-route, etc. (*X*_*i*_), and all of the data used to fit the model (*Y* _-*ν,i*∈ *ν*_).

We have not summed elpd values to generate BPIC values (Link et al. 2020); rather, we have compared model-based estimates of mean difference in elpd between pairs of models. We modeled the elpd values so that we could account for the imbalances in the BBS data in time and space (i.e., the variation in number of counts among strata and years). The raw sum of the elpd values would give greater weight to the regions with more data and to the recent years in the time-series, which have more counts. Therefore, expanding on the approach in Link et al. 2020 that used a z-score the estimated the significance of the difference in fit between two models, we used a hierarchical model to estimate the mean difference in predictive fit 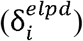. We first calculated the difference in the elpd of each observed count (*Y*_*i*_) under models 1 and 2, as 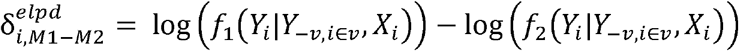, so that positive values of 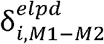 indicate more support for model 1. We then analysed these 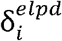 values using an additional Bayesian hierarchical model, with random effects for year and strata to account for the variation in sampling effort in time and space. These random effects account for the imbalances in the BBS-data among years and regions, and the inherent uncertainty associated with any cross-validation statistic (Vehtari et al. 2017, and Link et al. 2017). This model treated the elpd differences for a count from a given year *t* and stratum 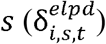 as having a t-distribution with an estimated variance 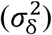 and degrees of freedom (*ν*). That is,

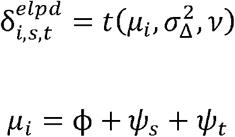

From the model, *ϕ* was our estimate of the overall comparison of the mean difference in predictive fit for Model 1 – Model 2 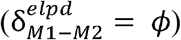 was the estimate of the mean difference in stratum *s*, and *ϕ + ψ*_*t*_ was the estimated difference in year *t*. The year and stratum effects (*ψ*_*t*_+ *ψ*_*t*_) were estimated as random effects with a mean of zero and estimated variances given uninformative inverse gamma priors. We used this t-distribution as a robust estimation approach, instead of the z-score approach used by Link et al. (2020) because of the extremely heavy tails in the distribution of the 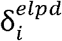 values (Supplemental Figure S7). Given these heavy tails, a statistical analysis assuming a normal distribution in the differences would give an inappropriately large weight to a few counts where the prediction differences were extremely large in magnitude (Gelman et al. 2014). In essence, our model is simply a “robust” version of the z-score approach (Lange et al. 1989) with the added hierarchical parameters to account for the spatial and temporal imbalance in the BBS data.

### Trends and population trajectories

For all models, we used the same definition of trend following Sauer and Link (2011) and Smith et al. (2015); that is, an interval-specific geometric mean of proportional changes in population size, expressed as a percentage. Thus, the trend estimate for the interval from year a (*t*_*a*_) through year b (*t*_*b*_) is given by

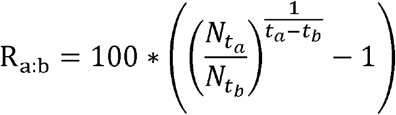

where *N*. represents the annual index of abundance in a given year (see below). Because this estimate of trend only considers the annual abundance estimates in the years at either end of the trend period, we refer to this estimate as an end-point trend. For the GAMYE model, we decomposed the trajectory (i.e., the series of annual indices of abundance) into long- and medium-term components represented by the GAM smooth and annual fluctuations represented by the random year-effects. This decomposition allowed us to estimate two kinds of trend estimates: R_a:b_ that include all aspects of the trajectory, and 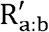 that removes the annual fluctuations, including only the GAM smooth components.

Population trajectories are the collection of annual indices of relative abundance across the time series. These indices approximate the mean count on an average BBS route, conducted by an average observer, in a given stratum and year. For all the models here, we calculated these annual indices for each year *t* and stratum *s* following Smith et al. (2019) as

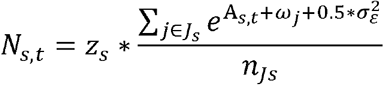

where each *N*_*s,t*_ are exponentiated sums of the relevant components of the model (A_*s,t*_), observer-route effects (*ω*_*j*_), and count-level extra-Poisson variance 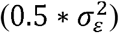, averaged over count-scale predictions across all of the *n*_*Js*_ observer-routes *j* in the set of observer-route combinations in stratum *s*(*J*_*s*_), and then multiplied by a correction factor for the proportion of routes in the stratum on which the species has been observed (*Z*_*s*_, i.e., the proportion of routes on which the species has been observed, on all other routes species abundance is assumed to equal zero and they are excluded from the model, see Sauer and Link 2011). This is slightly different from the approach described in Sauer and Link (2011) and Smith et al. (2015), and an area of ongoing research. We have found that this different annual index calculation ensures that the annual indices are scaled more similarly to the observed mean counts, which can affect the relative weight of different strata in trends estimated for broader regions (e.g., continental and national trends), but it has no effect of the trends estimated within each stratum and no effect on the cross-validation results presented here. For a discussion on the differences between these two ways of calculating annual indices, refer to the Supplemental Material.

For the GAMYE model, we calculated two versions of the species trajectory (*N*_*s*_): one that included the annual variation in the trajectory,

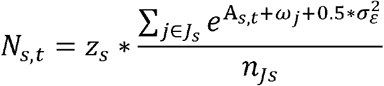

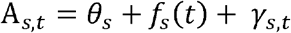

and a second that excluded the annual variations, including only the smoothing components of the GAM to estimate the time-series,

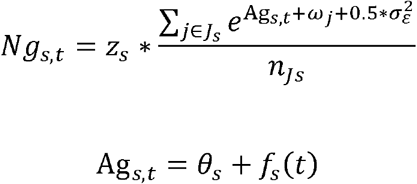

We calculated population trajectories and trends from the GAMYE model using both sets of annual indices (*N*_*s,t*_ and *Ng*_*s,t*_). When comparing predictions against the other models, we use the N_*s,t*_ values to plot and compare the population trajectories (i.e., including the year-effects), and the *Ng*_*s,t*_ values to calculate the trends (i.e., removing the year-effect fluctuations).

## RESULTS

### Model predictions

Population trajectories from the GAM, GAMYE, and DIFFERENCE are very similar. All three of these models suggest that Barn Swallow populations increased from the start of the survey to approximately the early 1980s, compared to the SLOPE model predictions that show a relatively steady rate of decline (Figure 1). The trajectories for all species from both GAMs and the DIFFERENCE model were less linear overall than the SLOPE model trajectories and tended to better track nonlinear patterns, particularly in the early years of the survey and often in more recent years as well (Figure 1, Supplemental Materials Figures S1 and S6). GAM and GAMYE trajectories vary a great deal among the strata, particularly in the magnitude and direction of the long-term change (Figure 2 for Barn Swallow). However, there are also many similarities among the strata, in the non-linear patterns that are evident in the continental mean trajectory (e.g., the downward inflection in the early 1980s in Figure 2 and Supplemental Materials Figure S2). Figure 3 shows the estimate trajectories for Barn Swallow in the 6 strata that make up BCR 23 from the GAMYE, DIFFERENCE, and SLOPE models. The GAMYE estimates suggest that the species’ populations increased in the early portion of the time series in all of the strata, and this is a pattern shared with the continental mean trajectory for the species (Figure 2). By contrast, the estimates from the SLOPE model only show an increase in the stratum with the most data, (i.e., the most stacked grey dots along the x-axis indicating the number of BBS routes contributing data in each year, US-WI-23), the DIFFERENCE model shows more of the early increase in many strata, except those with the fewest data. In the other strata with fewer data the SLOPE trajectories are strongly linear and the DIFFERENCE trajectories are particularly flat in the early years with particularly few data. The cross-validation results suggest that for Barn Swallow, the GAMYE is preferred over the SLOPE model, and generally preferred (some overlap with 0) to the DIFFERENCE model (Figure 4), particularly in the early years of the survey (pre-1975, Supplemental Materials Figure S6). Finally, the general benefits of sharing information among strata on the shape of the population trajectory are evident for the GAM, GAMYE, and the SLOPE models in Figure 5, where there is no relationship between the sample size and the absolute value of the long-term trend for Cooper’s Hawk (more below).

**Figure 1.**
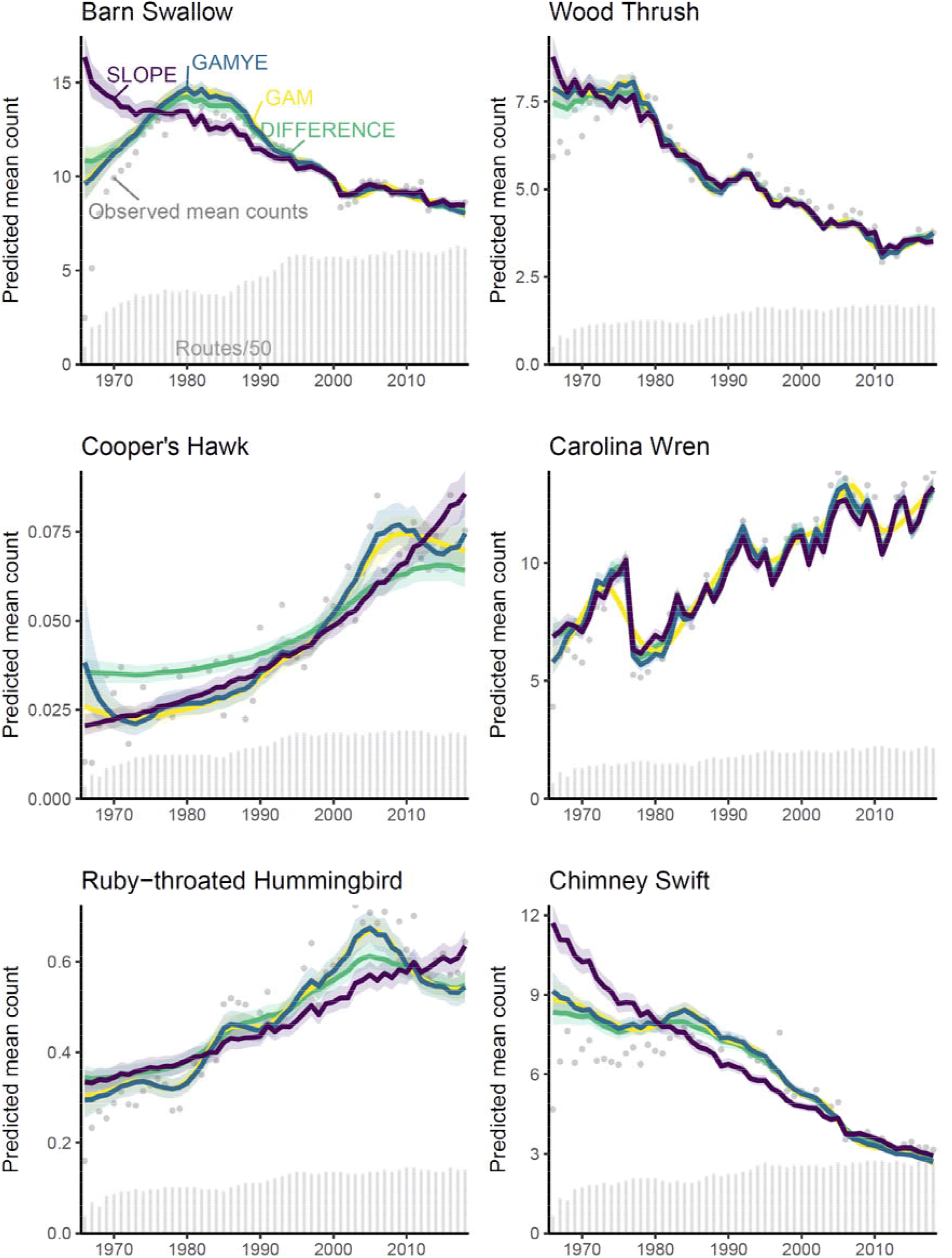
Survey-wide population trajectories for Barn Swallow (*Hirundo rustica*), Wood Thrush (*Hylocichla mustelina*), Cooper’s Hawk (*Accipiter cooperii*), Carolina Wren (*Thryothorus ludovicianus*), Ruby-throated Hummingbird (*Archilochus colubris*), and Chimney Swift (*Chaetura pelagica*), estimated from the BBS using two models described here that include a GAM smoothing function to model change over time (GAM and GAMYE) the standard regression-based model used for BBS status and trend assessments since 2011 (SLOPE), and a first-difference time-series model (DIFFERENCE). The stacked dots along the x-axis indicate the approximate number of BBS counts used in the model in each year; each dot represents 50 counts.

**Figure 2.**
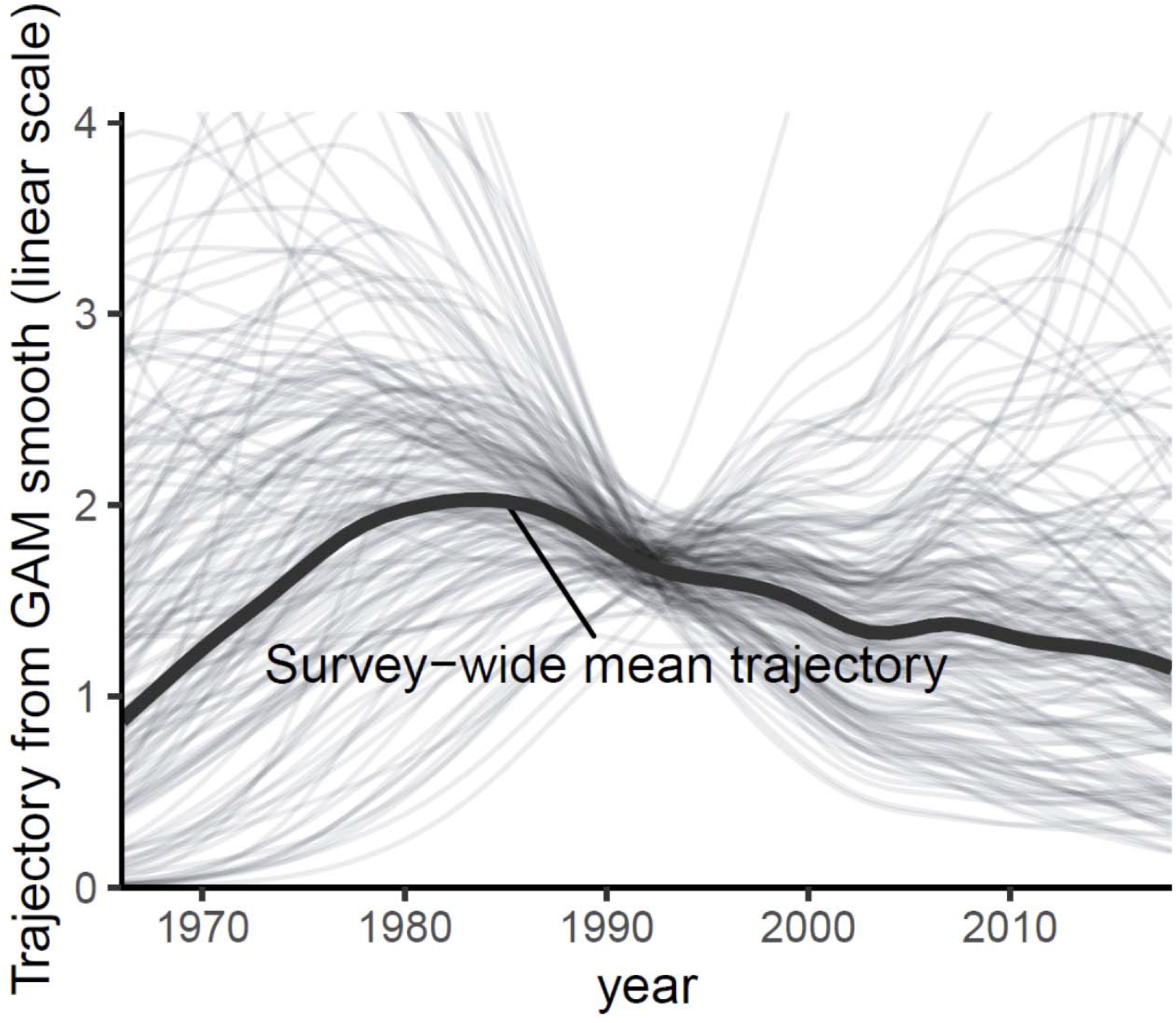
Variation among the spatial strata in the random-effect smooth components of the GAMYE model applied to Barn Swallow data from the BBS. Grey lines show the strata-level random-effect smooths, and the black lines shows the survey-wide mean trajectory.

**Figure 3.**
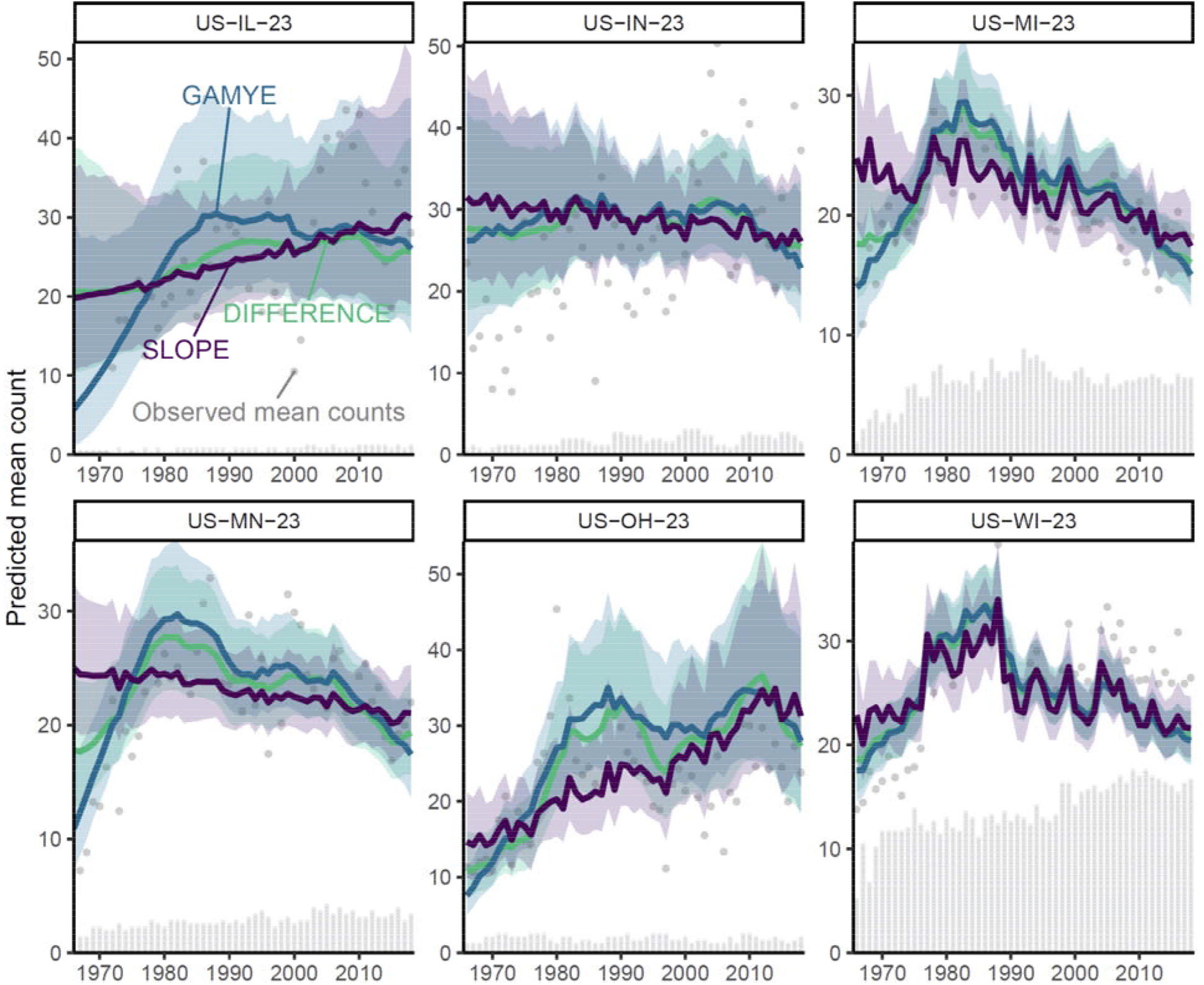
Stratum-level predictions for Barn Swallow population trajectories in BCR 23 from GAMYE against the predictions from the SLOPE and DIFFERENCE model. The similarity of the overall patterns in the GAMYE as compared to the SLOPE estimates, demonstrates the inferential benefits of the sharing of information among regions on the non-linear shape of the trajectory. In most strata the similar patterns of observed mean counts and the GAMYE trajectories suggests a steep increase in Barn Swallows across all of BCR 23 during the first 10-years of the survey. The GAMYE estimates show this steep increase in almost all of the strata, whereas the SLOPE predictions only show this pattern in the most data rich stratum, US-WI-23. The DIFFERENCE trajectories model the non-linear shapes well in all but the most data-sparse region (US-IL-23) and years (< 1972). The facet strip labels indicate the country and state-level division of BCR 23 that makes up each stratum. The first two letters indicate all strata are within the United States, and the second two letters indicate the state. The stacked dots along the x-axis indicate the number of BBS counts in each year and stratum; each dot represents one count.

**Figure 4.**
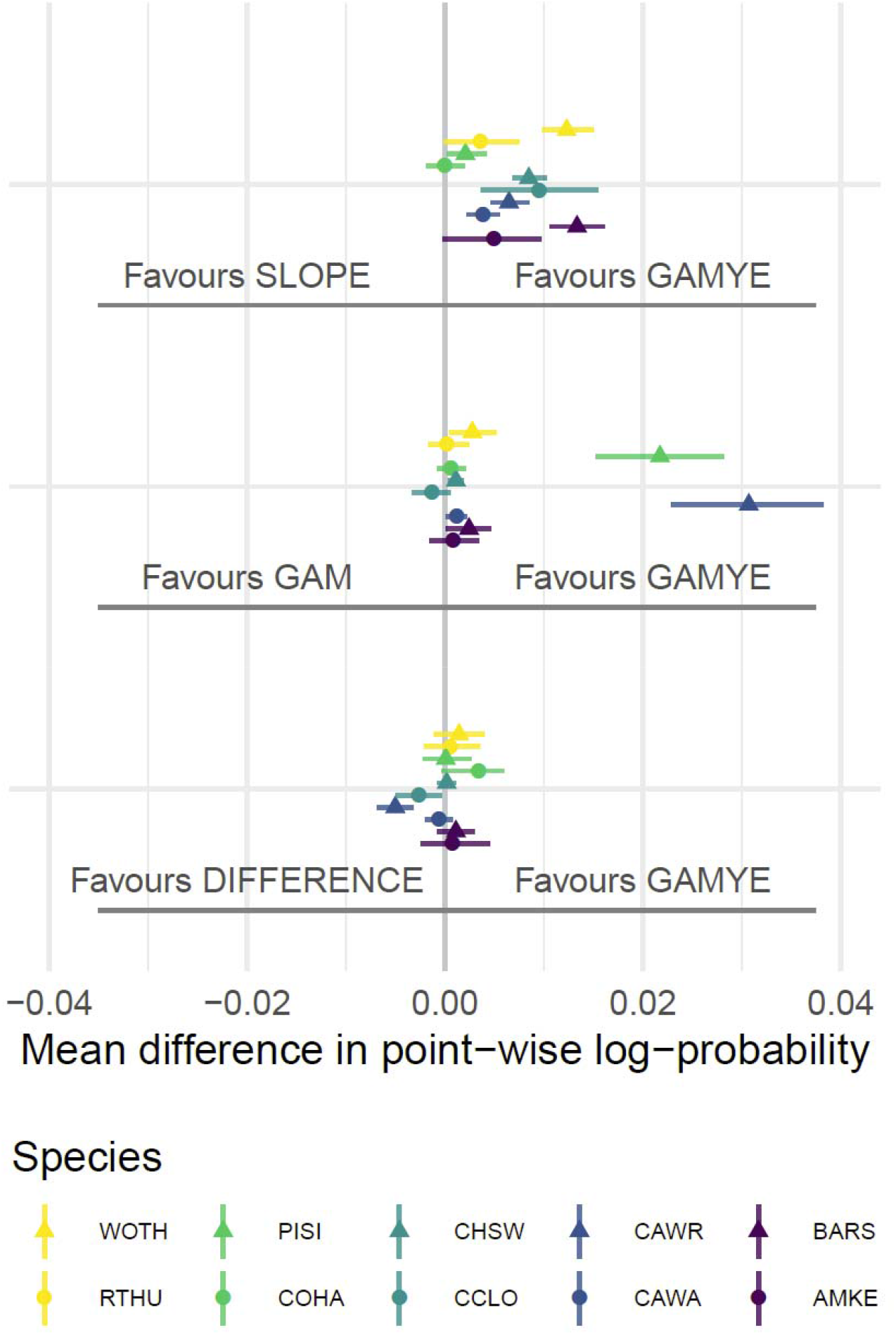
Overall differences in predictive fit between the GAMYE and SLOPE and GAMYE and GAM for Barn Swallow and 9 other selected species. Species short forms are WOTH = Wood Thrush (*Hylocichla mustelina*), RTHU = Ruby-throated Hummingbird (*Archilochus colubris*), PISI = Pine Siskin (*Spinus pinus*), Cooper’s Hawk (*Accipiter cooperii*), CHSW = Chimney Swift (*Chaetura pelagica*), CCLO = Chestnut-collared Longspur (*Calcarius ornatus*), CAWR = Carolina Wren (*Thryothorus ludovicianus*), CAWA = Canada Warbler (*Cardellina canadensis*), MAKE = American Kestrel (*Falco sparverius*).

**Figure 5.**
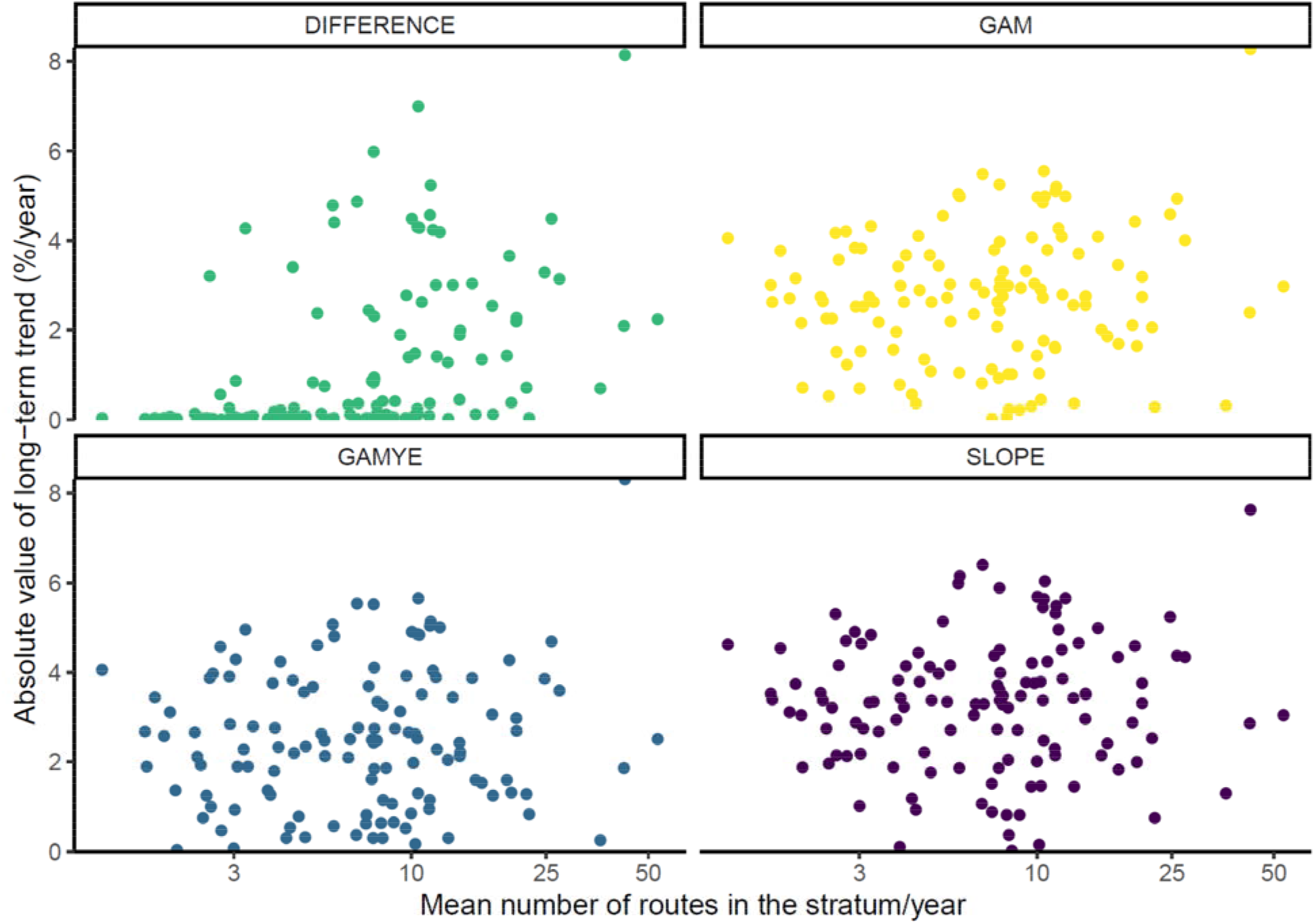
Relationship between the absolute value of estimated long-term trends (1966-2018) and the amount of data in each stratum, from the four models compared here for Cooper’s Hawk, a species with relatively sparse data in each individual stratum. More of the trends estimated with the DIFFERENCE model are close to zero, suggesting a stable population, and particularly where there are relatively few routes contributing data in each year. This relationship is not evident for the same data modeled with one of the three models that are able to share some information among strata on population change (GAM, GAMYE, and SLOPE).

For most species here, the GAMs or the DIFFERENCE model generally were preferred over the SLOPE model (Figure 4). For the two species with population trajectories that are known to include strong year-effects (Carolina Wren and Pine Siskin), the GAM model that does not include year-effects performed poorly (Figure 4). For Carolina Wren, the DIFFERENCE model was preferred clearly over the GAMYE (Figure 4), and yet the predicted trajectories from the two models are extremely similar (Figure 1). By contrast, for Pine Siskin the DIFFERENCE and GAMYE were very similar in their predictive accuracy (Figure 4) and yet the predicted trajectories are noticeably different in the first 10-years of the survey (Supplementary Materials Figure S1). For Cooper’s Hawk, the GAMYE model was generally preferred over the DIFFERENCE model, although there was some overlap with zero (Figure 4), but in this case, the predicted trajectories are very different. The DIFFERENCE trajectory for Cooper’s Hawk suggests much less change in the species’ population over time than the GAM or GAMYE (Figure 1).

Cooper’s Hawk provides an example of a species with very sparse data, for which the sharing of information in space may be particularly relevant. In a single stratum, the model has relatively few data with which to estimate changes in populations through time. For example, the mean counts for the species indicate that on average one bird was observed for every 40 BBS-routes run in the 1970s, and since the species population has increased it still requires more than 10 routes to observe a single bird. For this species, the models that share information among strata on population change (GAM, GAMYE, and SLOPE), suggest greater change in populations than the DIFFERENCE. For these models, where the stratum-level population change parameters are able to share information across the species’ range, the absolute change in the population does not depend on the sample size in the region. In addition, for each of these models, there is still large variability in the trends estimated for data-sparse regions, demonstrating that while the estimates benefit from the sharing of information among strata, the local trends are still influenced by the local data. By contrast, there is a strong relationship between the magnitude of the trend and the number of routes contributing data to the analysis for the DIFFERENCE model (Figure 5). In strata with fewer than 10 routes contributing data, the DIFFERENCE trends are almost all very close to zero. In these relatively data-sparse strata, the DIFFERENCE model has very little information available to estimate population change, and so the prior is more relevant and the population changes are shrunk towards zero. By contrast, the other models can integrate data from the local stratum with information on changes in the species’ population across the rest of the its range.

The decomposed trajectories from the GAMYE allow us to calculate trends from the smooth but also plot trajectories that show the annual fluctuations. For example, the smooth trajectory for the Carolina Wren captures the general patterns of increases and decreases well, while the full trajectory also shows the sharp population crash associated with the extreme winter in 1976 (Figure 6). Calculating trends from the smooth component generates short-term estimates that vary less from year to year for species with relatively strong annual fluctuations (Figure 7). For example, Figure 8 shows the series of short-term (10-year) trend estimates for Wood Thrush in Canada, from the smooth component of the GAMYE, the GAMYE including the year-effects, the DIFFERENCE model, and the SLOPE model used since 2011. In this example, the 10-year trend estimate from the GAMYE with the year-effects and the SLOPE model both cross the IUCN trend threshold criterion for Threatened (IUCN 2019) at least once in the last 12 years, including 2011, when the species’ status was assessed in Canada (COSEWIC 2012). By contrast, a trend calculated from the decomposed GAMYE model using only the smooth component (GAMYE – Smooth Only in Figure 8) fluctuates much less between years.

**Figure 6.**
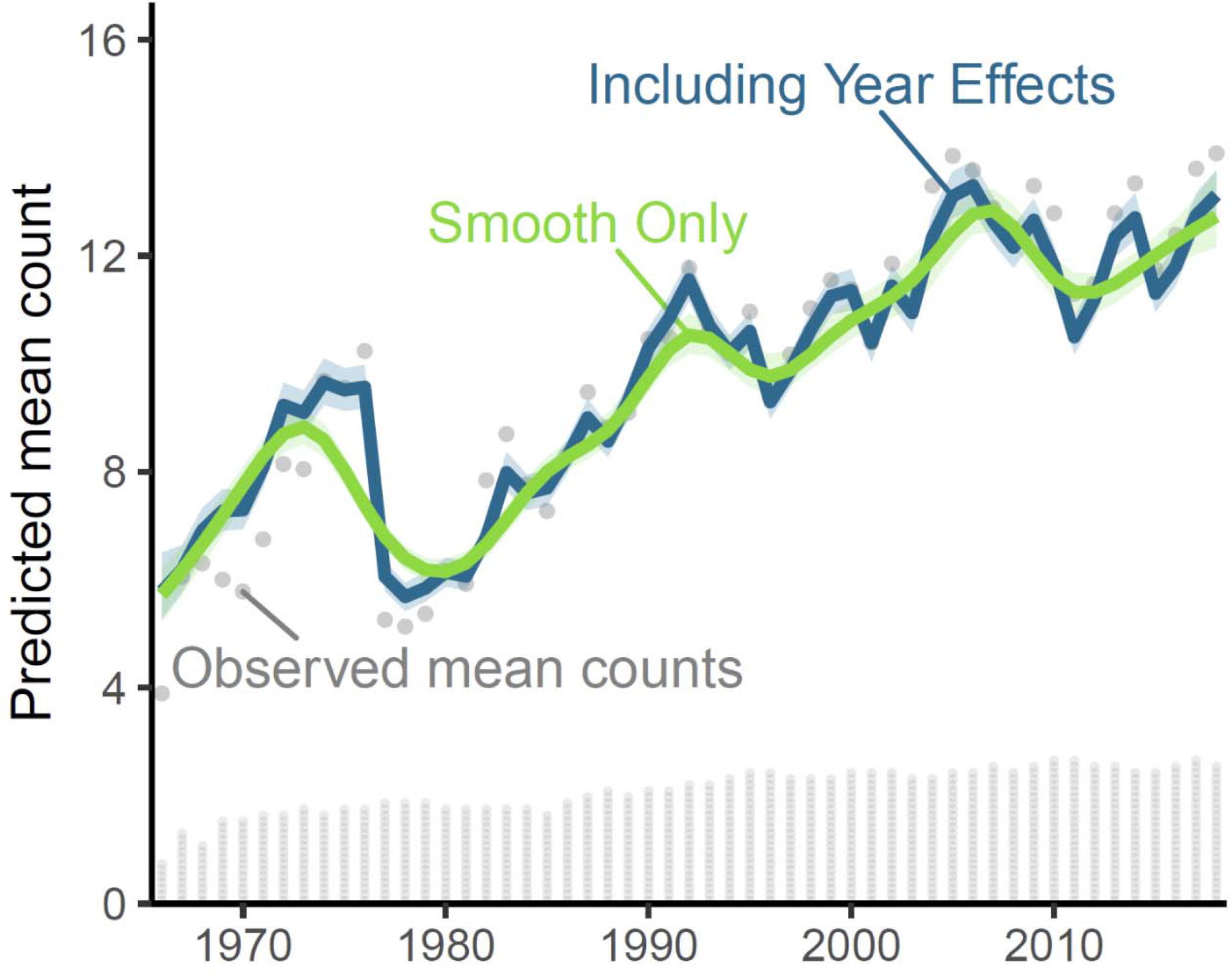
Decomposition of the survey-wide population trajectory for Carolina Wren (*Thryothorus ludovicianus*), from the GAMYE, showing the full trajectory (“Including Year Effects”, *N*_*s,t*_) and the isolated smooth component (“Smooth Only”, *Ng*_*s,t*_), which can be used to estimate population trends that are less sensitive to the particular year in which they are estimated. The stacked dots along the x-axis indicate the approximate number of BBS counts used in the model; each dot represents 50 counts.

**Figure 7.**
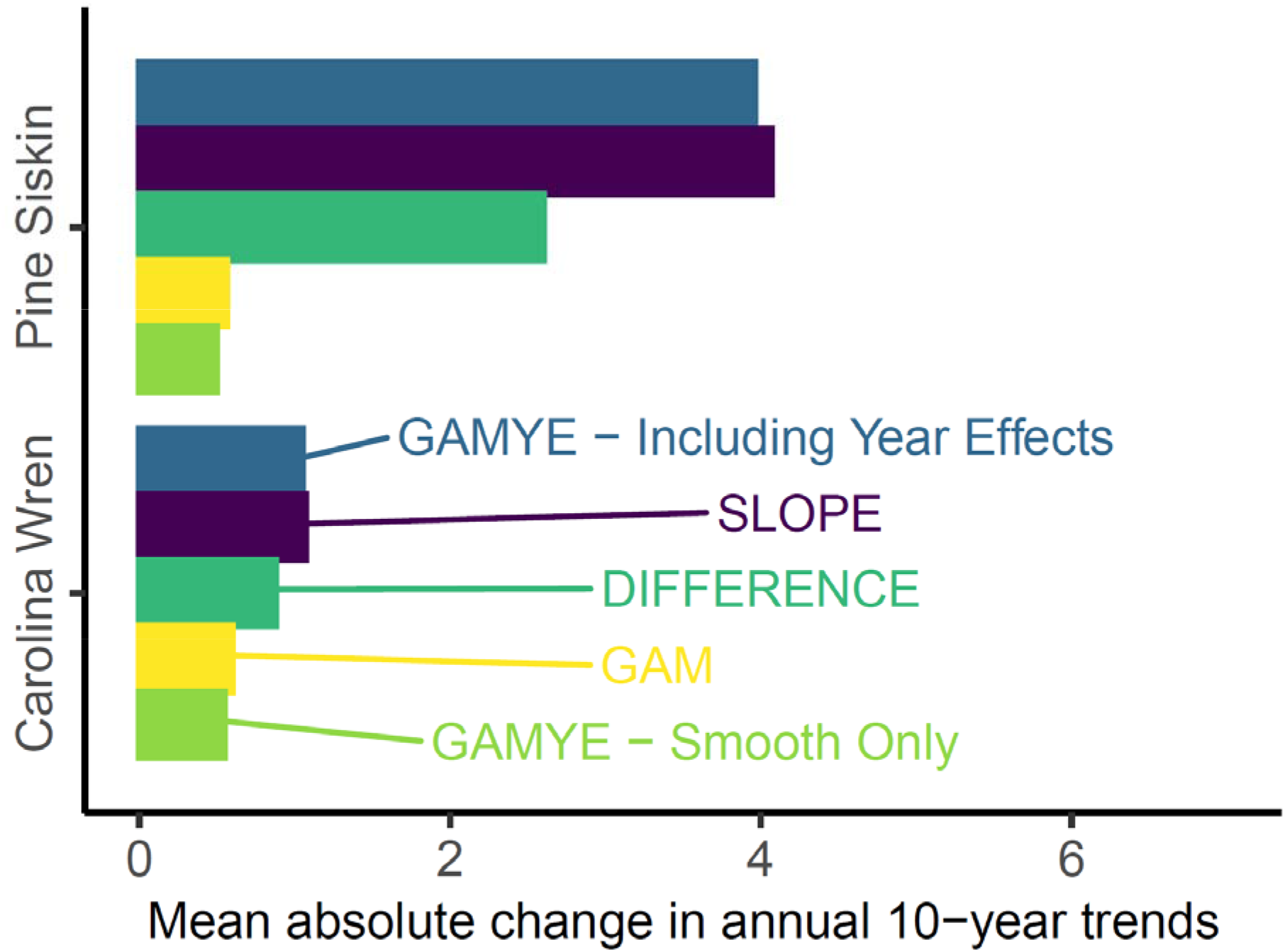
Inter annual variability of 10-year trend estimates for two species with large annual fluctuations (%/year). Trends from the GAM, which does not model annual fluctuations, and from the GAMYE using only the smooth component, which removes the effect of the annual fluctuations, are less variable between subsequent years (i.e., more stable) than trends from the GAMYE including the year-effects or the other two models that include the annual fluctuations.

**Figure 8.**
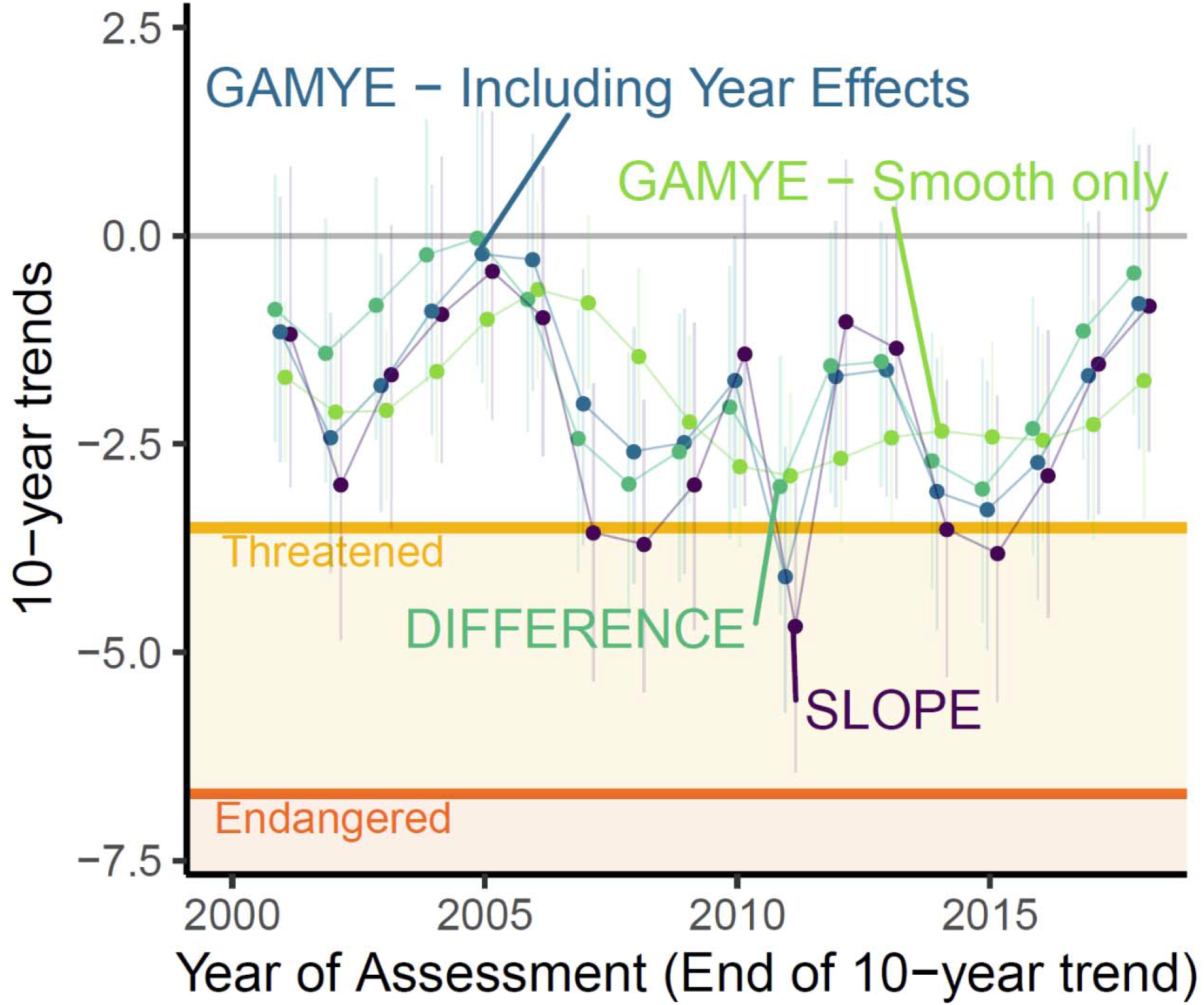
Sequential, short-term trend estimates for Wood Thrush (*Hylocichla mustelina*) in Canada from three alternative modeling approaches, and their comparison to the IUCN trend criteria for “Threatened” (in orange) and “Endangered” (in Red). Trends estimated from the decomposed trajectory of the GAMYE that include only the smooth component (in blue) are more stable between sequential years than trends from the other models that include annual fluctuations.

### Cross-validation varies in time and space

The preferred model from the pairwise predictive fit comparisons varied in time and space (Figures 4, 9, 10 and Supplemental Material Figure S6). The contrast between GAMYE and DIFFERENCE for Barn Swallow provide a useful example: Depending on the year or the region of the continent, either the GAMYE or the DIFFERENCE model was the preferred model, but overall, and in almost all regions and years, the 95% CI of the mean difference in fit between GAMYE and DIFFERENCE overlapped 0 (Figures 4, 9 and 10). For Barn Swallow, the GAMYE model has generally higher predictive fit during the first 5 years of the time-series, but then the DIFFERENCE model is preferred between approximately 1975 and 1983. The geographic variation in predictive fit is similarly complex. In the Eastern parts of the Barn Swallow’s range, the GAMYE model generally outperforms the DIFFERENCE model, whereas the reverse is generally true in the remainder of the species’ range (Figure 10). Although the mapped colours only represent the point-estimates, they suggest an interesting spatial pattern in the predictive fit of these two models for this species. Many of species considered here show similarly complex temporal and spatial patterns in the preferred model based on predictive fit (Supplemental Material Figures S6).

**Figure 9.**
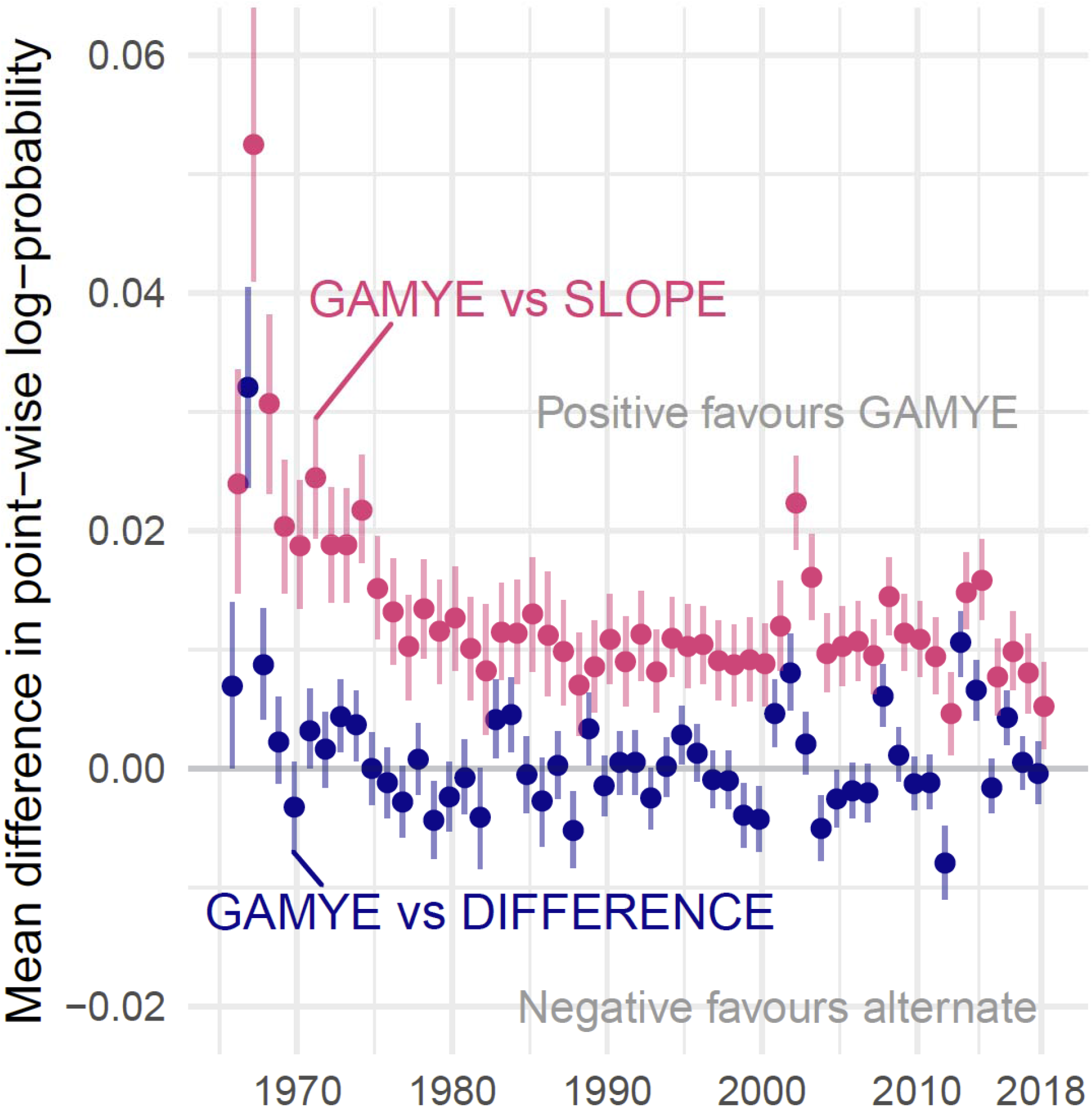
Annual differences in predictive fit between the GAMYE and SLOPE (blue) and the GAMYE and DIFFERENCE model (red) for Barn Swallow.

**Figure 10.**
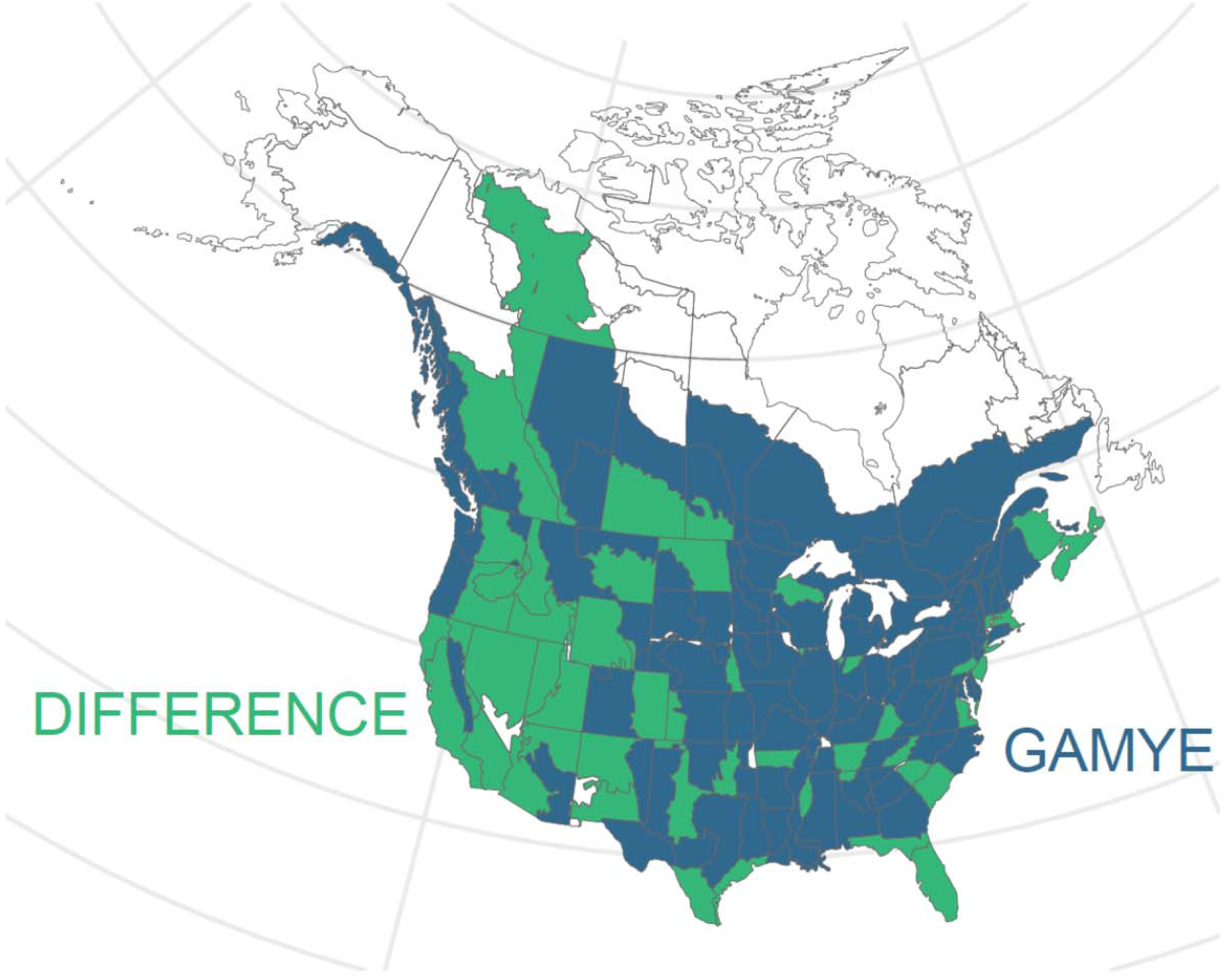
Geographic distribution of the preferred model for Barn Swallow, according to the point-estimate of the mean difference in predictive fit between GAMYE and DIFFERENCE. The GAMYE is generally preferred in the Eastern part of the species’ range, but the DIFFERENCE is preferred in many regions in the Western part of the species’ range. Note: in most regions, the differences in predictive fit were variable and neither model was clearly preferred (i.e., the 95% CI of the mean difference included 0).

## DISCUSSION

Using Bayesian hierarchical semi-parametric GAM smooths to model time series of population abundance with the North American Breeding Bird Survey generates useful estimates of population trajectories and trends and has better or comparable out of sample predictive accuracy, in comparison to the SLOPE or DIFFERENCE model. The flexibility of the GAM smoothing structure to model long- and medium-term temporal patterns, and the optional addition of random year-effects to model annual fluctuations, allow it to model a wide range of temporal patterns within a single base-model (Fewster et al. 2000, Wood 2017). We fit the smoothing parameters as random effects, to share information across geographic strata within a species’ range, and to improve the estimates of population trajectories for data-sparse regions (Pedersen et al. 2018). For almost all species included here, the two GAM-based models clearly out-performed the standard model (SLOPE) used for the CWS and USGS analyses since 2011 (Sauer and Link 2011, Smith et al. 2014), and showed similar out of sample predictive accuracy as a first-difference, random-walk trajectory model (Link et al. 2020). On a practical note, the GAM-based models required approximately 40% more time than the SLOPE or DIFFERENCE model to generate a similar number of posterior samples, but given the 53 years of effort to collect the data, we suggest this is a small price to pay for useful status and trend estimates.

The decomposition of the estimated population trajectory into the smooth and year-effect components is a feature of the GAMYE that is particularly useful for conservation applications. It allows the user to estimate and visualize separate trends and trajectories that include or exclude the annual fluctuations (Knape 2016). This allows the estimates to suit a range of conservation and management applications that rely on visualizing and estimating multiple aspects of population change. For example, the smoothed population trajectories capture the medium- and long-term changes in populations that are most relevant to broad-scale, multi-species assessments like the “State of the Birds” reports (NABCI-Canada 2019) where the annual fluctuations of a given species are effectively noise against the signal of community level change over the past 50 years (e.g., Rosenberg et al. 2019). Similarly, estimates of population trends (interval-specific, rates of annual change) derived from the smooth component are responsive to medium-term changes and so can be used to identify change points in trends such as the recovery of Species at Risk (Environment Climate Change Canada 2016).

Trends derived from the smooth component of the GAMYE are responsive to medium-term changes, but also much less likely to fluctuate from year to year and therefore more reliable for use in species at risk status assessments (James et al. 1996). In many status assessments, such as those by IUCN and COSEWIC, population declines beyond a particular threshold rate can trigger large investments of resources related to policy and conservation actions. For example, in both the IUCN red-listing and Canada’s federal species at risk assessments (IUCN 2019) estimated population declines greater than 30% over three generations is one criteria that results in a “Threatened” designation. If the estimated rate of population decline fluctuates from year to year, and is therefore strongly dependent on the particular year in which a species is assessed, there is an increased risk of inaccurate assessments. These inaccuracies could result in failures to protect species or inefficient investments of conservation resources. Of course, the full assessments of species’ status are sophisticated processes that consider more than just a single trend estimate. However, the example of Wood Thrush trends for Canada in Figure 8 shows that trends used to assess the species were below the threshold for “Threatened” status in 2011, but not in either year adjacent to 2011. The smooth-only trend never dips below the threshold (Figure 8) and raises the question of whether Wood Thrush would have been assessed as Threatened in Canada if the relevant trend had not been estimated in 2011, or had been estimated using a different model (COSEWIC 2012).

Alternative metrics of population trends that remove the annual fluctuations have been used with for the BBS, such as LOESS smooths (James et al. 1996) or slopes of log-linear regression lines calculated as part of the underlying model (Link and Sauer 1994) or as derived parameters from series of estimated annual indices (Sauer and Link 2011). Trend estimates that remove the effect of the annual fluctuations are generally a very common approach to summarizing average rates of change in other monitoring programs (e.g., Fewster et al. 2000 for UK breeding birds, Bogaart, et al. 2020, for European breeding birds). Many alternative definitions of trend could be calculated using the annual indices derived from any one of the models compared here. However for the last decade, both national agencies have supplied authoritative trend estimates based on end-point comparisons of annual indices, which include the annual fluctuations (Sauer and Link 2011, Smith et al. 2015). Similarly, calculating alternative metrics of trend from the annual indices in a way that propagates uncertainty would be done best using information from the full posterior distribution of each annual index. Given that these full posterior distributions are challenging for users to manipulate and summarize, we suggest that providing the authoritative trends based on the smooth component from the GAMYE is a practical and simple solution. These smooth-based trends are responsive to cycles or changes in rates of population change (discussed in James et al. 1996 and Sauer and Link 2011) while they also limit the annual fluctuations that might otherwise undermine the utility and credibility of BBS-trends for species status assessments (see also Smith et al. 2015).

In some conservation or scientific uses of the BBS-based population trajectories, the annual fluctuations may be important components of the trajectory (e.g., winter-related mortality of Carolina Wrens), and in these situations both components from the GAMYE can be presented. This comprehensive estimate of a species’ population trajectory is likely the best approach for the official presentation of a time series. At a glance, managers, conservation professionals, and researchers can glean information about fluctuations that might relate to annual covariates such as precipitation, wintering ground conditions, or cone-crop cycles. The GAMYE structure allows an agency like the CWS to provide estimates in multiple versions (e.g., full trajectories and smoothed trajectories in the same presentation, such as Figure 6), drawn from a coherent model, to suit a wide range of conservation applications, and to produce them in an efficient way. For example, there are situations where the ability for a user to access a ready-made separation of the yearly fluctuations from the underlying smooth could be helpful in the initial formulation of an ecological hypothesis. In addition, for custom analyses (Edwards and Smith 2020) a researcher can modify the basic GAMYE model to include annual covariates on the yearly fluctuations (e.g., extreme weather during migration, or spruce cone mast-years) and other covariates on the smooth component (e.g., climate cycles).

### Predictive accuracy

Overall, the cross-validation comparisons generally support the GAMYE, GAM, or DIFFERENCE model over the SLOPE model for the species considered here, in agreement with Link et al. (2020). For Barn Swallow, the overall difference in predictive fit, and particularly the increasing predictive error of the SLOPE model in the earliest years, strongly suggests that in the period between the start of the BBS (1966) and approximately 1983 (Smith et al. 2015), Barn Swallow populations increased. All models agree, however, that since the mid-1980’s populations have decreased.

Using all data in our cross-validations allowed us to explore the spatial and temporal variation in fit, and to compare the fit across all data used in the model. We have not reported absolute values of predictive fit because estimates of fit from a random selection of BBS counts, or simple summaries of predictive fit from the full dataset, are biased by the strong spatial and temporal dependencies in the BBS data (Roberts et al. 2017). However, because our folds were identical across models, and we modeled the differences in fit with an additional hierarchical model that accounted for repeated measures among strata and years, we are reasonably confident that relative-fit assessments are unbiased within a species and among models. Alternative approaches, such as blocked cross-validation (Roberts et al. 2017) to assess the predictive success of models in time and space, and targeted cross-validation (Link et al. 2017) to explore the predictive success in relation to particular inferences (e.g., predictive accuracy in the end-point years used for short- and long-term trend assessments) are an area of ongoing research. The overall predictive fit assessments provided some guidance on model selection for the species here, but not in all cases. The SLOPE model compared poorly against most other models in the overall assessment, similar to Link et al. 2020. However, among the other three models, many of the overall comparisons failed to clearly support one model, even in cases where the predicted population trajectories suggested very different patterns of population change (e.g., Cooper’s Hawk). For a given species, the best model varied among years and strata. These temporal and spatial patterns in predictive fit complicate the selection among models, given the varied uses of the BBS status and trend estimates (Rosenberg et al. 2017).

In general, estimates of predictive accuracy are one aspect of a thoughtful model building and assessment process, but are insufficient on their own (Gelman et al. 2013 pg. 180, Burnham and Anderson 2002 pg. 16). This is particularly true if there is little or no clear difference in overall predictive accuracy, but important differences in model predictions. For example, the overall cross validation results do not clearly distinguish among the SLOPE, DIFFERENCE, and GAMYE for Cooper’s Hawk, and yet predictions are very different between the DIFFERENCE model and the others (Figures 1, 4, and 5). Interestingly, the cross-validation approach in Link et al. 2020 suggested that the DIFFERENCE model was preferred over the SLOPE for Cooper’s Hawk, but we did not find that here (Supplemental Material Figure S3). The important differences in trend estimates and the equivocal cross-validation results suggests further research is needed into the criteria for, and consequences of, model selection for BBS status and trend estimates. Model selection is also complicated when overall predictive accuracy appears to clearly support one model and yet the important parameters (trends and trajectories) are not noticeably different. For example, the overall cross validation results for Carolina Wren suggest the DIFFERENCE model is preferred over the GAMYE, and yet the trajectories are almost identical (Figures 1 and 4). Predictive accuracy is also complicated when robust predictions are required for years or regions with relatively few data against which predictions can be assessed (e.g., the earlier years of the BBS, or data-sparse strata that still have an important influence on the range-wide trend). Model selection is complicated, and predictive accuracy would never be the only criterion used to select a model for the BBS analyses. Limits to computational capacity and a desire to avoid a data-dredging all-possible-models approach ensure that some thoughtful process to select the candidate models is necessary.

We agree with Link et al. (2020) that we should not select models based on a particular pattern in the results. In fact, the necessary subjective process occurs before any quantitative analyses (Burnham and Anderson 2002), and relies on “careful thinking” to balance the objectives; the model; and the data (Chatfield 1995). The careful thinking required to select a BBS model or to interpret the BBS status and trend estimates, is to consider the consequences of the potential conflicts between the model structures (“constraints on the model parameters” sensu Chatfield 1995) and the objectives of the use of the modeled estimates. For example, one of the models that shares information on population change among strata is likely preferable to the DIFFERENCE model for species with relatively sparse data in any given stratum, because the prior of the DIFFERENCE model (stable-population) will be more influential when the data are sparse. This prior-dependency of the results may not be identified by lower predictive accuracy of the estimates, as we the results for Cooper’s Hawk demonstrate (Figures 5). Similarly, a user of estimates from the DIFFERENCE model should carefully consider the conservation-relevant consequences of the prior and model structure when assessing potential changes in the population trends of declining and relatively rare species. These species’ short-term rates of decline could appear to decrease, suggesting a stabilizing population, simply due to the increasing influence of the prior, if the species observations decline to a point where it is not observed in some years. In contrast, if a user wished to explicitly compare estimates of population change among political jurisdictions or ecological units, the sharing of information among those units in the GAM-based models here might be problematic. We suggest that the GAMYE’s strong cross-validation performance, its sharing of information across a species range, its decomposition of the population trajectory, and its broad utility that suits the most common uses of the BBS status and trends estimates, make it a particularly useful model for the sort of omnibus analyses conducted by the CWS and other agencies.

## ACKNOWLEDGEMENTS

We sincerely thank the thousands of U.S. and Canadian participants who annually perform and coordinate the North American Breeding Bird Survey. We also wish to acknowledge Courtney Amundson for sharing some code on similar models, and John Sauer and Bill Link for sharing code that helped with the cross-validations and for many spirited, collegial discussions that have informed this work. We also thank the many biologists within the Canadian Wildlife Service and other users of the BBS status and trend estimates whose insightful questions and suggestions motived much of this work, including Charles Francis, Marie-Anne Hudson, Veronica Aponte, Marcel Gahbauer, Pete Blancher, and Ken Rosenberg.

## Data Depository

R scripts to download the BBS data and to perform the analyses in this paper and are archived at www.github.com/AdamCSmithCWS/GAM_Paper_Script

## Ethics Statement

This research was conducted in compliance with the Environment and Climate Change Canada Values and Ethics Code.

## REFERENCES CITED

Bogaart, P., van der Loo, M., and Pannekoek, J. (2020). rtrim: Trends and Indices for Monitoring Data. R package version 2.1.1.

Burnham, K.P., and D.R. Anderson. (2002). Model selection and multimodel inference: a practical information-theoretic approach. Second edition. Springer-Verlag, New York, New York, USA.

Chatfield, C. (1995). Model uncertainty, data mining and statistical inference (with discussion).Journal of the Royal Statistical Society (London), Series A 158:419–466. doi:10.2307/2983440

COSEWIC. (2012). COSEWIC assessment and status report on the Wood Thrush Hylocichla mustelina in Canada. Committee on the Status of Endangered Wildlife in Canada.Ottawa.

Crainiceanu CM, Ruppert D, Wand MP (2005). Bayesian Analysis for Penalized Spline Regression Using WinBUGS. Journal of Statistical Software, 14 (14). doi:10.18637/jss.v014.i14

Duan, N. (1983). Smearing Estimate: A Nonparametric Retransformation Method. Journal of the American Statistical Association, Vol. 78, No. 383, pp. 605–610. doi:10.2307/2288126

Edwards, B.P.M. and A.C. Smith (2020). bbsBayes v2.1.0 (Version 2.1.0). Zenodo. doi:10.5281/zenodo.3727279

Environment Climate Change Canada (2016). Recovery Strategy for the Canada Warbler (Cardellina canadensis) in Canada. Species at Risk Act Recovery Strategy Series.

Environment Canada, Ottawa. vii + 56 pp. Available at: http://www.registrelep-sararegistry.gc.ca, accessed February 13, 2020.

Fewster, R. M., S. T. Buckland, G. M. Siriwardena, S. R. Baillie, and J. D. Wilson (2000). Analysis of population trends for farmland birds using generalized additive models. Ecology 81:1970–1984. doi: 10.2307/177286

Gelman, A. (2006). Prior distributions for variance parameters in hierarchical models. Bayesian Analysis. 1:515–533. doi:10.1214/06-BA117A

Gelman, A., J. Hwang, and A. Vehtari (2014). Understanding predictive information criteria for Bayesian models. Statistics and Computing 24:997–1016. doi:10.1007/s11222-013-9416-2

Gelman, A., J. B. Carlin, H. S. Stern, D. B. Dunson, A. Vehtari and D. B. Rubin. (2013).Bayesian Data Analysis, Chapman and Hall/CRC Boca Raton.

Hudson, M.-A. R., Francis, C. M., Campbell, K. J., Downes, C. M., Smith, A. C., & Pardieck, K.L. (2017). The role of the North American Breeding Bird Survey in conservation. The Condor, 119(3), 526–545. https://doi.org/10.1650/CONDOR-17-62.1

IUCN Standards and Petitions Committee. (2019). Guidelines for Using the IUCN Red List Categories and Criteria. Version 14. Prepared by the Standards and Petitions Committee. Downloadable from http://www.iucnredlist.org/documents/RedListGuidelines.pdf.

es, F. C., McCulloch, C. E., & Wiedenfeld, D. A. (1996). New Approaches to the Analysis of Population Trends in Land Birds. Ecology, 77(1), 13–27. https://doi.org/10.2307/2265650

Lange, K. L., Little, R. J. A, and J. M. G. Taylor (1989). Robust Statistical Modeling Using the t Distribution. Journal of the American Statistical Association, 84:408, 881-896, DOI:10.2307/2290063

Knape, J. (2016). Decomposing trends in Swedish bird populations using generalized additive mixed models. Journal of Applied Ecology, 53, 1852–1861. doi: 10.1111/1365-2664.12720

Kohavi, R. (1995). A study of cross-validation and bootstrap for accuracy estimation and model selection. Proceedings of the 14th International Joint Conference on Artificial Intelligence - Volume 2, 1137–1143.

Link, W. A., & Sauer, J. R. (2002). A Hierarchical Analysis of Population Change with Application to Cerulean Warblers. Ecology, 83(10), 2832–2840.

Link, W., & Sauer, J.R. (2007). Seasonal Components of Avian Population Change: Joint Analysis of Two Large-Scale Monitoring Programs. Ecology, 88(1), 49–55.

Link, W. A. and J. R. Sauer (2016). Bayesian cross-validation for model evaluation and selection, with application to the North American Breeding Bird Survey. Ecology 97:1746–1758. doi: 10.1890/15-1286.1

Link, W.A., J.R. Sauer, and D.K. Niven. (2017). Model selection for the North American Breeding Bird Survey: A comparison of methods. Condor 119(3):546–556. doi: 10.1650/CONDOR-17-1.1

Link, W. A., J.R. Sauer, and D.K. Niven. (2020). Model selection for the North American Breeding Bird Survey Ecological Applications. https://doi.org/10.1002/eap.2137.

Microsoft and Weston, S. (2019). foreach: Provides Foreach Looping Construct. R package version 1.4.7. https://CRAN.R-project.org/package=foreach

North American Bird Conservation Initiative Canada. (2019). The State of Canada’s Birds, 2019.Environment and Climate Change Canada, Ottawa, Canada. 12 pages. www.stateofcanadasbirds.org

Partners in Flight. (2019).Avian Conservation Assessment Database, version 2019. Available at http://pif.birdconservancy.org/ACAD

Plummer, Martyn. (2003). JAGS: A program for analysis of Bayesian graphical models using Gibbs sampling.

Pedersen EJ, Miller DL, Simpson GL, Ross N. (2019). Hierarchical generalized additive models in ecology: an introduction with mgcv. PeerJ 7:e6876. doi: 10.7717/peerj.6876

R Core Team (2019). R: A language and environment for statistical computing. R Foundation for Statistical Computing, Vienna, Austria. URL https://www.R-project.org/.

Roberts, D. R., V. Bahn, S. Ciuti, M. S. Boyce, J. Elith, G. Guillera-Arroita, S. Hauenstein, J. J. Lahoz-Monfort, B. Schroder, W. Thuiller, D. I. Warton, B. A. Wintle, F. Hartig, and C. F. Dormann. (2017). Cross-validation strategies for data with temporal, spatial, hierarchical or phylogenetic structure. - Ecography doi: 10.1111/ecog.02881.

Robbins, C. S., D. Bystrak, and P. H. Geissler (1986). The Breeding Bird Survey: Its first fifteen years, 1965–1979. U.S. Fish and Wildlife Service Resource Publication 157.

Rosenberg, K. V., P. J. Blancher, J. C. Stanton, and A. O. Panjabi (2017). Use of North American Breeding Bird Survey Data in avian conservation assessments. The Condor: Ornithological Applications 119:594–606. doi: 10.1650/CONDOR-17-57.1

Rosenberg, K. V., Dokter, A. M., Blancher, P. J., Sauer, J. R., Smith, A. C., Smith, P. A., Stanton, J.C., Panjabi, A., Helft, L., Parr, M., Marra, P.P. (2019). Decline of the North American avifauna. Science 366, 120–124. doi: 10.1126/science.aaw1313

Sauer, J. R., and W. A. Link. (2011). Analysis of the North American Breeding Bird Survey using hierarchical models. The Auk 128:87–98. doi: 10.1525/auk.2010.09220

Sauer, J.R., J.E. Hines, J.E. Fallon, K.L. Pardieck, D.J. Ziolkowski Jr., and W.A. Link. (2014). The North American Breeding Bird Survey, Results and Analysis 1966 – 2013. Version 01.30.2015 USGS Patuxent Wildlife Research Center, Laurel, MD.

Sauer, J. R., Pardieck, K. L., Ziolkowski, D. J., Smith, A. C., Hudson, M.-A. R., Rodriguez, V., Berlanga, H., Niven, D. K., & Link, W. A. (2017). The first 50 years of the North American Breeding Bird Survey. The Condor, 119(3), 576–593.

Smith A.C., M.-A.R. Hudson, C. Downes, and C.M. Francis (2014). Estimating breeding bird survey trends and annual indices for Canada: how do the new hierarchical Bayesian estimates differ from previous estimates. Canadian Field-Naturalist 128:119–134. doi: 10.22621/cfn.v128i2.1565

Smith, A. C., M.-A.R. Hudson, C. Downes, and C.M. Francis (2015). Change points in the population trends of aerial-insectivorous birds in North America: synchronized in time across species and regions. PLoS One 10:e0130768. doi: 10.1371/journal.pone.0130768

Smith, A.C., Hudson, M-A.R. Aponte, V., and Francis, C.M. (2019). North American Breeding Bird Survey - Canadian Trends Website, Data-version 2017. Environment and Climate Change Canada, Gatineau, Quebec, K1A 0H3

Vehtari, A., Gelman, A., and Gabry, J. (2017). Practical Bayesian model evaluation using leave-one-out cross-validation and WAIC. Statistics and Computing. 27(5), 1413--1432. doi:10.1007/s11222-016-9696-4.

Wickham, H. (2016). ggplot2: Elegant Graphics for Data Analysis. Springer-Verlag New York

Wilson, S., Smith, A. C., & Naujokaitis□Lewis, I. (2018). Opposing responses to drought shape spatial population dynamics of declining grassland birds. Diversity and Distributions, 24, 1687–1698. doi: 10.1111/ddi.12811

Wood, S. N. (2017). Generalized additive models: an introduction with R; 2nd ed. CRC Press. Portland, OR, 2017

Zhang, Y., & Yang, Y. (2015). Cross-validation for selecting a model selection procedure. Journal of Econometrics, 187(1), 95–112. https://doi.org/10.1016/j.jeconom.2015.02.006

