## Supplemental Appendix for "North American Breeding Bird Survey status and trend estimates to inform a wide-range of conservation needs, using a flexible Bayesian hierarchical generalized additive model"

Annual indices for all the models here were calculated following Smith et al. (2019) which is conceptually similar to the approach described in Sauer and Link (2011) and Smith et al. (2015), with one small variation. Here, instead of using a retransformation factor that assumes a log-normal distribution of observer-route effects to re-scale the annual indices (Sauer and Link 2011, Smith et al. 2015), we generated count scale predictions for every observer-route in a given stratum and averaged across the collection of predictions. More precisely, in the standard approach described in Sauer and Link (2011), the annual indices in a given year $t$ and stratum $s$ are calculated as

$$N_{s,t}^{'}=z_{s}*e^{A_{s,t}+0.5*\sigma_{\omega}^{2}+0.5*\sigma_{\varepsilon}^{2}}$$

where each ${N'}_{s,t}$ are exponentiated sums of the relevant components of the model ($A_{s,t}$), multiplied by a correction factor for the proportion of routes in the stratum on which the species has been observed ($z_{s}$, routes where the species has never been observed are dropped from the analysis). These components include the stratum-level intercepts and all of the parameters that estimate the time-series (i.e., the slopes, year-effects, and GAM smooths), plus two variance components that account for the asymmetric retransformation from the log-scale parameters to the count-scale annual indices ($0.5*\sigma_{\omega}^{2}+0.5*\sigma_{\varepsilon}^{2}$). The variance component associated with the observer-route effects ($\sigma_{\omega}^{2}$) is problematic, because it assumes that a global estimate of variance among observers and routes represents the true observer-route variance within each stratum equally well, and it assumes that the distribution of the estimated observer-route effects is approximately normal (Duan 1983). For many species, one or both of these two assumptions are not well supported, and as a result, annual indices for some species and regions are over-estimated (Smith et al. 2015).

We therefore calculated the annual indices $N_{s,t}$ as

$$N_{s,t}=z_{s}*\frac{\sum_{j\in J_{s}} e^{A_{s,t}+\omega_{j}+0.5*\sigma_{\varepsilon}^{2}}}{n_{Js}}$$

so that instead of relying on the half-variance, log-normal re-scaling factor ($0.5*\sigma_{\omega}^{2}$), we averaged count-scale predictions across all of the $n_{Js}$ observer-routes $j$ in the set of observer-route combinations in stratum $s$ ($J_{s}$). The conceptual difference is that $N_{s,t}$ values represent mean expected counts from among the existing collection of observer routes in a given stratum (i.e., all observer-route combinations from every year). In contrast, the $N_{s,t}^{'}$ values (i.e., the standard approach) represent the mean expected count from a hypothetical new observer-route combination. However, because the variance of the observer-route effects ($\sigma_{\omega}^{2}$) is not specific to the stratum, the hypothetical new observer-route is not necessarily a route from within the relevant stratum. In effect, the observer-route effects account for much of the variation in abundance of a species among different strata, because the routes are nested within strata; and therefore, the observer-route effects are not exchangeable. The practical effect of this difference is that the $N_{s,t}$ annual indices calculated here more closely reflect the observed average counts, across all years, on BBS routes in a given stratum. This difference has no effect on local trend estimates or the pattern of estimated population trajectories. Therefore, the stratum’s contribution to the overall trajectory and trend estimate is unbiased with respect to the observed relative abundance of the species in that stratum. In the R-package bbsBayes, both versions of $N$ are available for all of the models, but the $N_{s,t}$ approach is the default. To our knowledge, the variance component that relates to the count-level extra-Poisson variance ($0.5*\sigma_{\varepsilon}^{2}$) better meets the necessary assumptions, although the specific re-scaling factor used to reflect the t-distribution of the error is an area of ongoing research (Link et al. 2020).
