## Supplemental Figures S1-S3 for "North American Breeding Bird Survey status and trend estimates to inform a wide-range of conservation needs, using a flexible Bayesian hierarchical generalized additive model"

### Supplemental Material Figures S1 through S4, Smith and Edwards 2020, Improved status and trend estimates from the North American Breeding Bird Survey using a Bayesian hierarchical generalized additive model

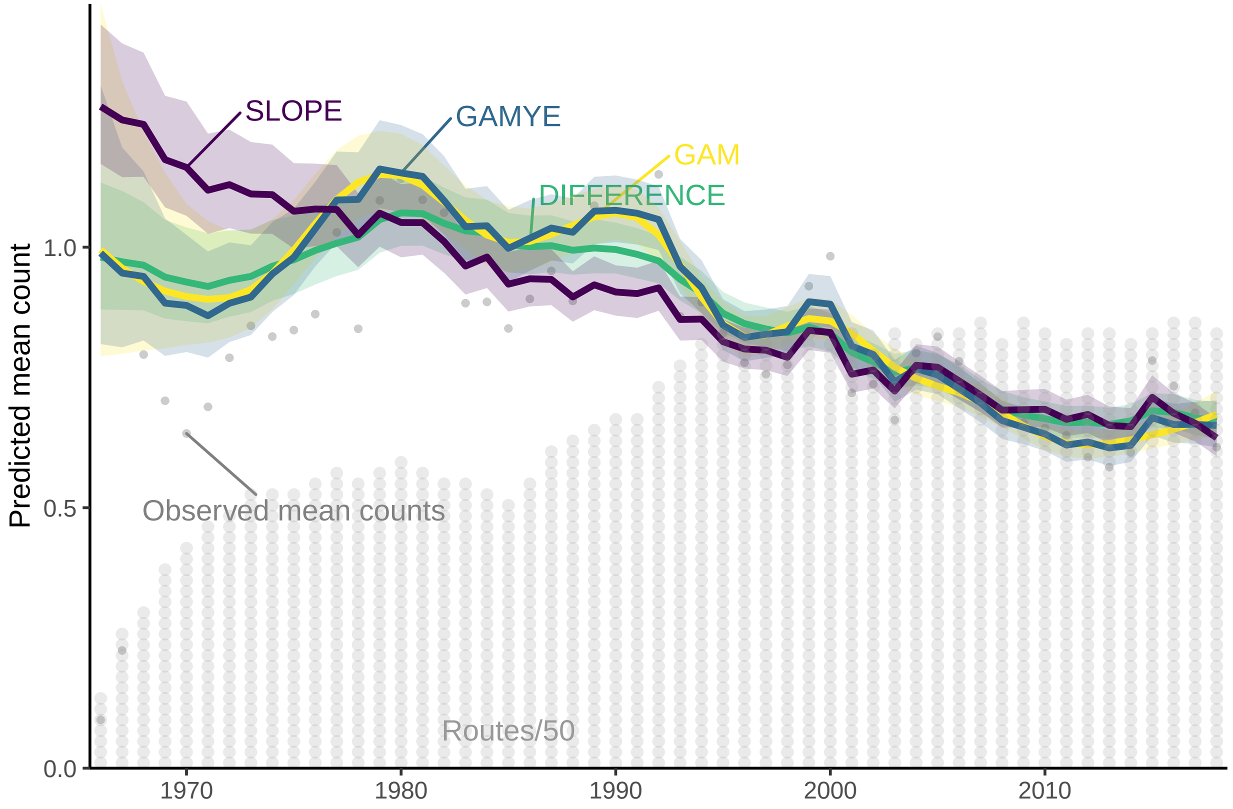


Figure 1: S1.A: Survey-wide population trajectories for American Kestrel estimated from the BBS using two models described here that include a GAM smoothing function to model change over time (GAM and GAMYE) the standard regression-based model used for BBS status and trend assessments since 2011 (SLOPE), and a first-difference time-series model (DIFFERENCE). The stacked dots along the x-axis indicate the approximate number of BBS counts used in the model in each year; each dot represents 50 counts

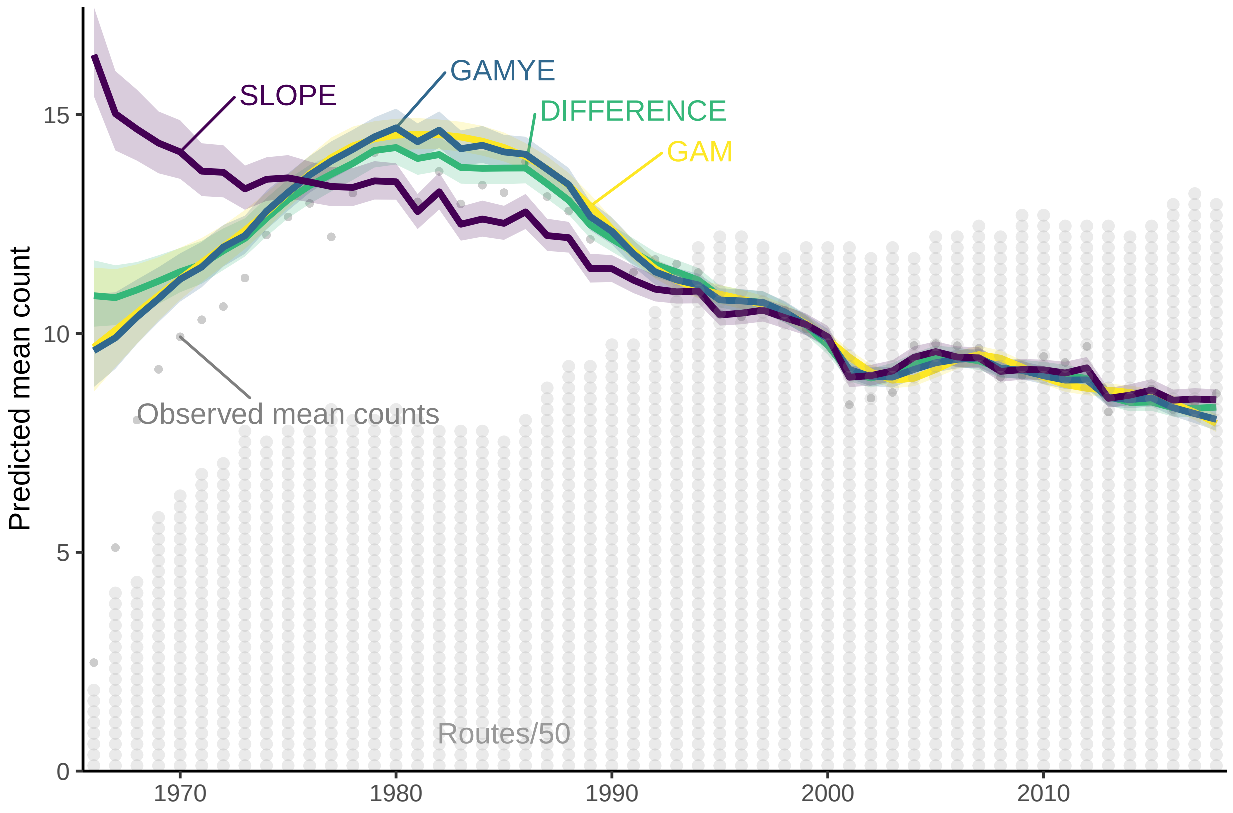


Figure 2: S1.B: Survey-wide population trajectories for Barn Swallow estimated from the BBS using two models described here that include a GAM smoothing function to model change over time (GAM and GAMYE) the standard regression-based model used for BBS status and trend assessments since 2011 (SLOPE), and a first-difference time-series model (DIFFERENCE). The stacked dots along the x-axis indicate the approximate number of BBS counts used in the model in each year; each dot represents 50 counts

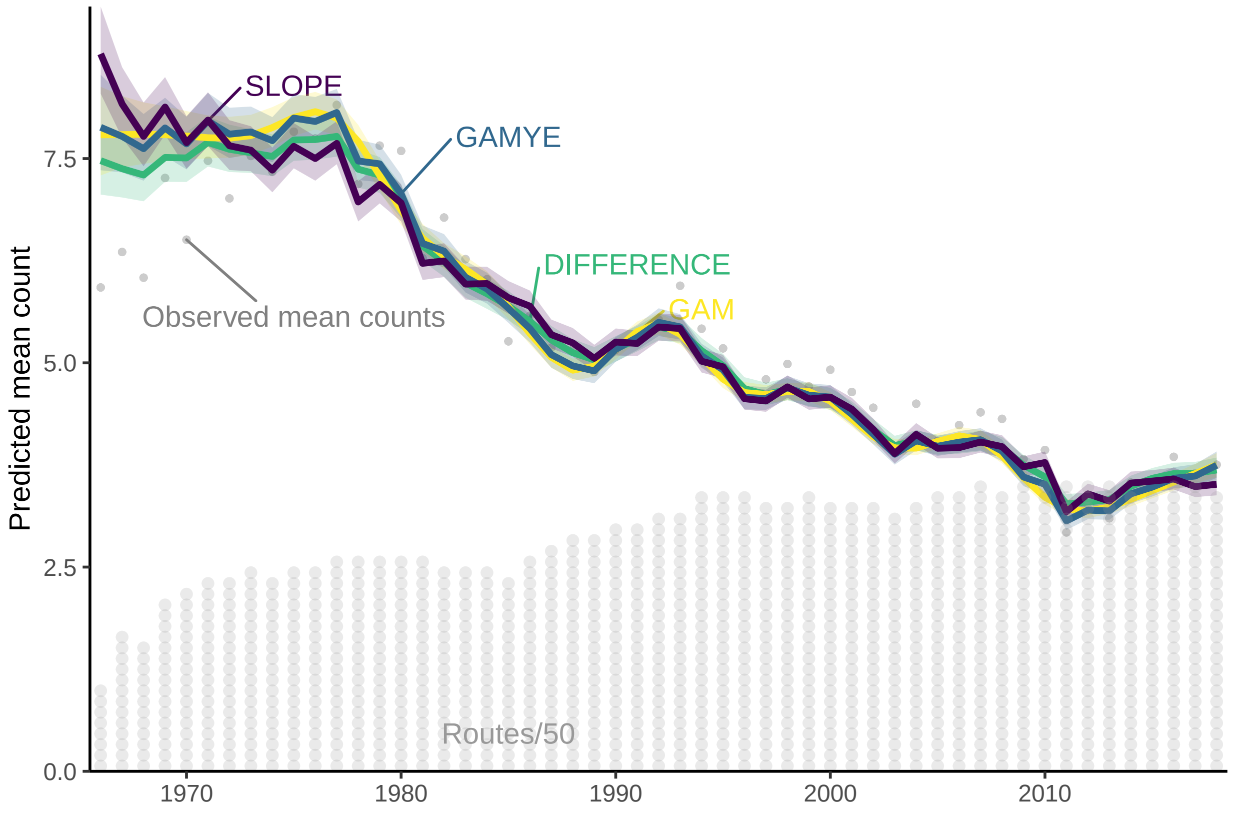


Figure 3: S1.C: Survey-wide population trajectories for Wood Thrush estimated from the BBS using two models described here that include a GAM smoothing function to model change over time (GAM and GAMYE) the standard regression-based model used for BBS status and trend assessments since 2011 (SLOPE), and a first-difference time-series model (DIFFERENCE). The stacked dots along the x-axis indicate the approximate number of BBS counts used in the model in each year; each dot represents 50 counts

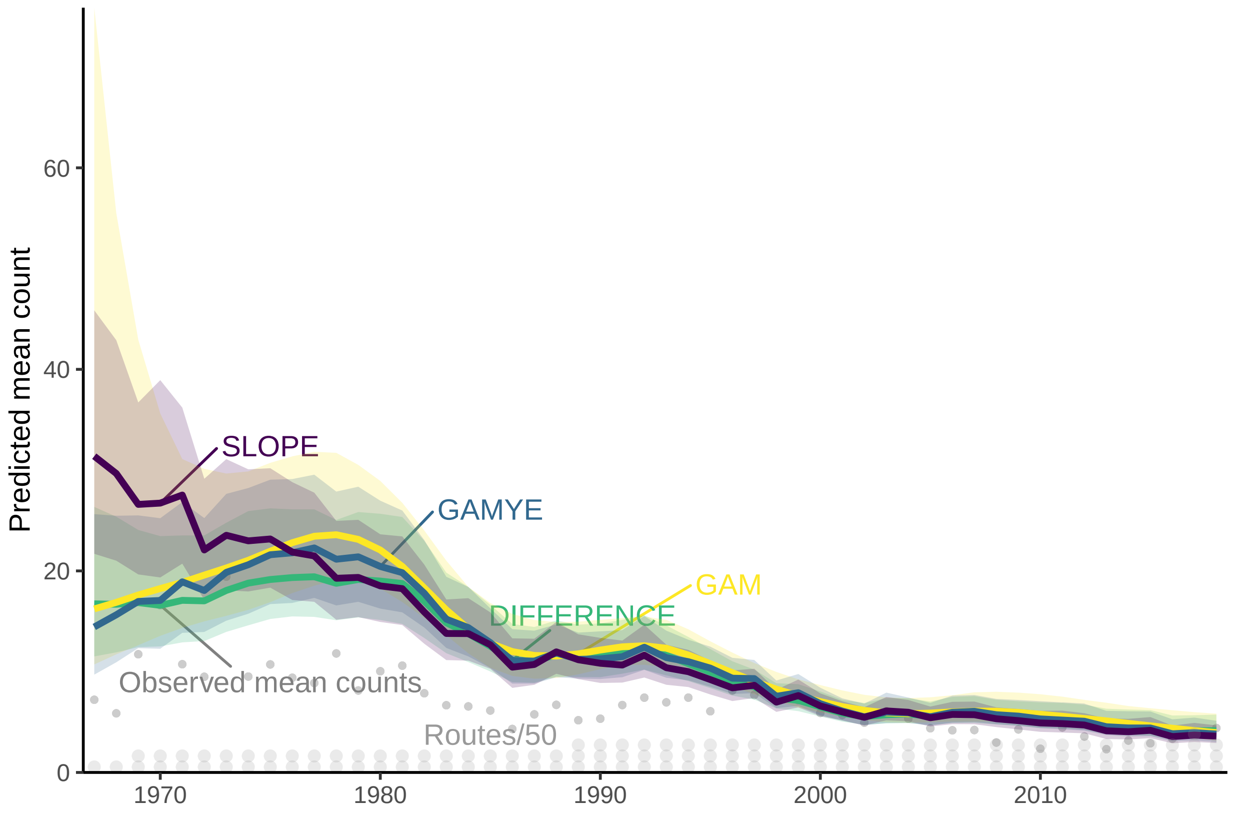


Figure 4: S1.D: Survey-wide population trajectories for Chestnut-collared Longspur estimated from the BBS using two models described here that include a GAM smoothing function to model change over time (GAM and GAMYE) the standard regression-based model used for BBS status and trend assessments since 2011 (SLOPE), and a first-difference time-series model (DIFFERENCE). The stacked dots along the x-axis indicate the approximate number of BBS counts used in the model in each year; each dot represents 50 counts

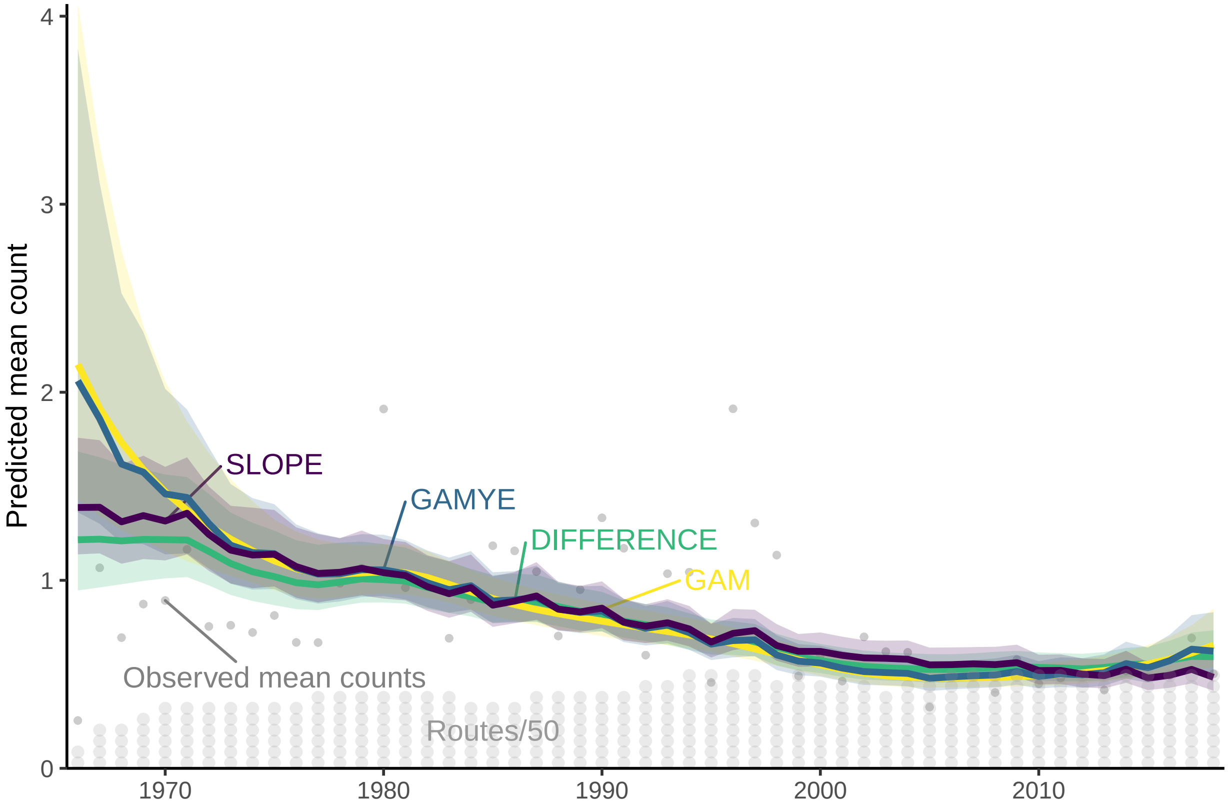


Figure 5: S1.E: Survey-wide population trajectories for Canada Warbler estimated from the BBS using two models described here that include a GAM smoothing function to model change over time (GAM and GAMYE) the standard regression-based model used for BBS status and trend assessments since 2011 (SLOPE), and a first-difference time-series model (DIFFERENCE). The stacked dots along the x-axis indicate the approximate number of BBS counts used in the model in each year; each dot represents 50 counts

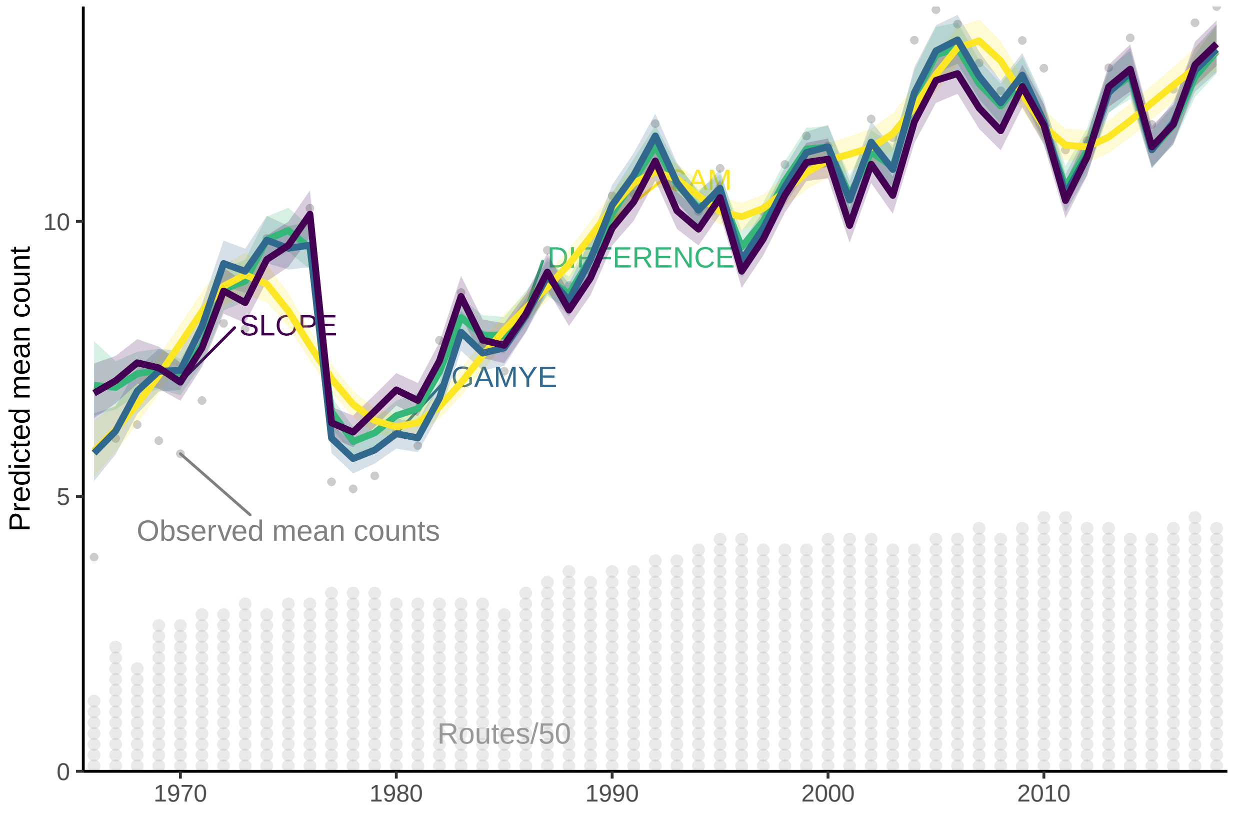


Figure 6: S1.F: Survey-wide population trajectories for Carolina Wren estimated from the BBS using two models described here that include a GAM smoothing function to model change over time (GAM and GAMYE) the standard regression-based model used for BBS status and trend assessments since 2011 (SLOPE), and a first-difference time-series model (DIFFERENCE). The stacked dots along the x-axis indicate the approximate number of BBS counts used in the model in each year; each dot represents 50 counts

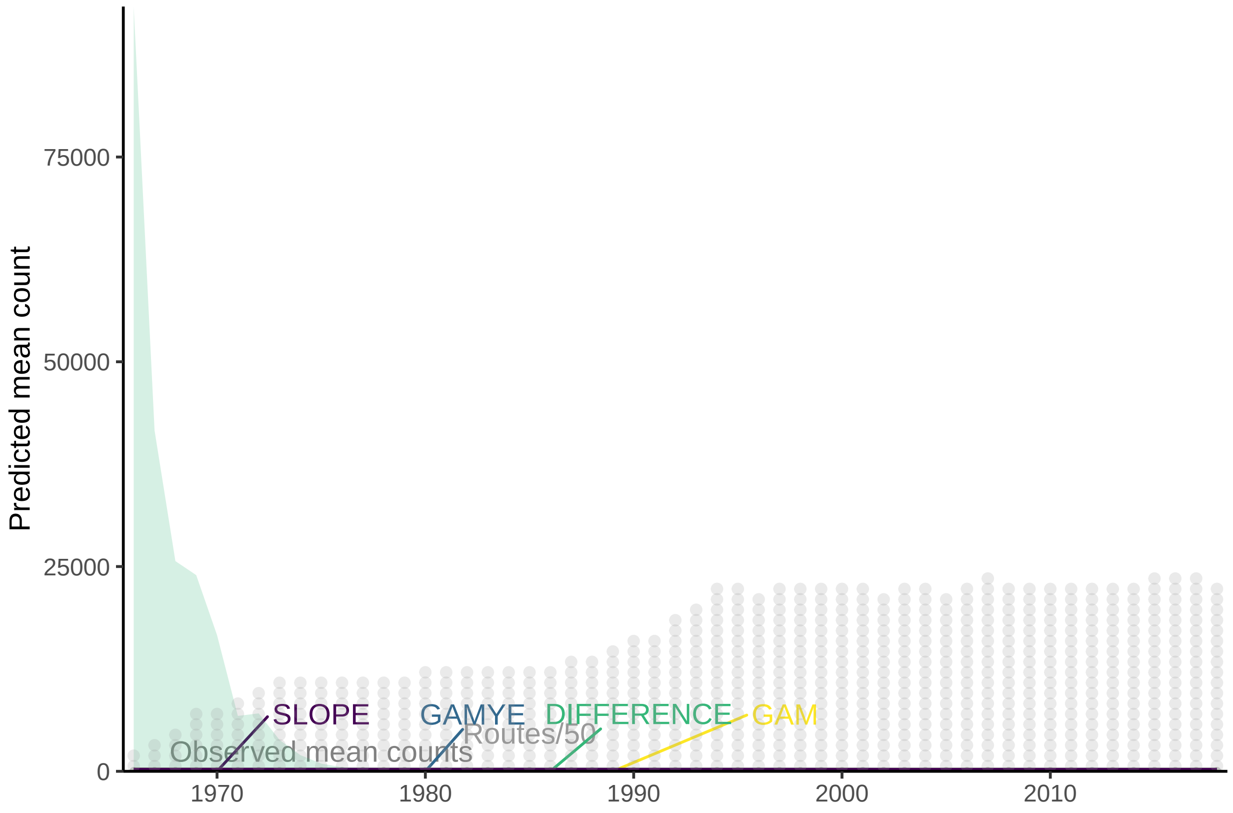


Figure 7: S1.G: Survey-wide population trajectories for Pine Siskin estimated from the BBS using two models described here that include a GAM smoothing function to model change over time (GAM and GAMYE) the standard regression-based model used for BBS status and trend assessments since 2011 (SLOPE), and a first-difference time-series model (DIFFERENCE). The stacked dots along the x-axis indicate the approximate number of BBS counts used in the model in each year; each dot represents 50 counts

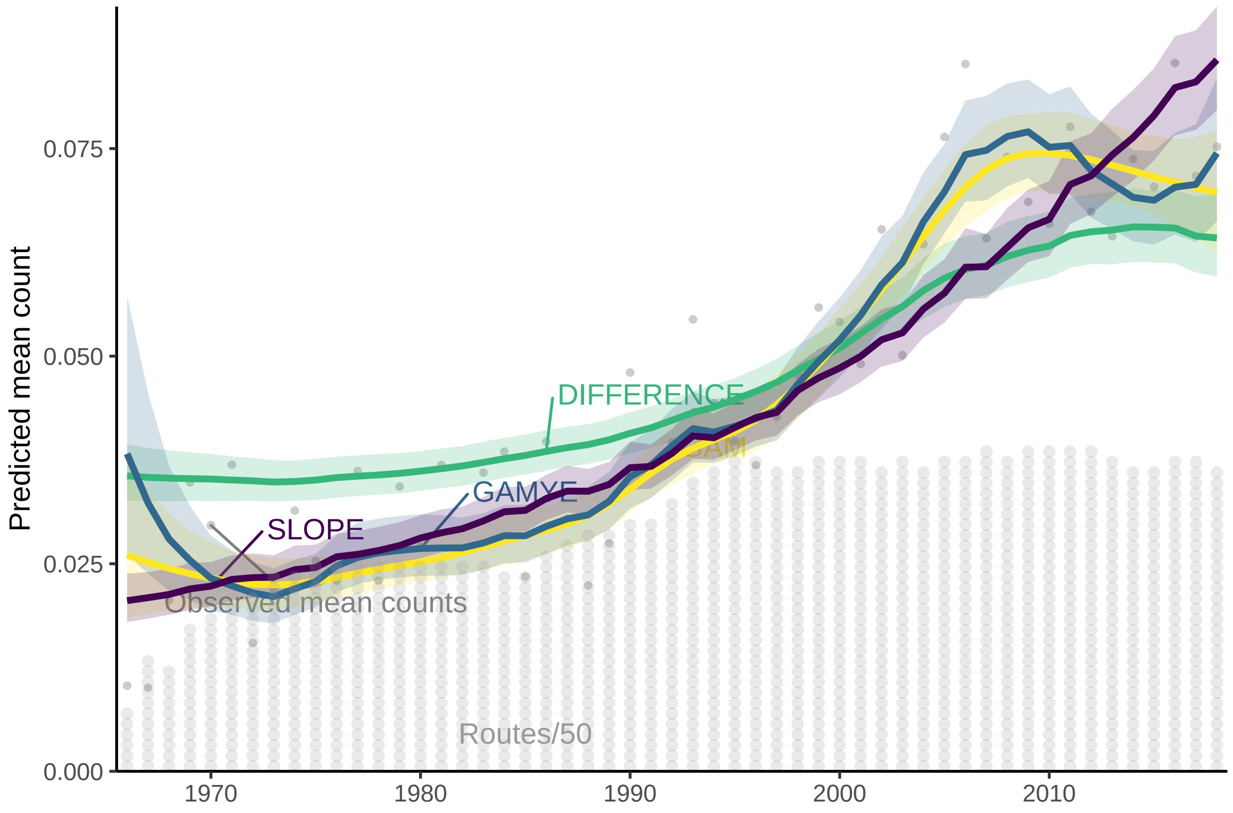


Figure 8: S1.H: Survey-wide population trajectories for Cooper's Hawk estimated from the BBS using two models described here that include a GAM smoothing function to model change over time (GAM and GAMYE) the standard regression-based model used for BBS status and trend assessments since 2011 (SLOPE), and a first-difference time-series model (DIFFERENCE). The stacked dots along the x-axis indicate the approximate number of BBS counts used in the model in each year; each dot represents 50 counts

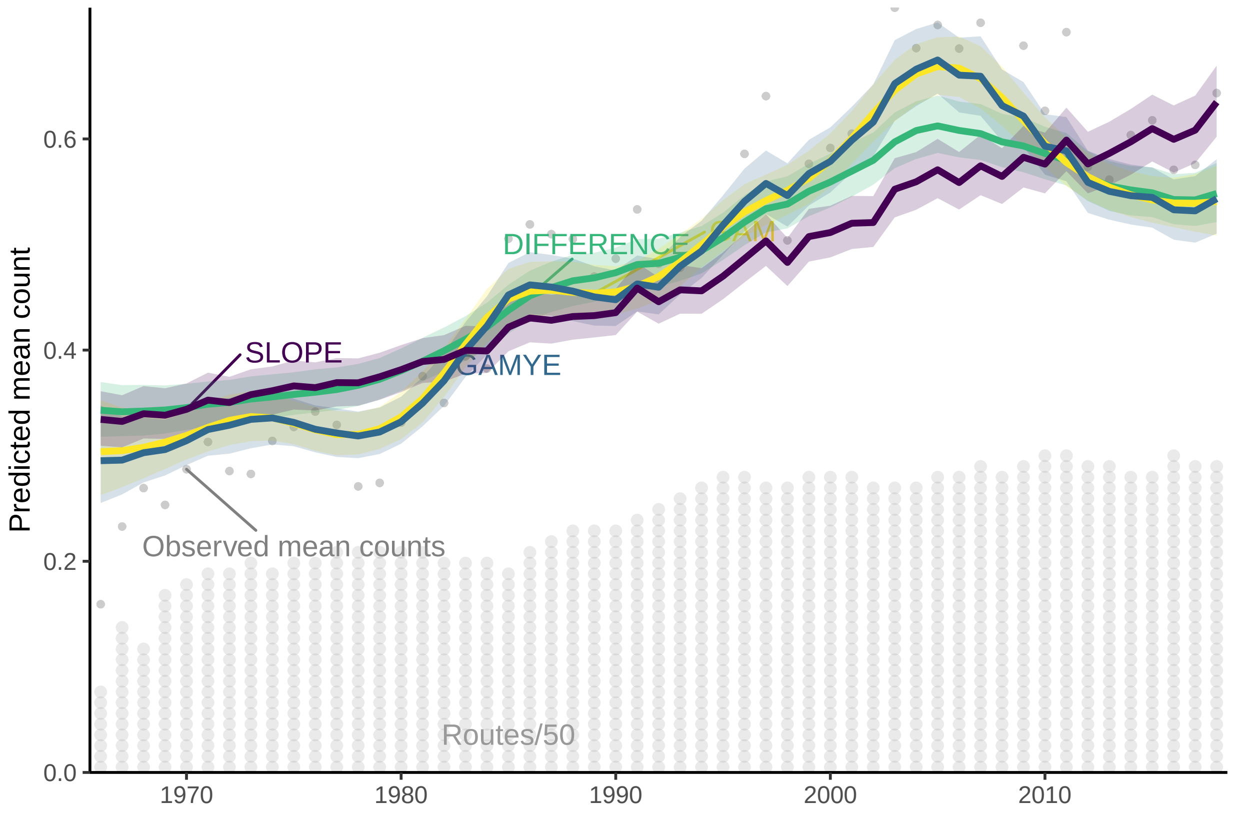


Figure 9: S1.I: Survey-wide population trajectories for Ruby-throated Hummingbird estimated from the BBS using two models described here that include a GAM smoothing function to model change over time (GAM and GAMYE) the standard regression-based model used for BBS status and trend assessments since 2011 (SLOPE), and a first-difference time-series model (DIFFERENCE). The stacked dots along the x-axis indicate the approximate number of BBS counts used in the model in each year; each dot represents 50 counts

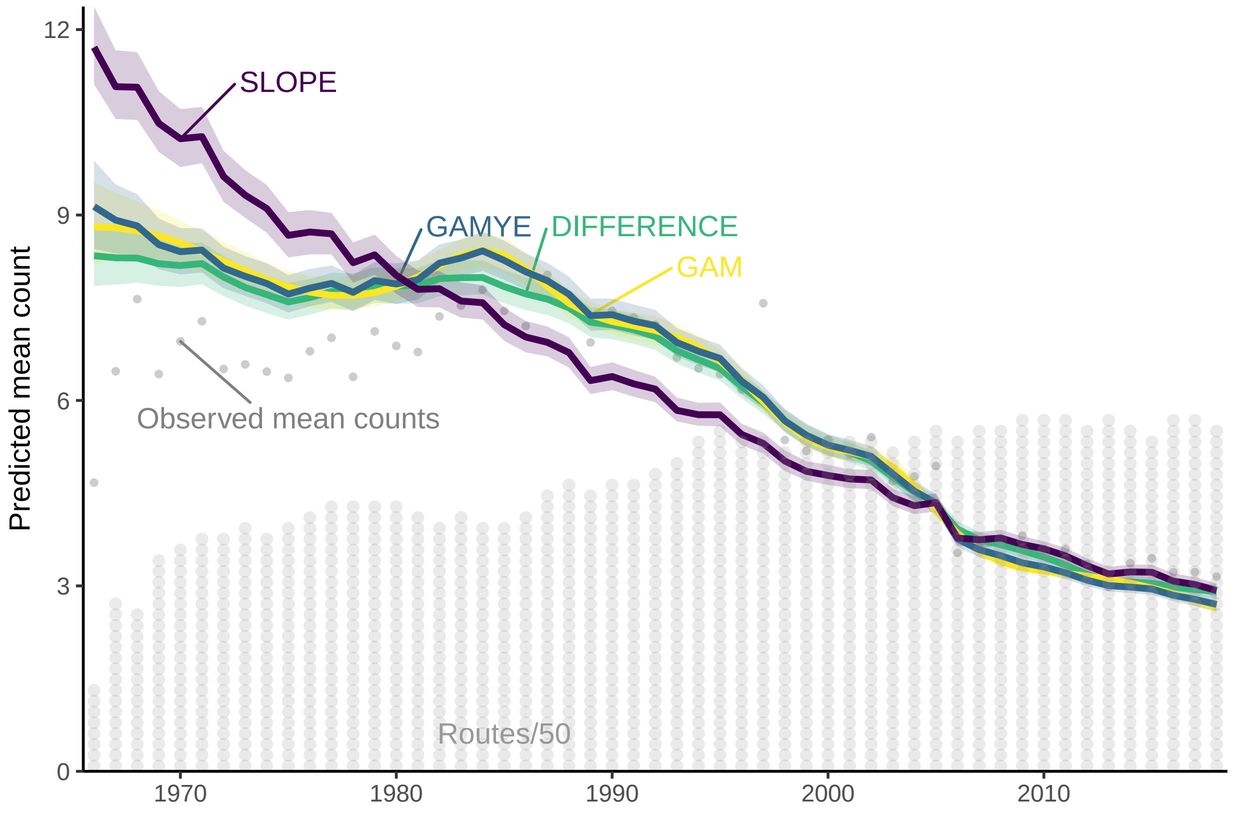


Figure 10: S1.J: Survey-wide population trajectories for Chimney Swift estimated from the BBS using two models described here that include a GAM smoothing function to model change over time (GAM and GAMYE) the standard regression-based model used for BBS status and trend assessments since 2011 (SLOPE), and a first-difference time-series model (DIFFERENCE). The stacked dots along the x-axis indicate the approximate number of BBS counts used in the model in each year; each dot represents 50 counts

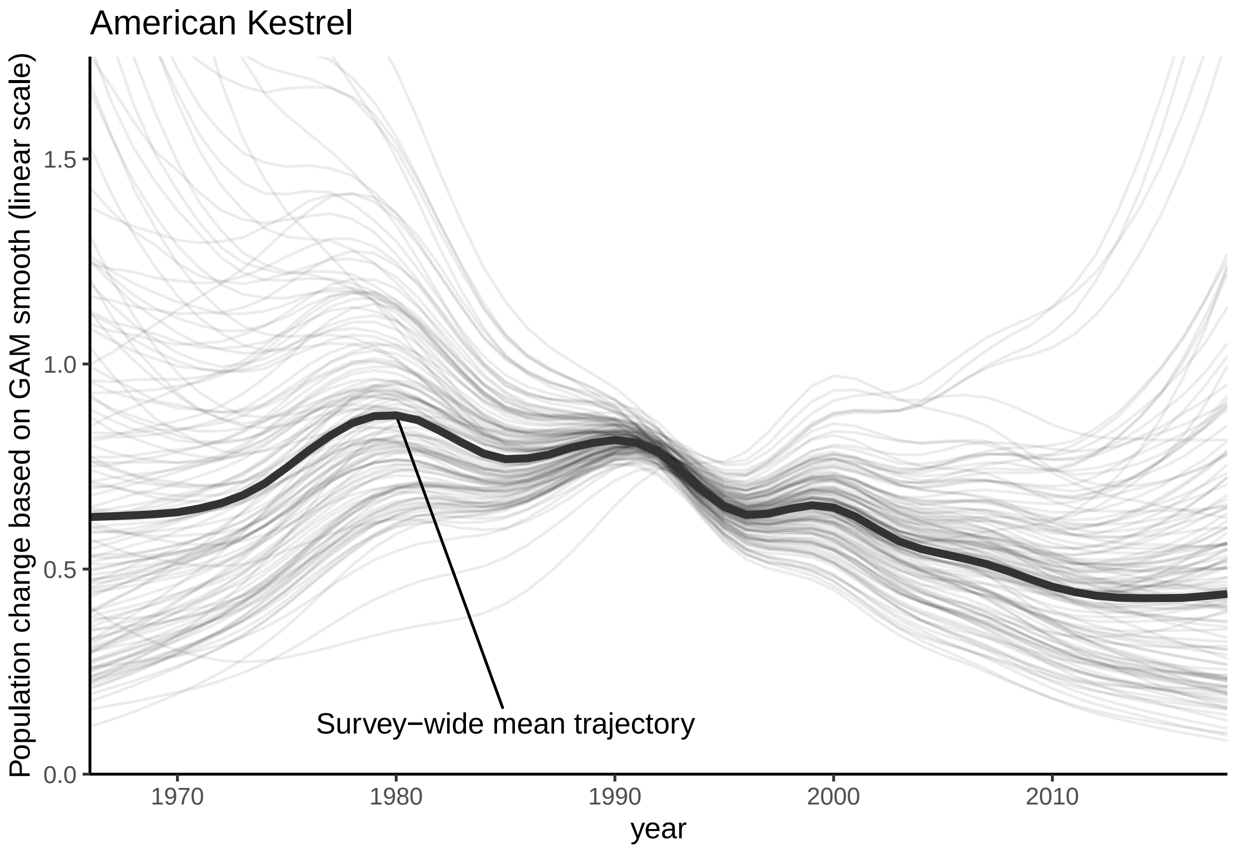


Figure 11: S2.A: Variation among the spatial strata in the random effect smooth components of the GAMYE model applied to American Kestrel data from the BBS. Grey lines show the strata level random effect smooths, and the black lines shows the survey wide mean trajectory.

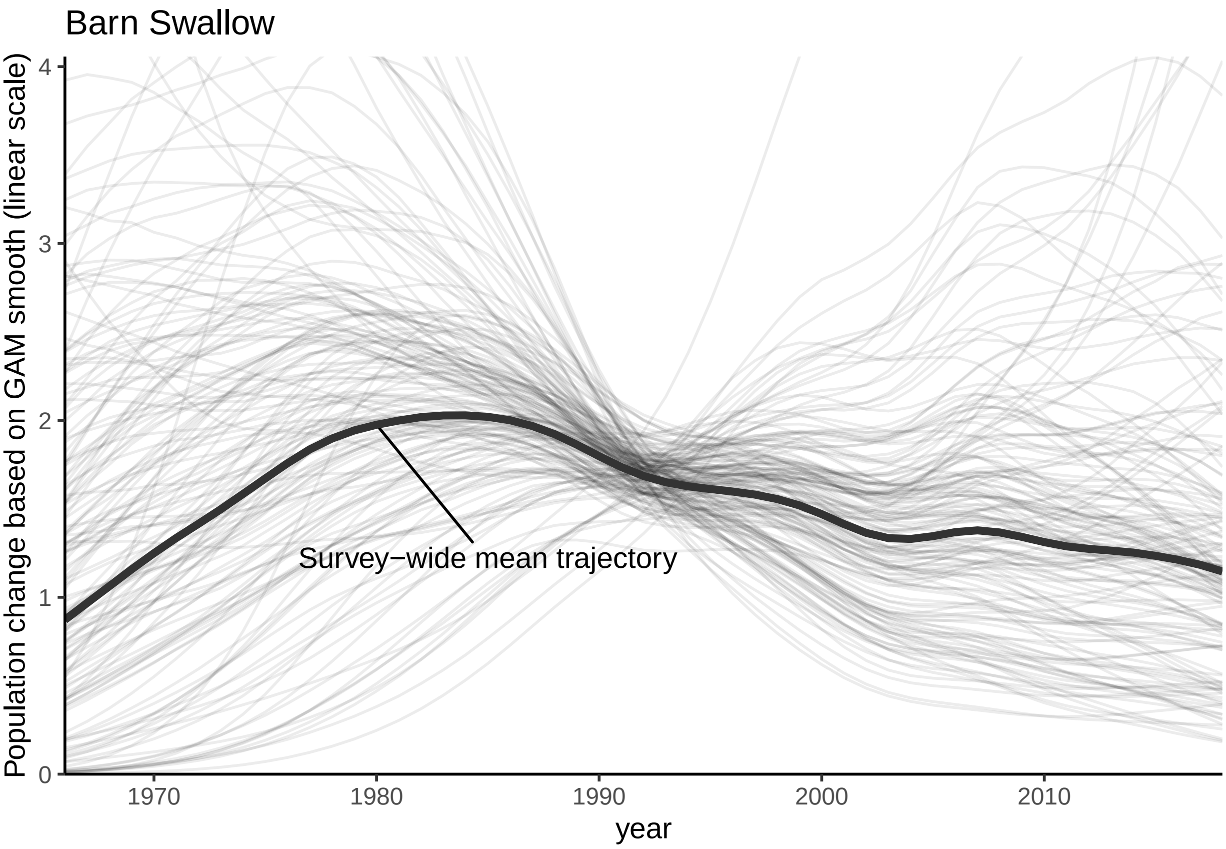


Figure 12: S2.B: Variation among the spatial strata in the random effect smooth components of the GAMYE model applied to Barn Swallow data from the BBS. Grey lines show the strata level random effect smooths, and the black lines shows the survey wide mean trajectory.

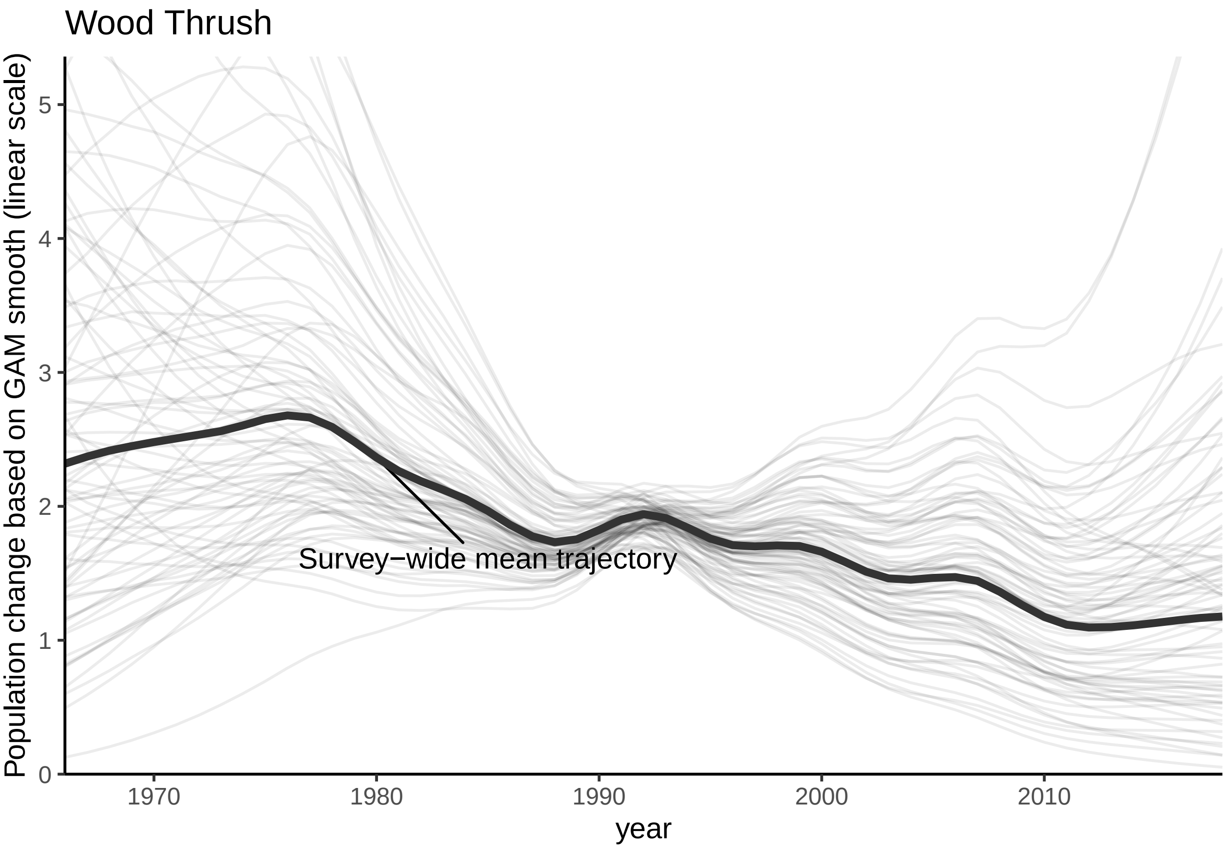


Figure 13: S2.C: Variation among the spatial strata in the random effect smooth components of the GAMYE model applied to Wood Thrush data from the BBS. Grey lines show the strata level random effect smooths, and the black lines shows the survey wide mean trajectory.

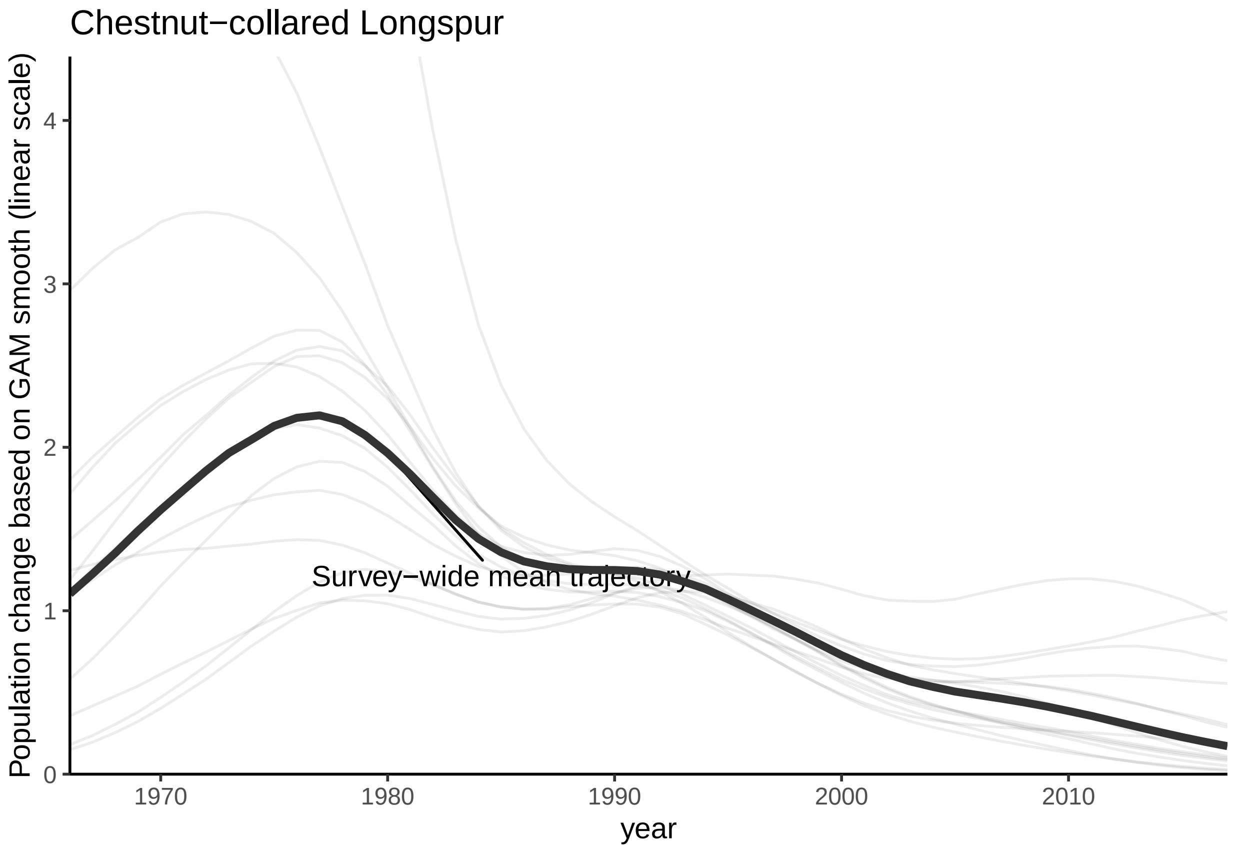


Figure 14: S2.D: Variation among the spatial strata in the random effect smooth components of the GAMYE model applied to Chestnut-collared Longspur data from the BBS. Grey lines show the strata level random effect smooths, and the black lines shows the survey wide mean trajectory.

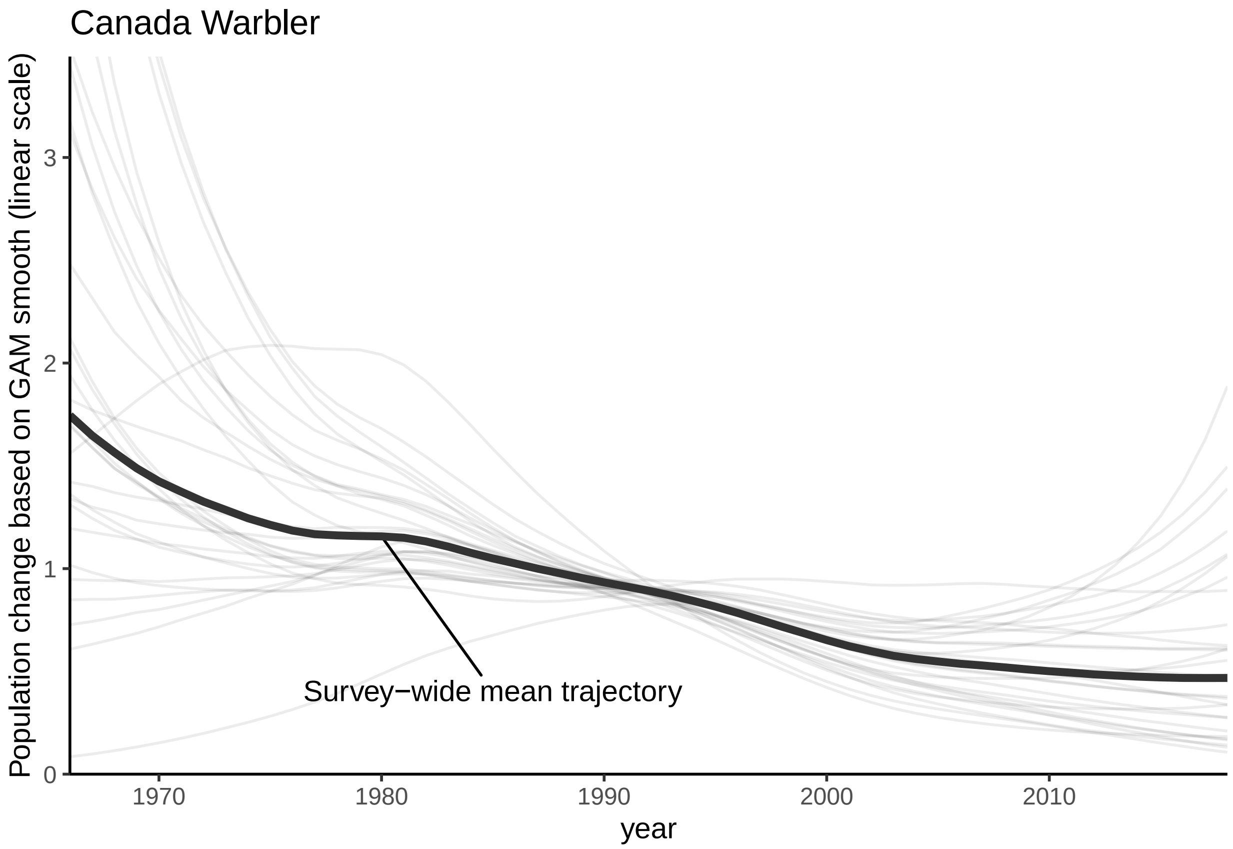


Figure 15: S2.E: Variation among the spatial strata in the random effect smooth components of the GAMYE model applied to Canada Warbler data from the BBS. Grey lines show the strata level random effect smooths, and the black lines shows the survey wide mean trajectory.

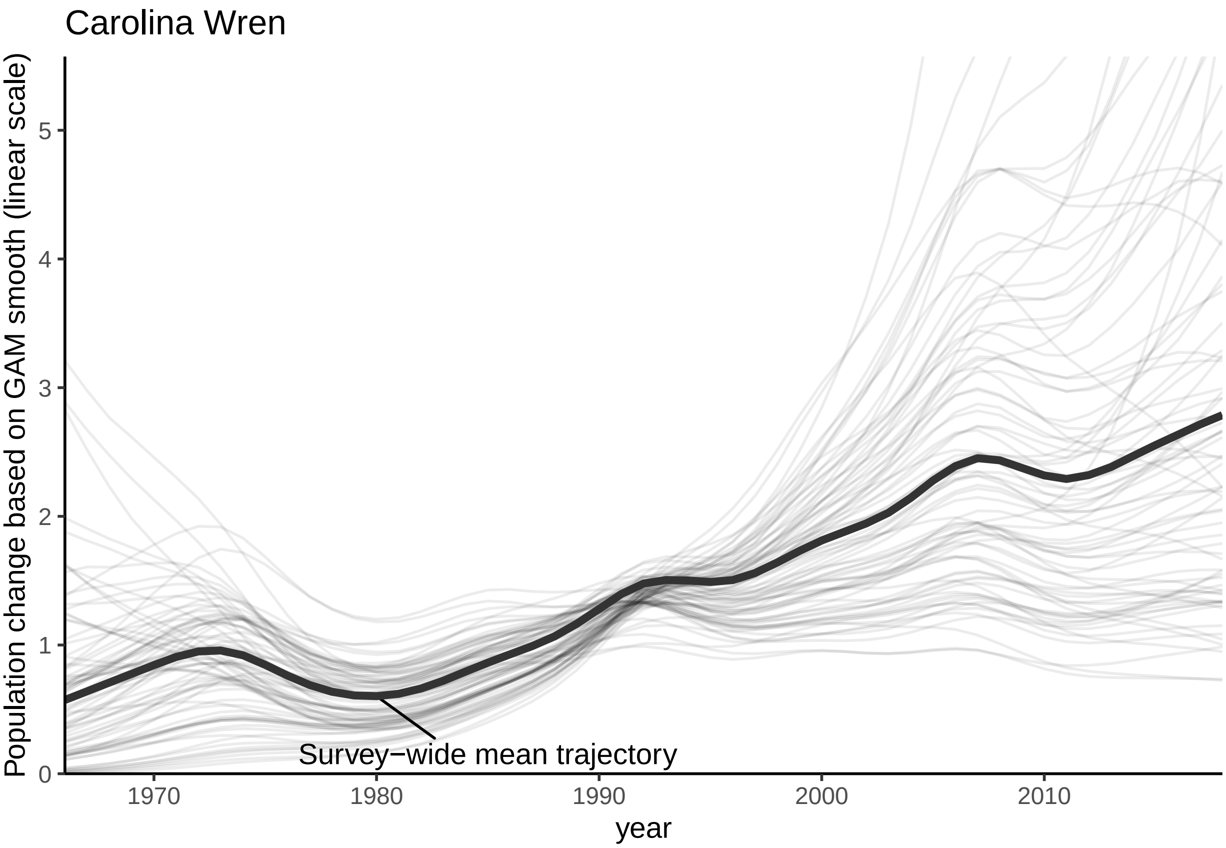


Figure 16: S2.F: Variation among the spatial strata in the random effect smooth components of the GAMYE model applied to Carolina Wren data from the BBS. Grey lines show the strata level random effect smooths, and the black lines shows the survey wide mean trajectory.

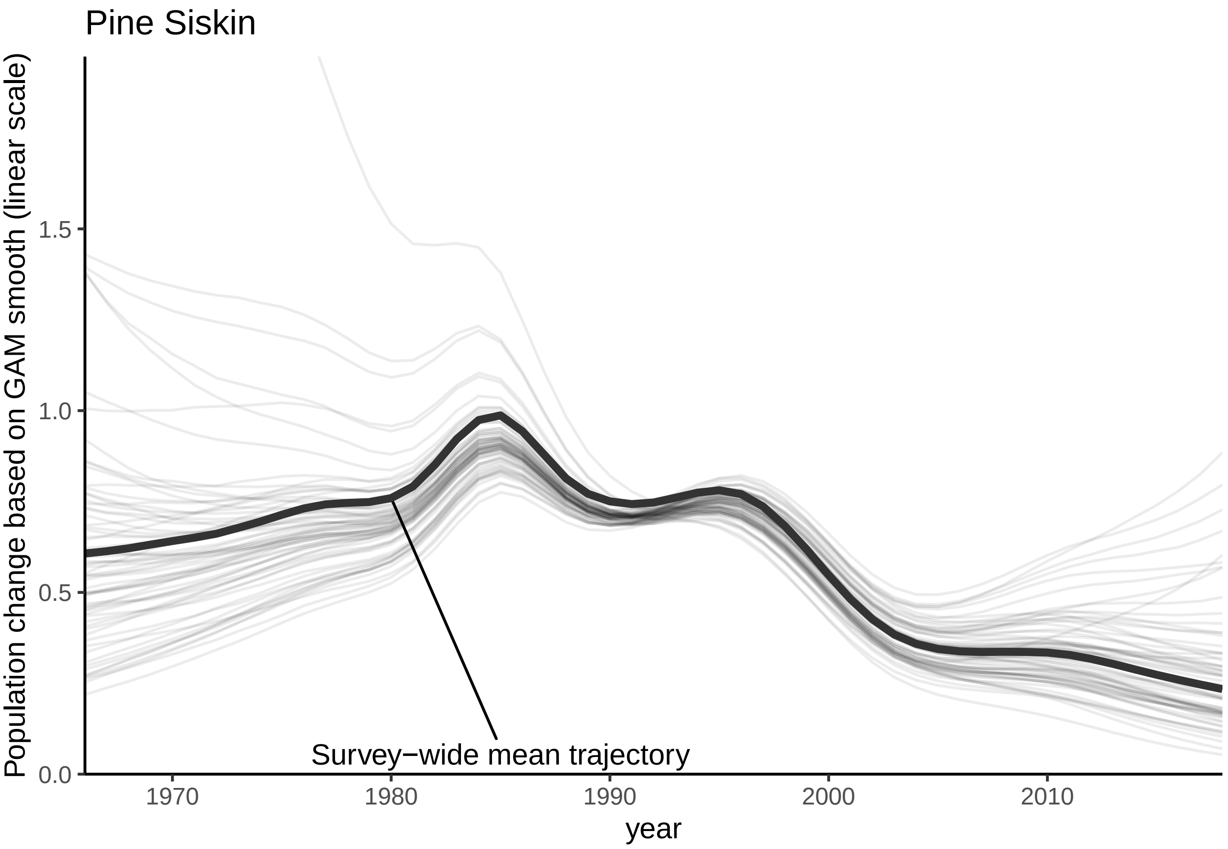


Figure 17: S2.G: Variation among the spatial strata in the random effect smooth components of the GAMYE model applied to Pine Siskin data from the BBS. Grey lines show the strata level random effect smooths, and the black lines shows the survey wide mean trajectory.

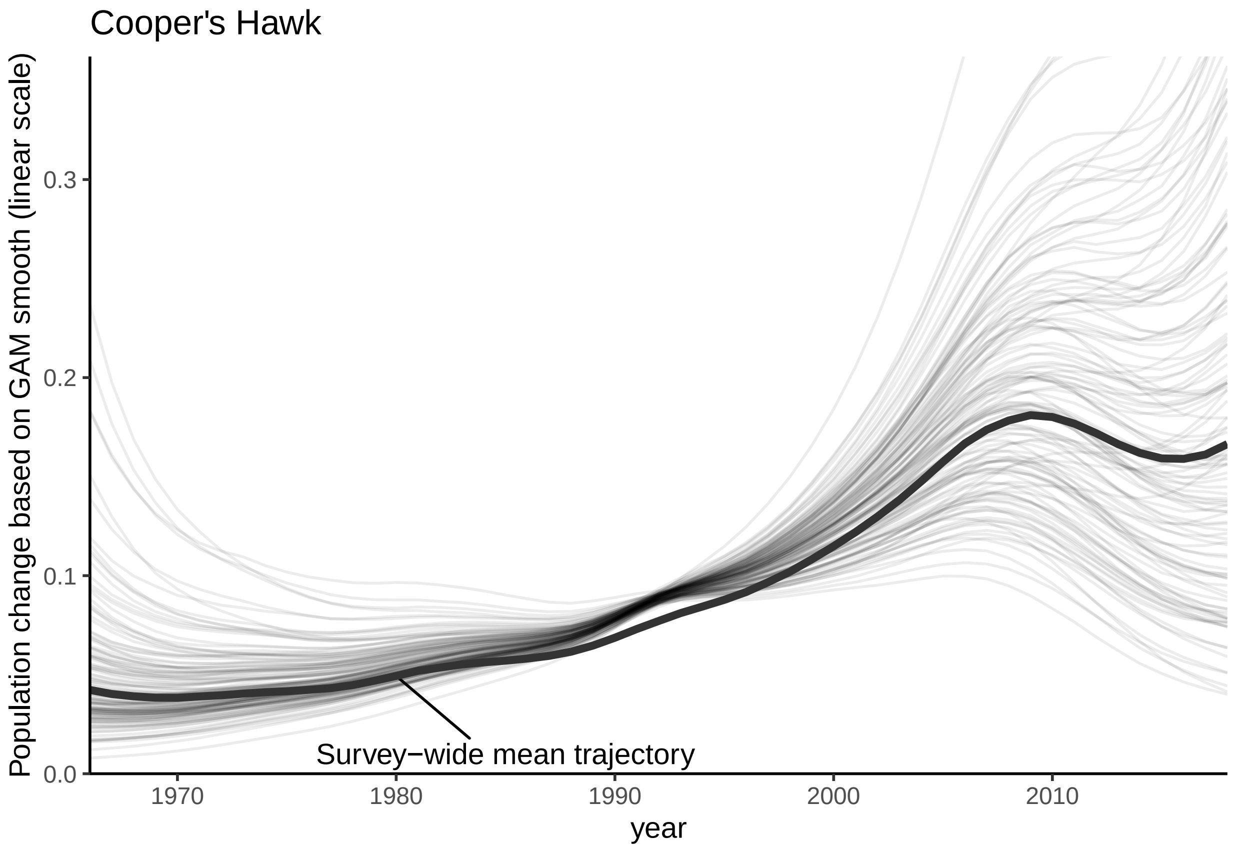


Figure 18: S2.H: Variation among the spatial strata in the random effect smooth components of the GAMYE model applied to Cooper's Hawk data from the BBS. Grey lines show the strata level random effect smooths, and the black lines shows the survey wide mean trajectory.

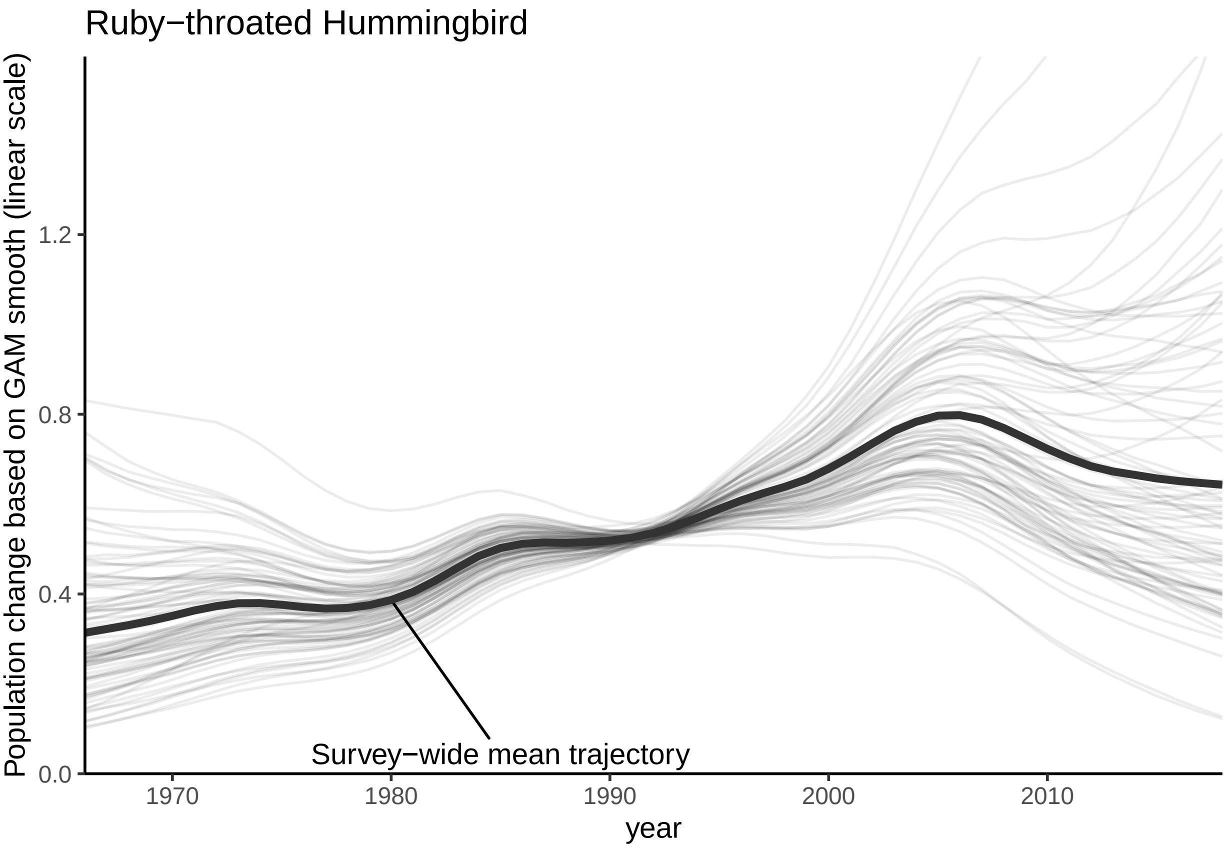


Figure 19: S2.I: Variation among the spatial strata in the random effect smooth components of the GAMYE model applied to Ruby-throated Hummingbird data from the BBS. Grey lines show the strata level random effect smooths, and the black lines shows the survey wide mean trajectory.

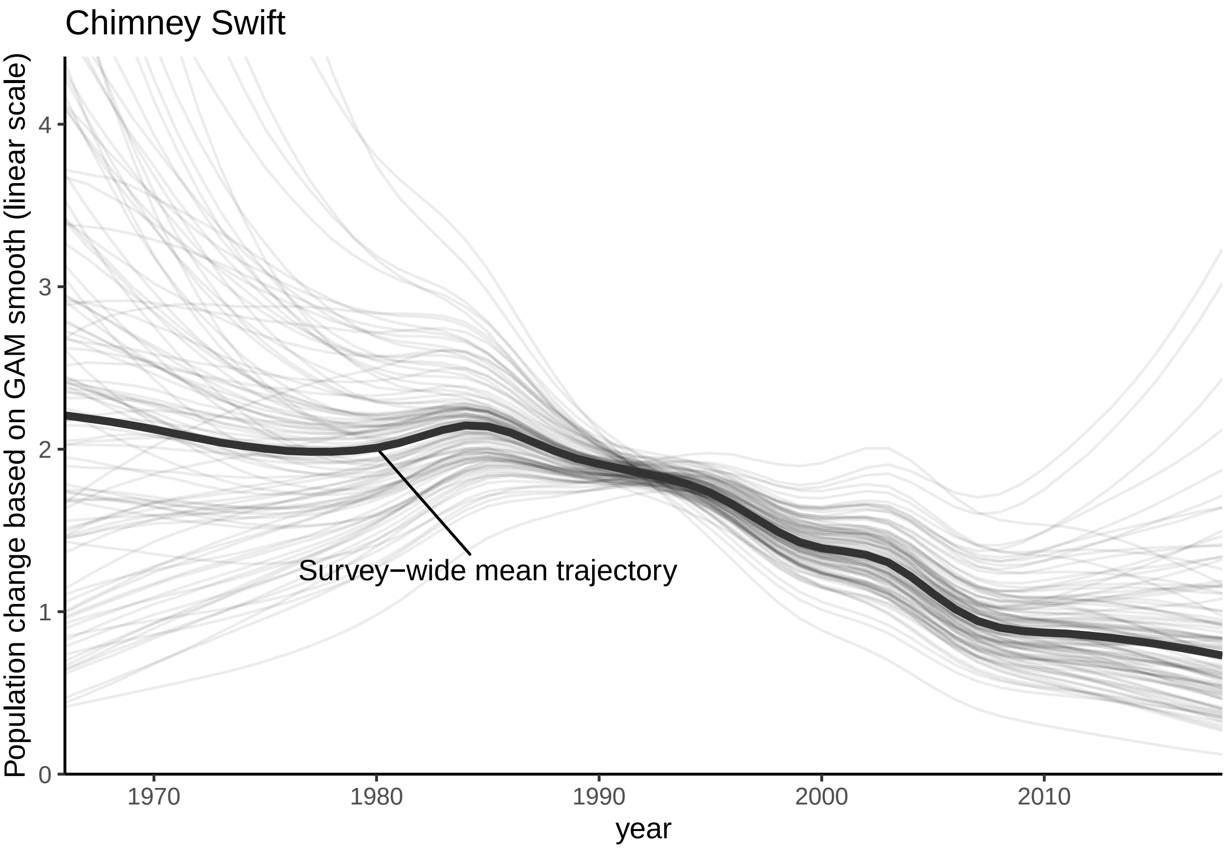


Figure 20: S2.J: Variation among the spatial strata in the random effect smooth components of the GAMYE model applied to Chimney Swift data from the BBS. Grey lines show the strata level random effect smooths, and the black lines shows the survey wide mean trajectory.

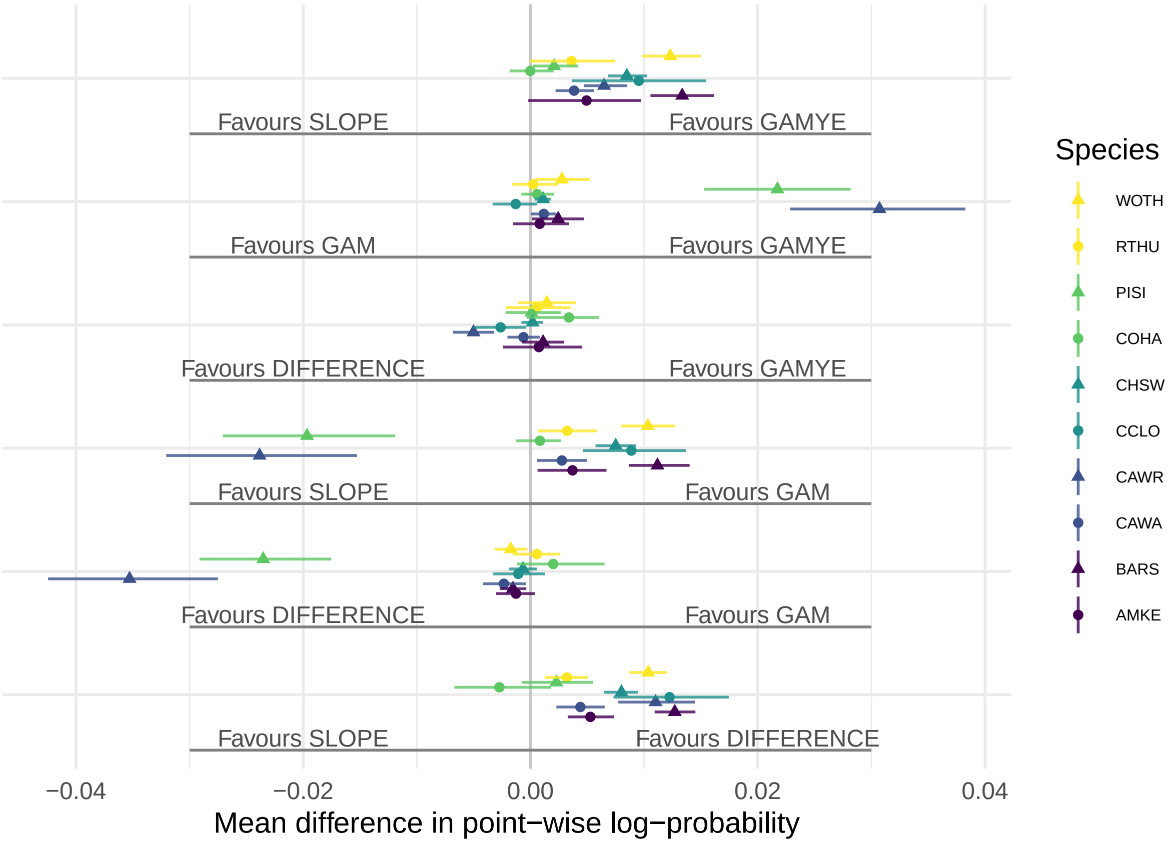


Figure 21: S3.A: Overall pair wise differences in predictive fit between all models for Barn Swallow (BARS) and 9 other selected species. Species short forms are: WOTH is Wood Thrush (Hylocichla mustelina), RTHU is Ruby-throated Hummingbird (Archilochus colubris), PISI is Pine Siskin (Spinus pinus), COHA is Cooper's Hawk (Accipiter cooperii), CHSW is Chimney Swift (Chaetura pelagica), CCLO is Chestnut-collared Longspur (Calcarius ornatus), CAWR is Carolina Wren (Thryothorus ludovicianus), CAWA is Canada Warbler (Cardellina canadensis), MAKE is American Kestrel (Falco sparverius).
