## Supplemental Figures S4-S7 for "North American Breeding Bird Survey status and trend estimates to inform a wide-range of conservation needs, using a flexible Bayesian hierarchical generalized additive model"

### Supplemental Material Figures S4 to S7, Smith and Edwards 2020, Improved status and trend estimates from the North American Breeding Bird Survey using a Bayesian hierarchical generalized additive model

```
## `geom_smooth()` using formula 'y ~ x'
```

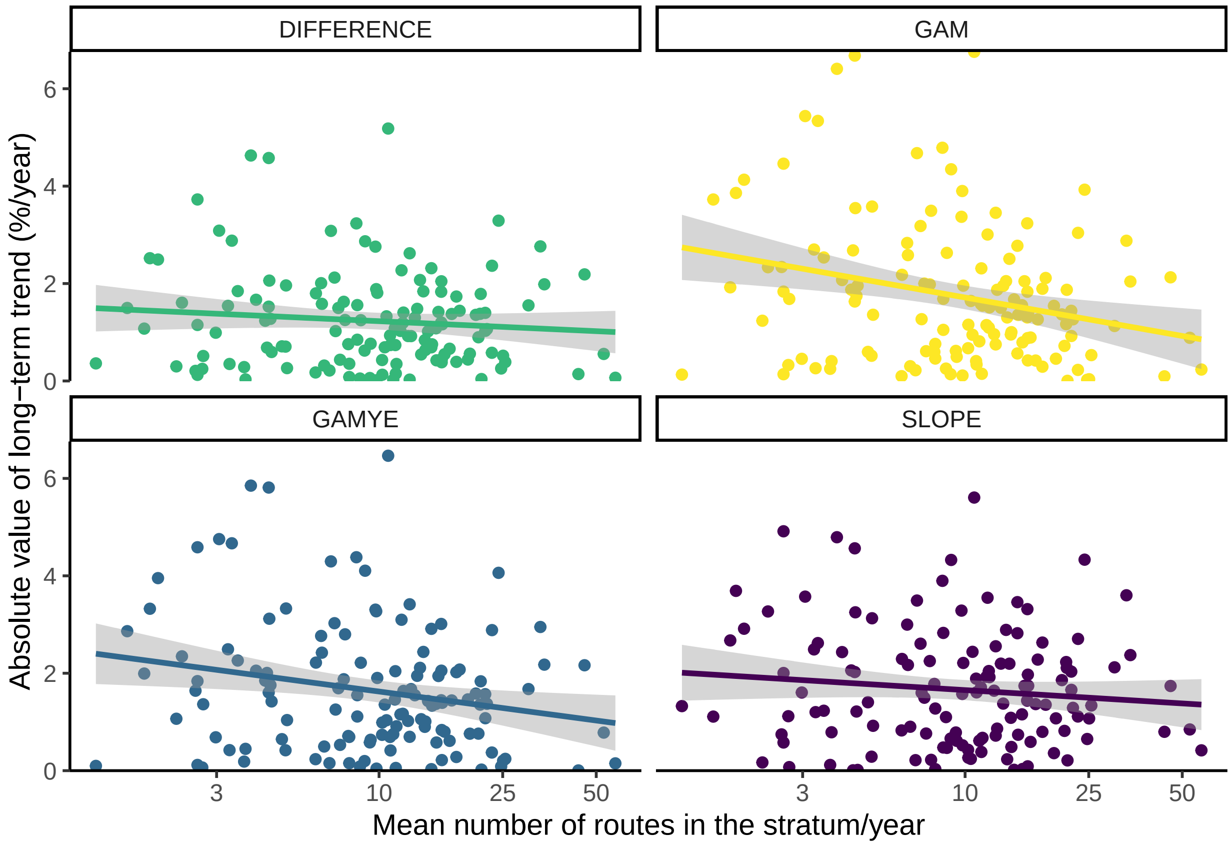


Figure 1: S4.A: Relationship between the absolute value of estimated long-term trends (1966-2018) and the amount of data in each stratum, from the four models compared here for American Kestrel.

```
## `geom_smooth()` using formula 'y ~ x'

## `geom_smooth()` using formula 'y ~ x'

## `geom_smooth()` using formula 'y ~ x'

## `geom_smooth()` using formula 'y ~ x'

## `geom_smooth()` using formula 'y ~ x'

## `geom_smooth()` using formula 'y ~ x'

## Warning: Removed 11 rows containing missing values (geom_smooth).
```

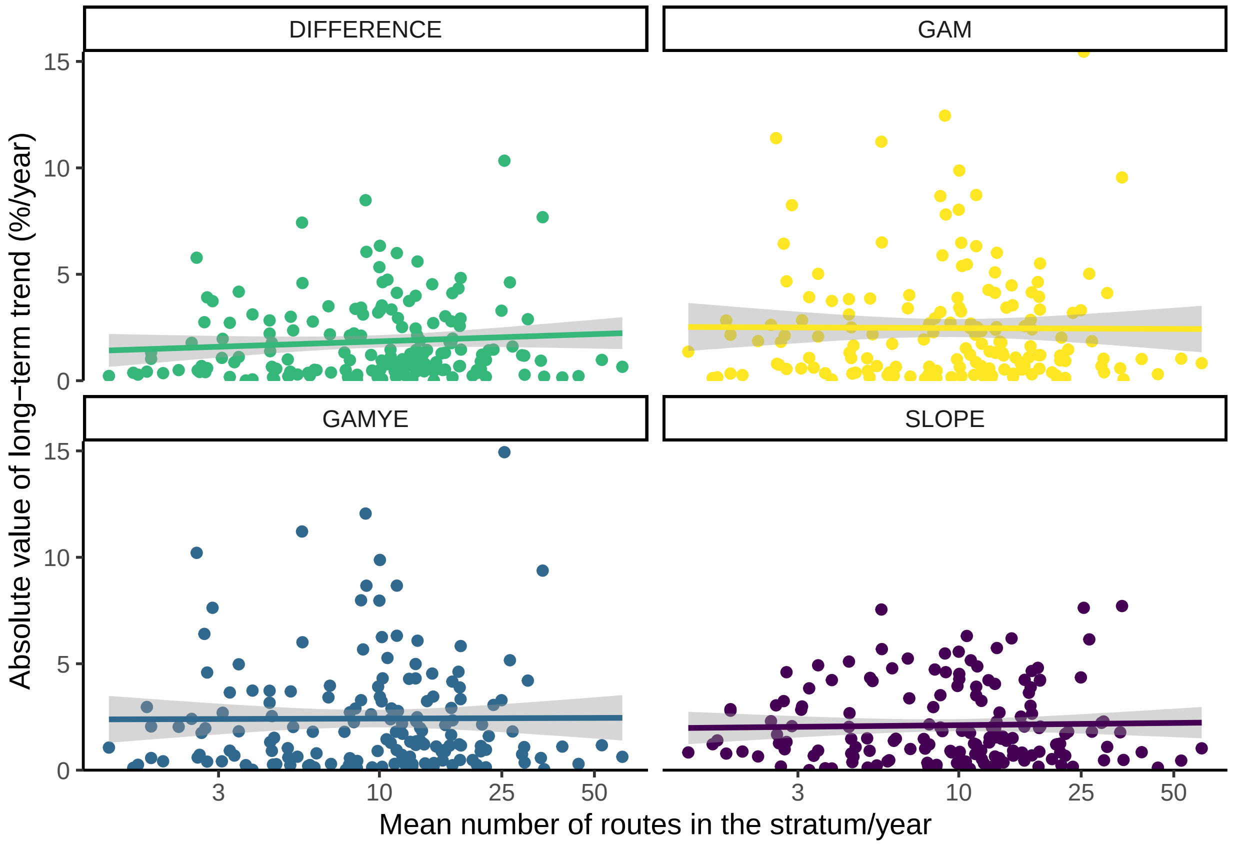


Figure 2: S4.B: Relationship between the absolute value of estimated long-term trends (1966-2018) and the amount of data in each stratum, from the four models compared here for Barn Swallow.

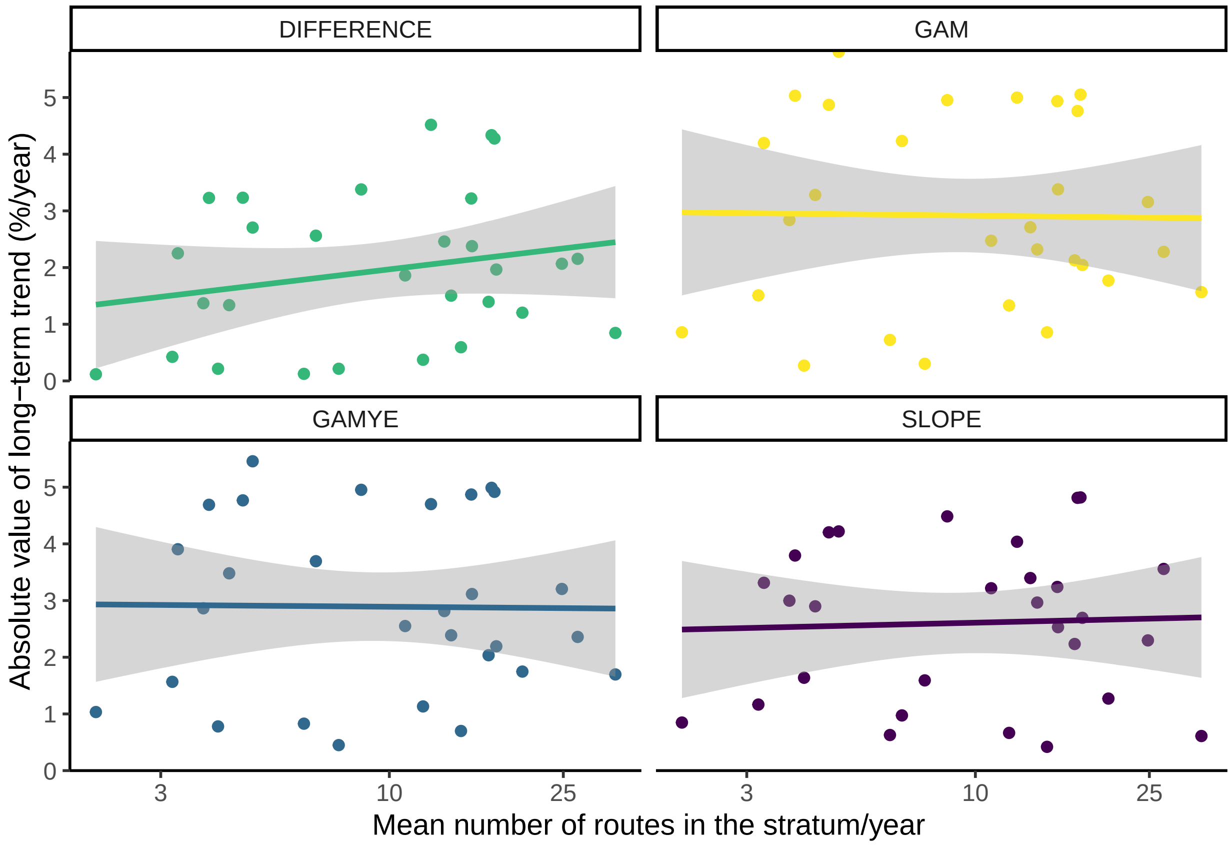


Figure 3: S4.C: Relationship between the absolute value of estimated long-term trends (1966-2018) and the amount of data in each stratum, from the four models compared here for Canada Warbler.

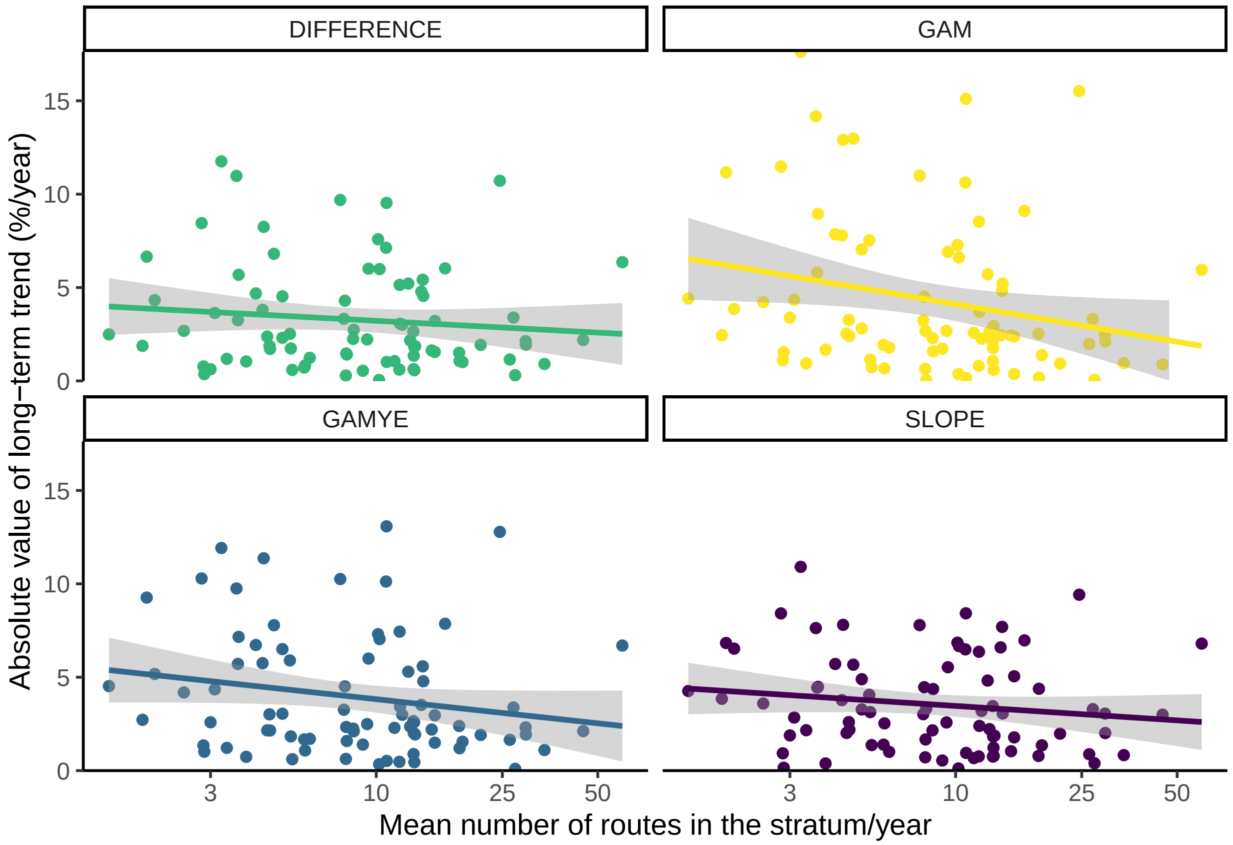


Figure 4: S4.D: Relationship between the absolute value of estimated long-term trends (1966-2018) and the amount of data in each stratum, from the four models compared here for Carolina Wren.

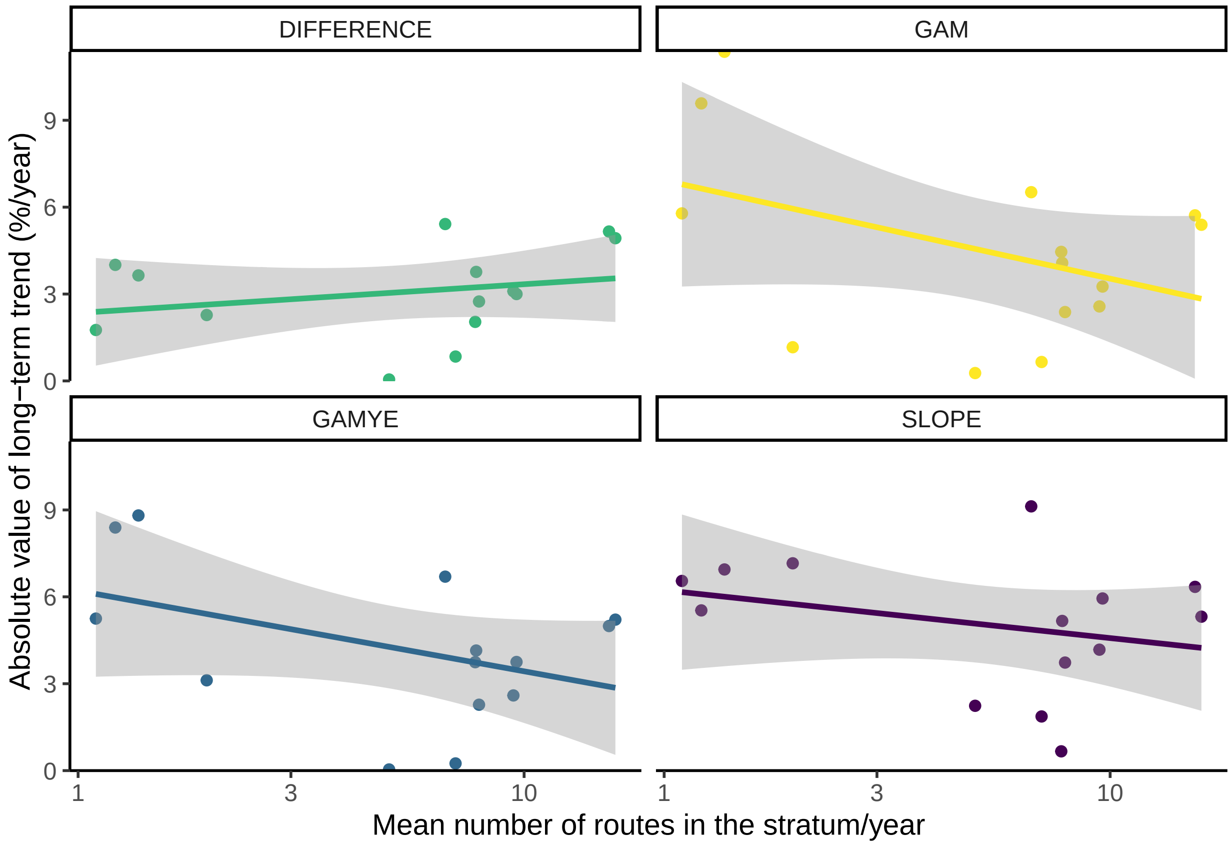


Figure 5: S4.E: Relationship between the absolute value of estimated long-term trends (1966-2018) and the amount of data in each stratum, from the four models compared here for Chestnut-collared Longspur.

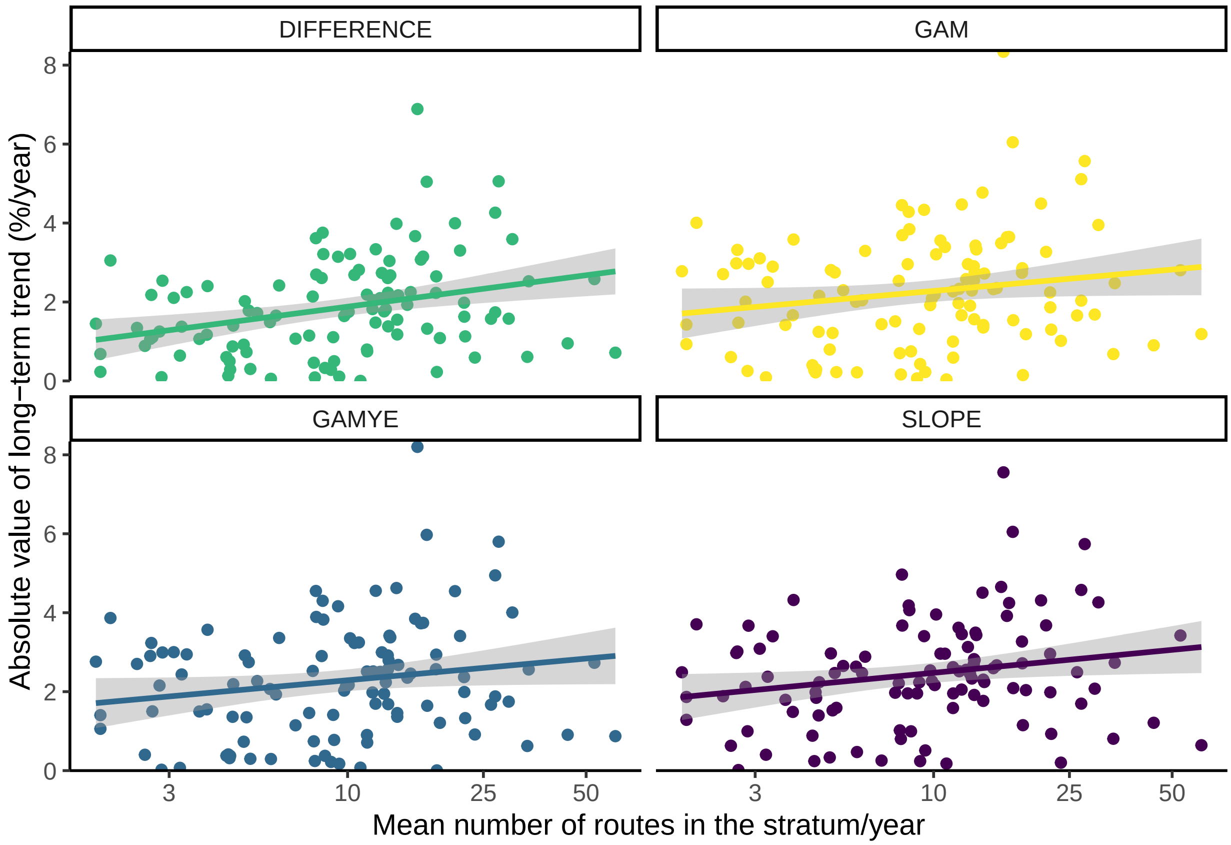


Figure 6: S4.F: Relationship between the absolute value of estimated long-term trends (1966-2018) and the amount of data in each stratum, from the four models compared here for Chimney Swift.

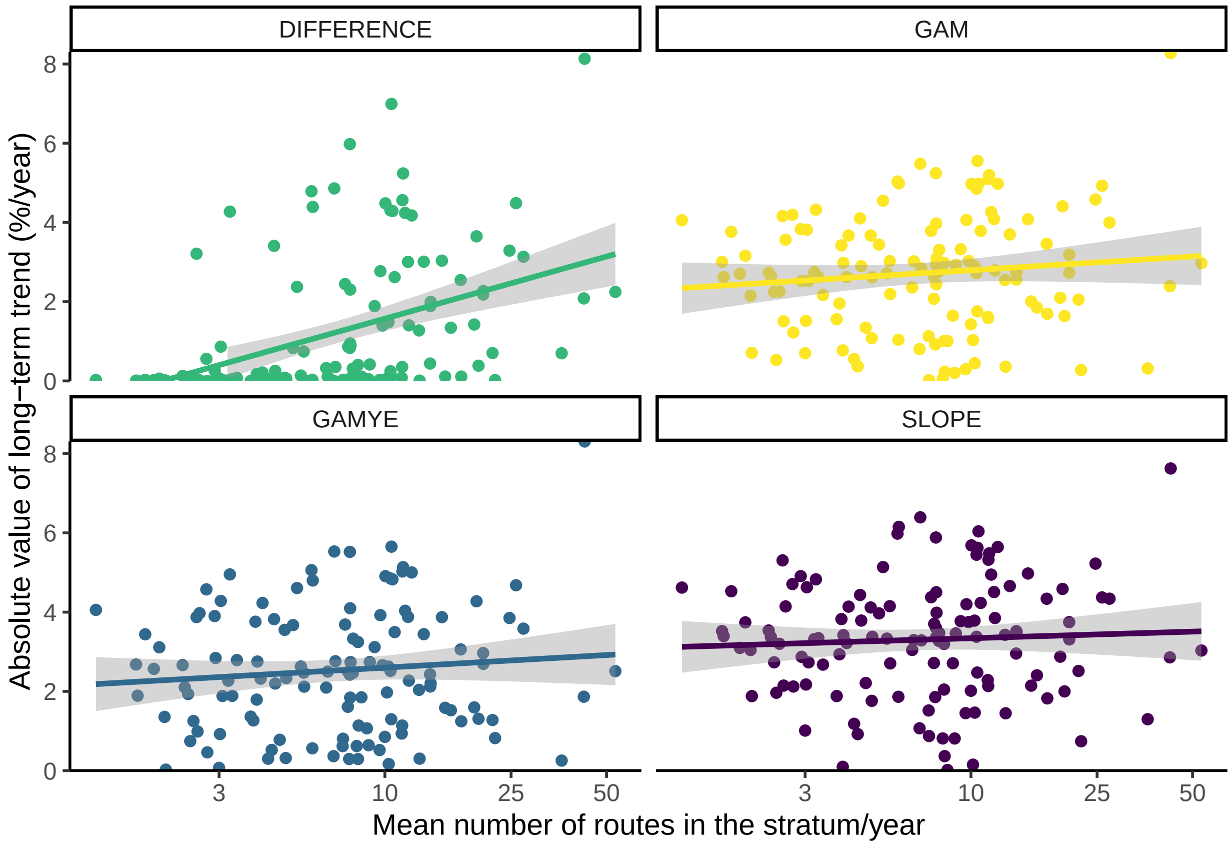


Figure 7: S4.G: Relationship between the absolute value of estimated long-term trends (1966-2018) and the amount of data in each stratum, from the four models compared here for Cooper's Hawk.

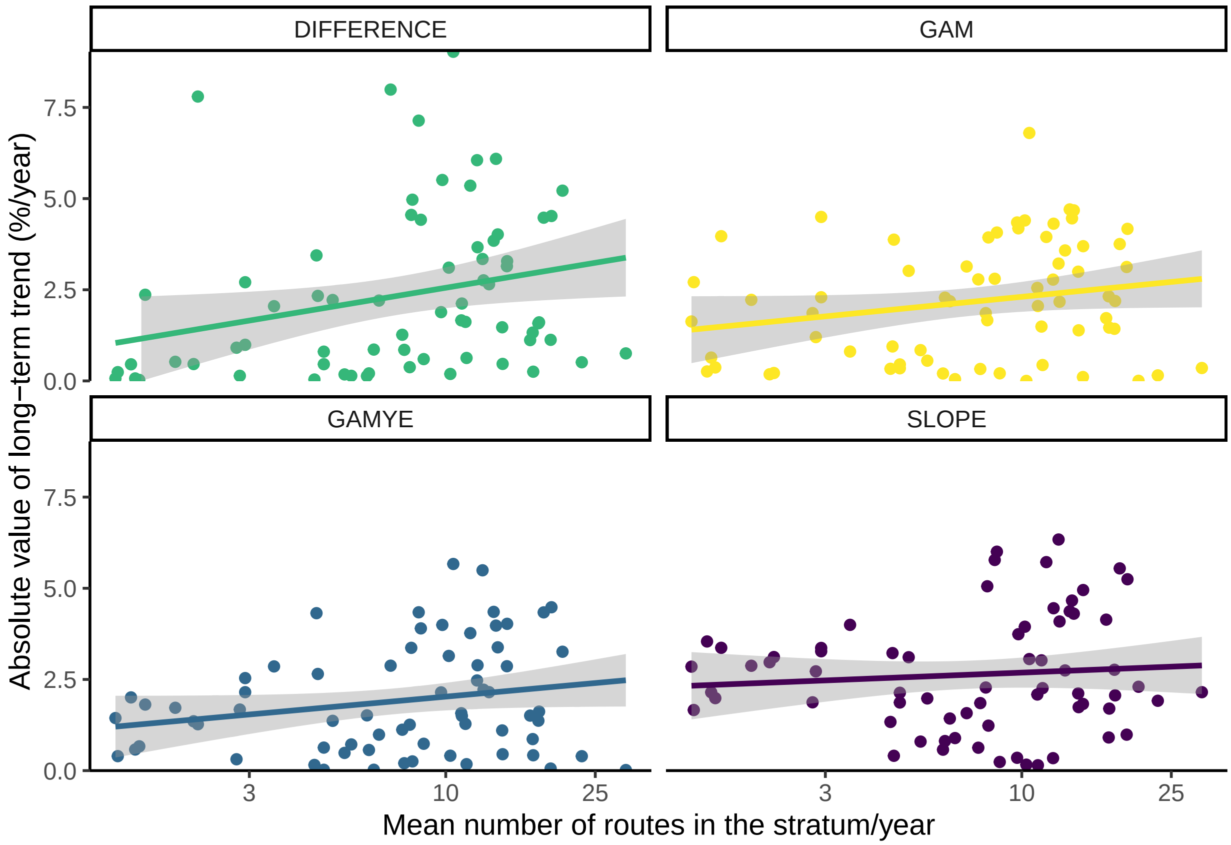


Figure 8: S4.H: Relationship between the absolute value of estimated long-term trends (1966-2018) and the amount of data in each stratum, from the four models compared here for Pine Siskin.

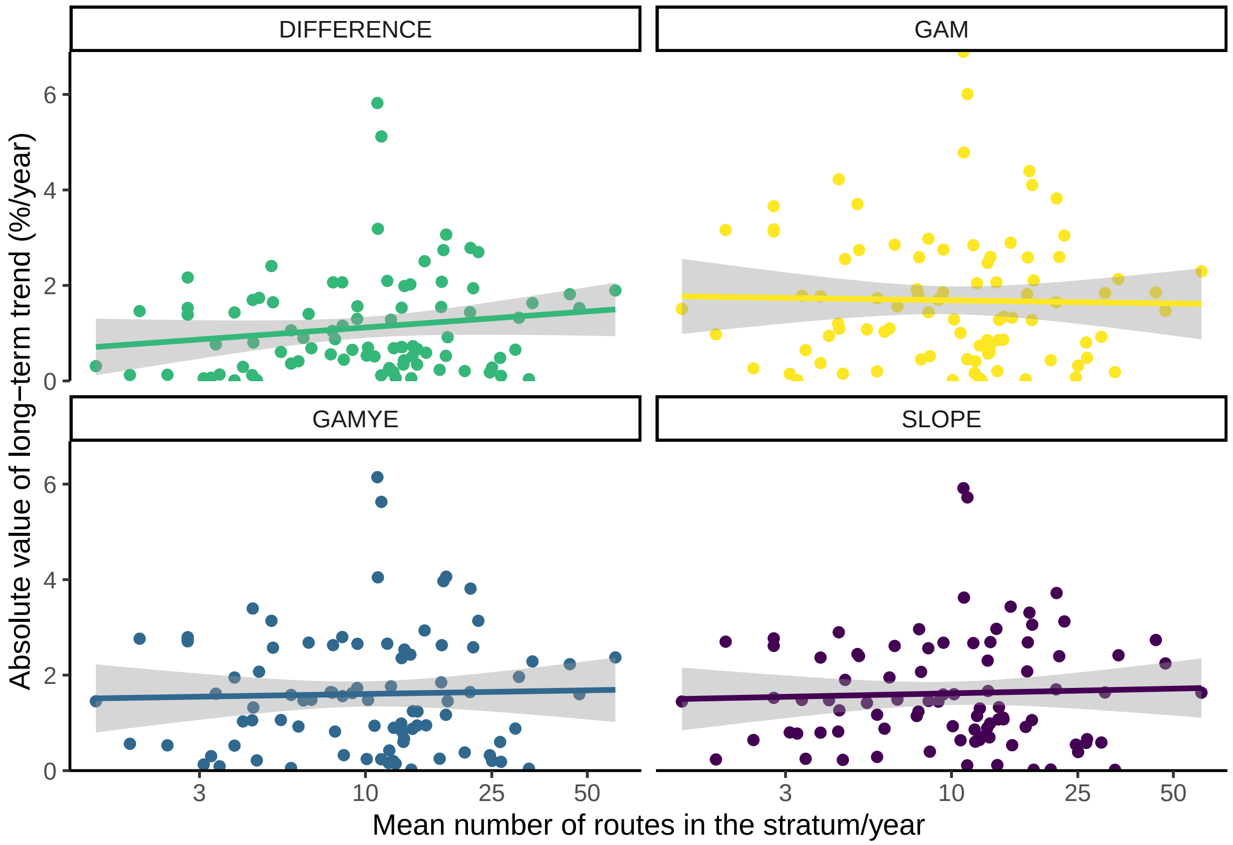


Figure 9: S4.I: Relationship between the absolute value of estimated long-term trends (1966-2018) and the amount of data in each stratum, from the four models compared here for Ruby-throated Hummingbird.

```
## `geom_smooth()` using formula 'y ~ x'
```

```
## `geom_smooth()` using formula 'y ~ x'
```

```
## `geom_smooth()` using formula 'y ~ x'
```

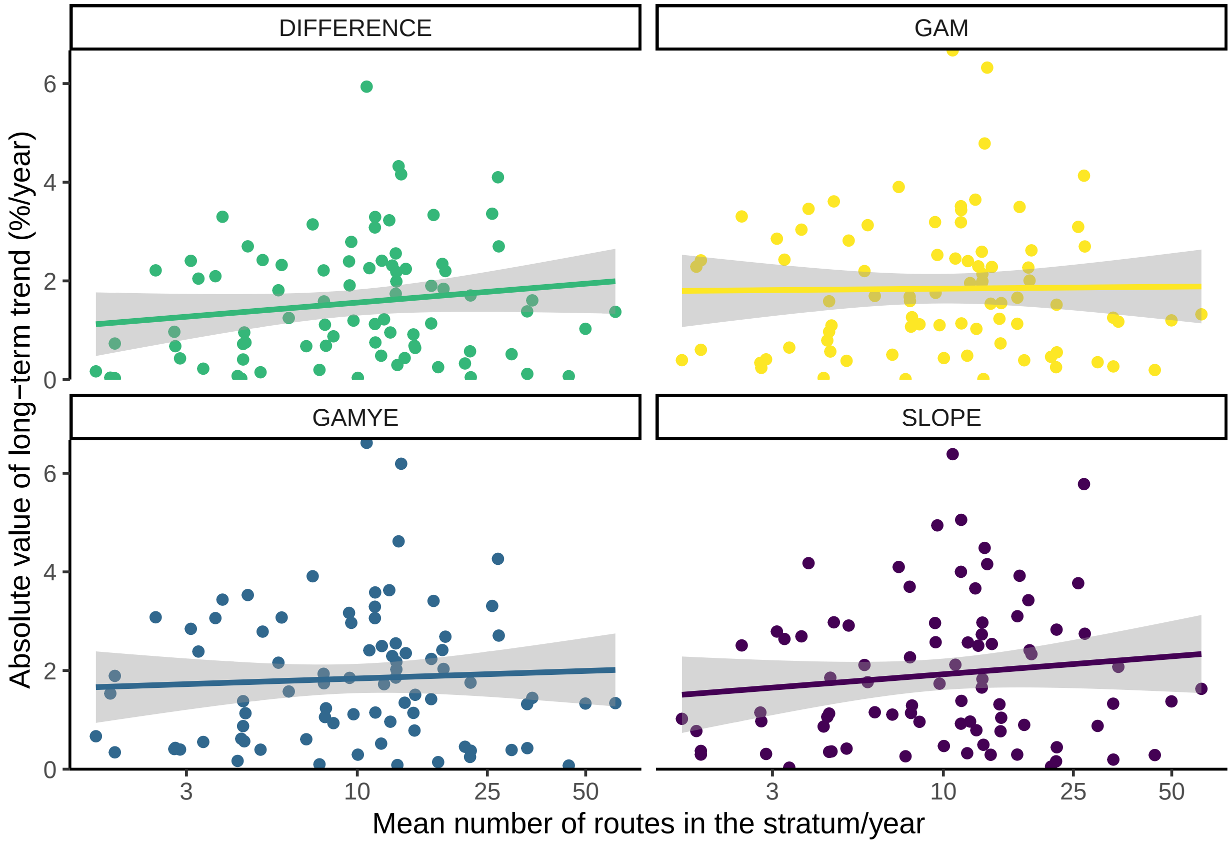


Figure 10: S4.J: Relationship between the absolute value of estimated long-term trends (1966-2018) and the amount of data in each stratum, from the four models compared here for Wood Thrush.

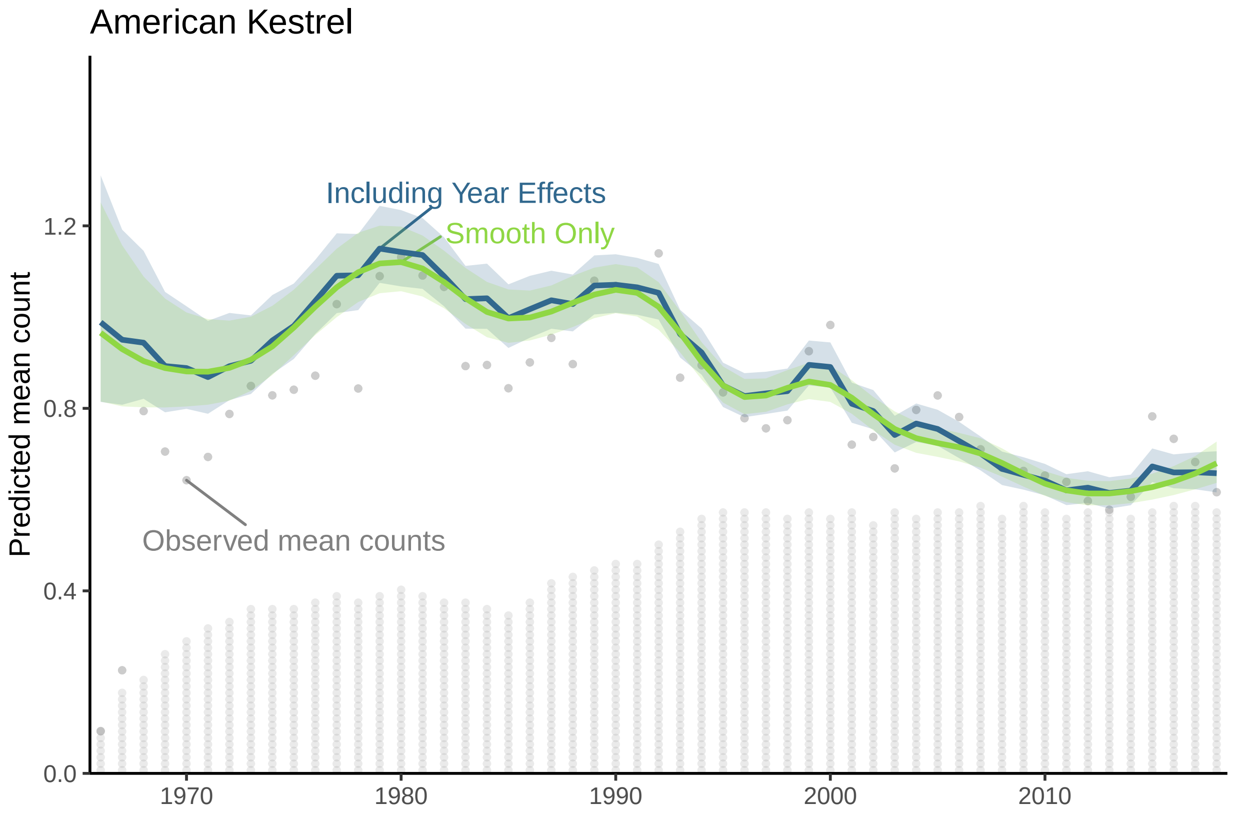


Figure 11: S5.A: Decomposition of the survey-wide population trajectory for American Kestrel from the GAMYE, showing the full trajectory (Including Year Effects) and the isolated smooth component (Smooth Only), which can be used to estimate population trends that are less sensitive to the particular year in which they are estimated. The stacked dots along the x axis indicate the approximate number of BBS counts used in the model; each dot represents 50 counts.

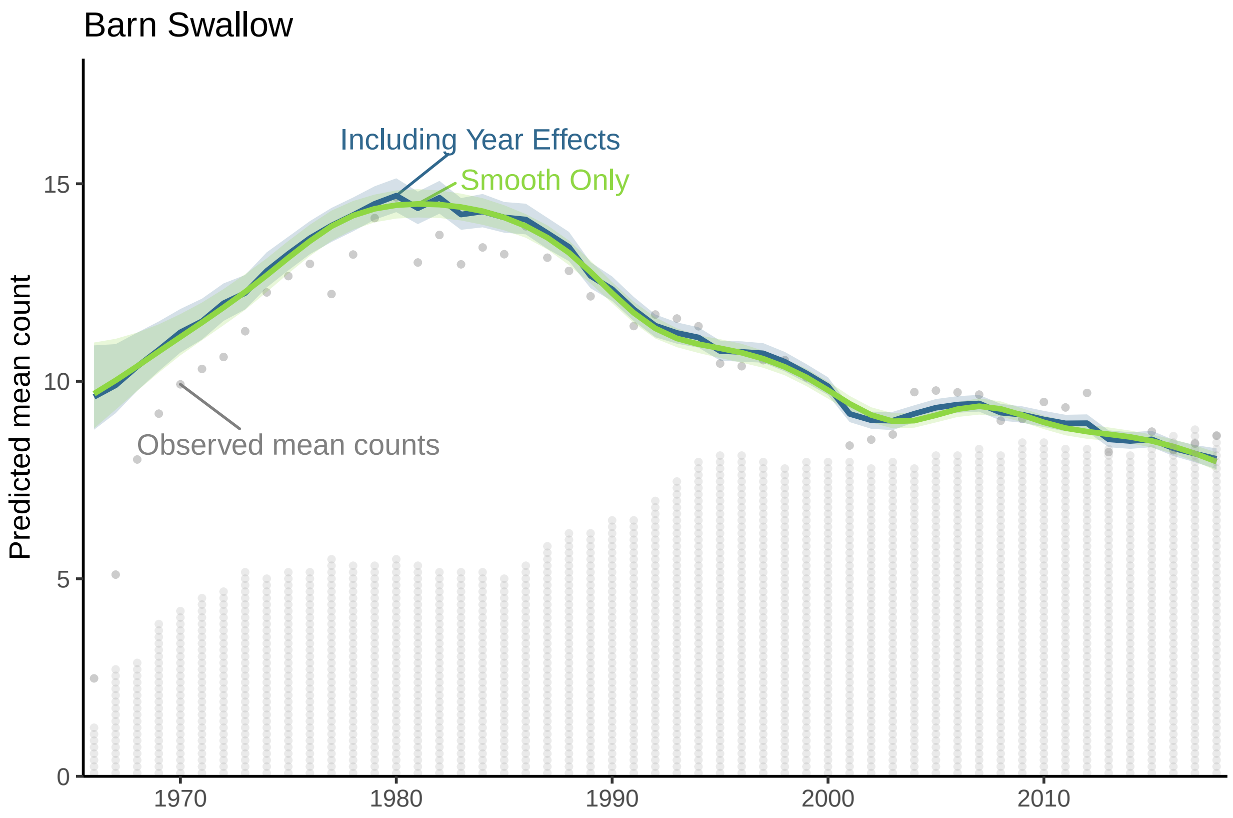


Figure 12: S5.B: Decomposition of the survey-wide population trajectory for Barn Swallow from the GAMYE, showing the full trajectory (Including Year Effects) and the isolated smooth component (Smooth Only), which can be used to estimate population trends that are less sensitive to the particular year in which they are estimated. The stacked dots along the x axis indicate the approximate number of BBS counts used in the model; each dot represents 50 counts.

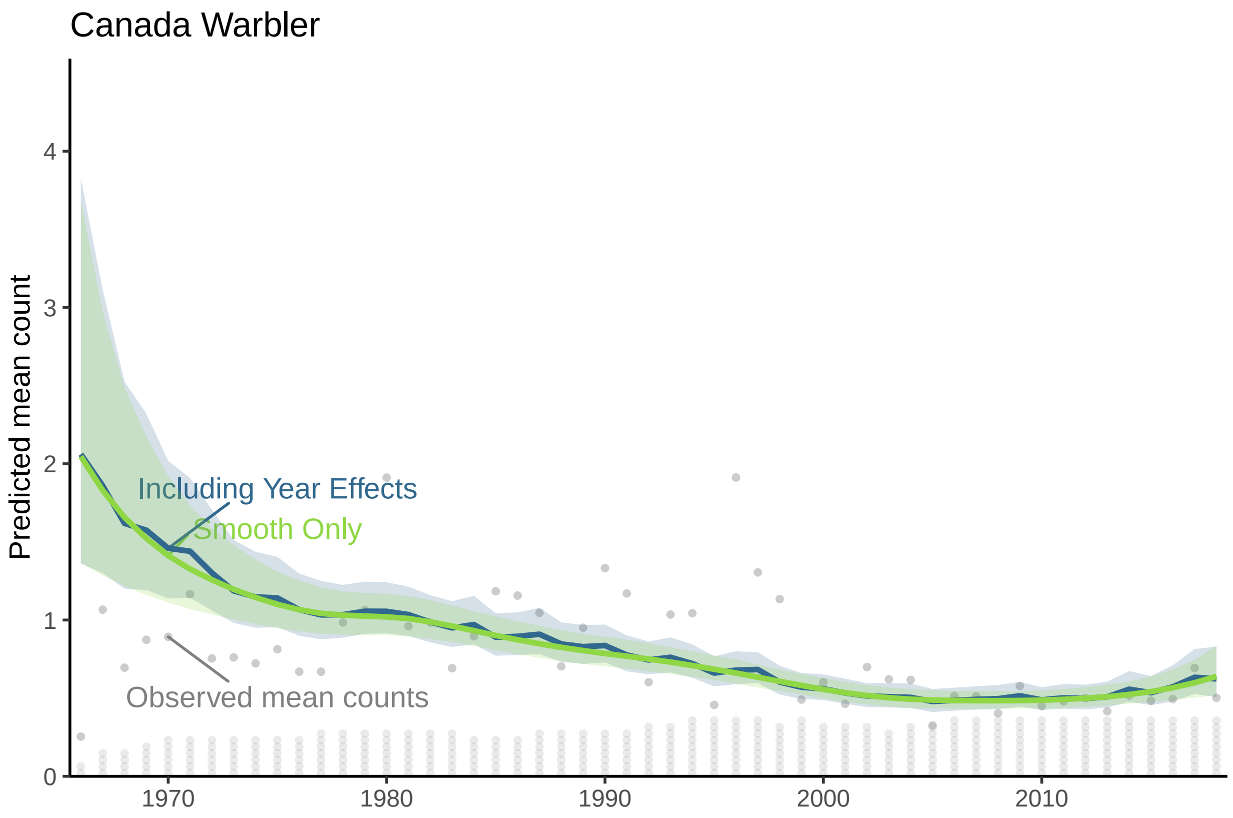


Figure 13: S5.C: Decomposition of the survey-wide population trajectory for Canada Warbler from the GAMYE, showing the full trajectory (Including Year Effects) and the isolated smooth component (Smooth Only), which can be used to estimate population trends that are less sensitive to the particular year in which they are estimated. The stacked dots along the x axis indicate the approximate number of BBS counts used in the model; each dot represents 50 counts.

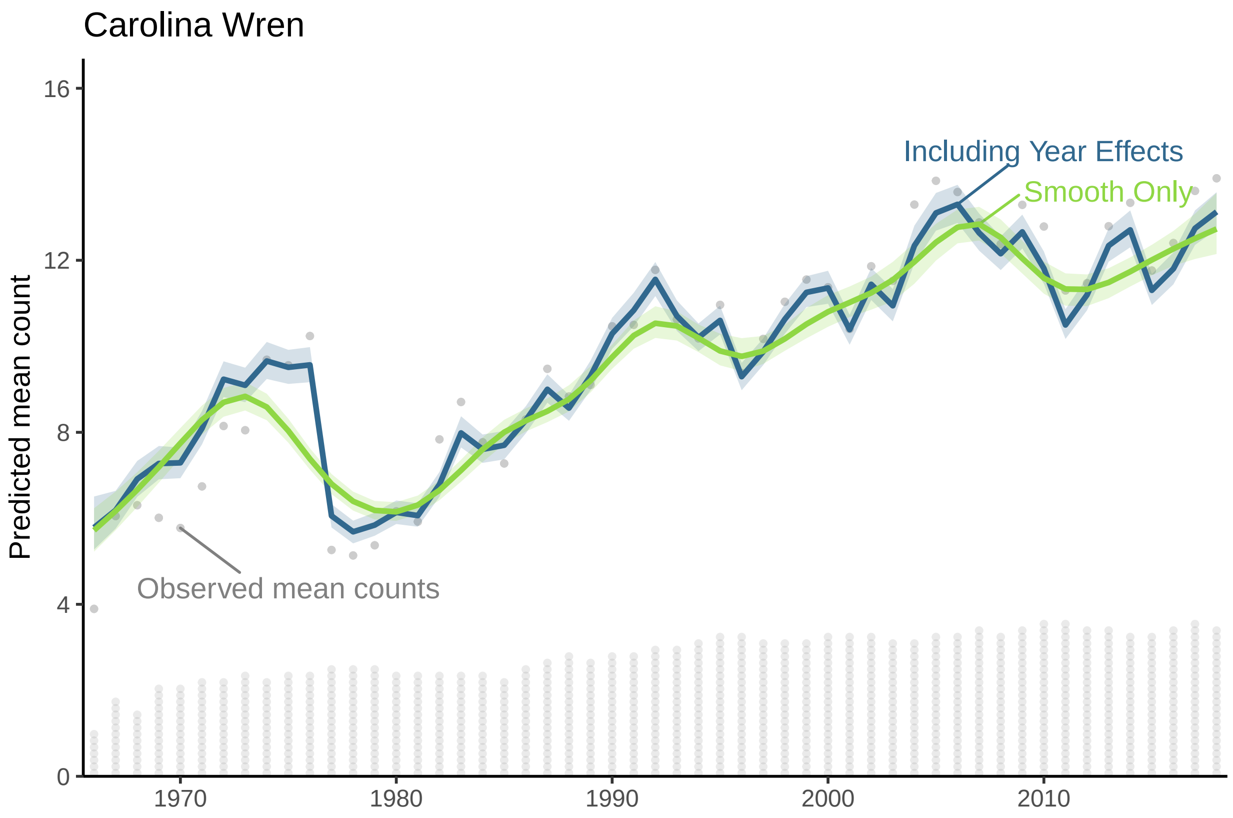


Figure 14: S5.D: Decomposition of the survey-wide population trajectory for Carolina Wren from the GAMYE, showing the full trajectory (Including Year Effects) and the isolated smooth component (Smooth Only), which can be used to estimate population trends that are less sensitive to the particular year in which they are estimated. The stacked dots along the x axis indicate the approximate number of BBS counts used in the model; each dot represents 50 counts.

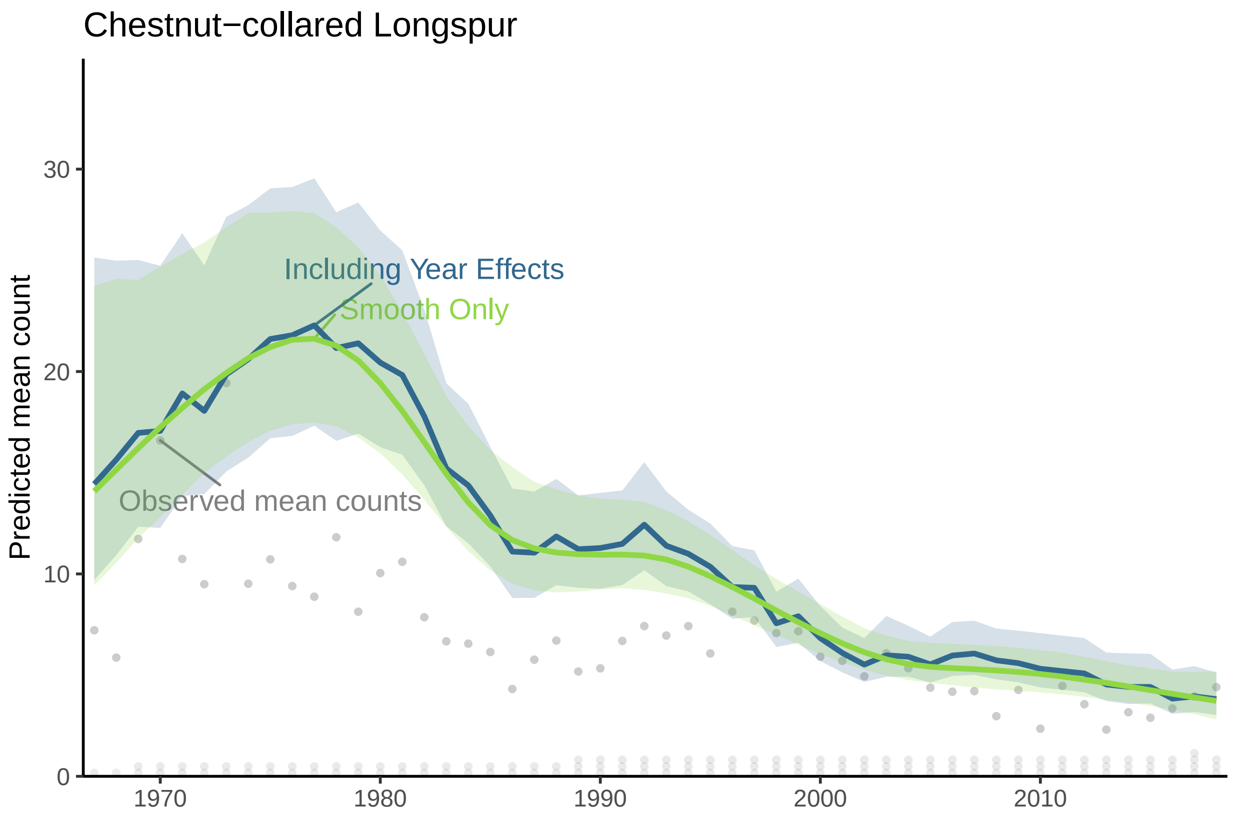


Figure 15: S5.E: Decomposition of the survey-wide population trajectory for Chestnut-collared Longspur from the GAMYE, showing the full trajectory (Including Year Effects) and the isolated smooth component (Smooth Only), which can be used to estimate population trends that are less sensitive to the particular year in which they are estimated. The stacked dots along the x axis indicate the approximate number of BBS counts used in the model; each dot represents 50 counts.

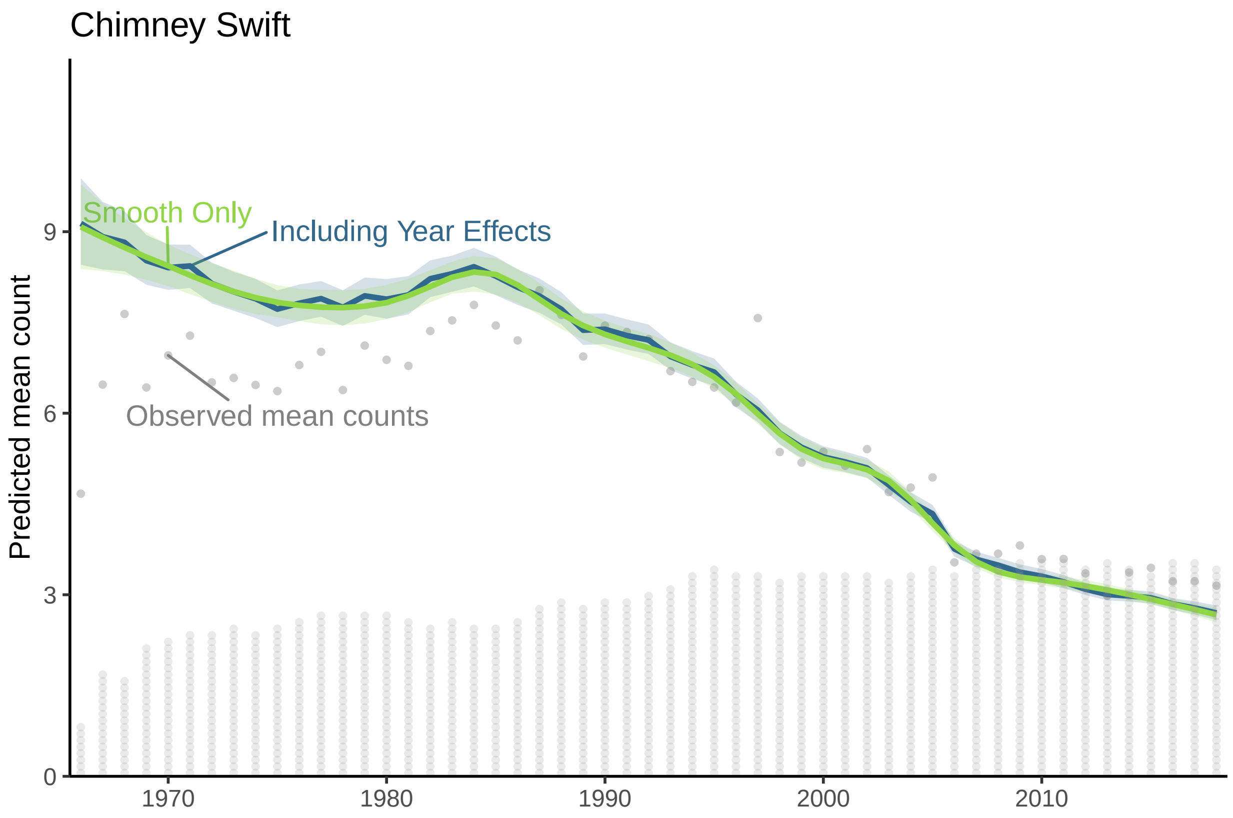


Figure 16: S5.F: Decomposition of the survey-wide population trajectory for Chimney Swift from the GAMYE, showing the full trajectory (Including Year Effects) and the isolated smooth component (Smooth Only), which can be used to estimate population trends that are less sensitive to the particular year in which they are estimated. The stacked dots along the x axis indicate the approximate number of BBS counts used in the model; each dot represents 50 counts.

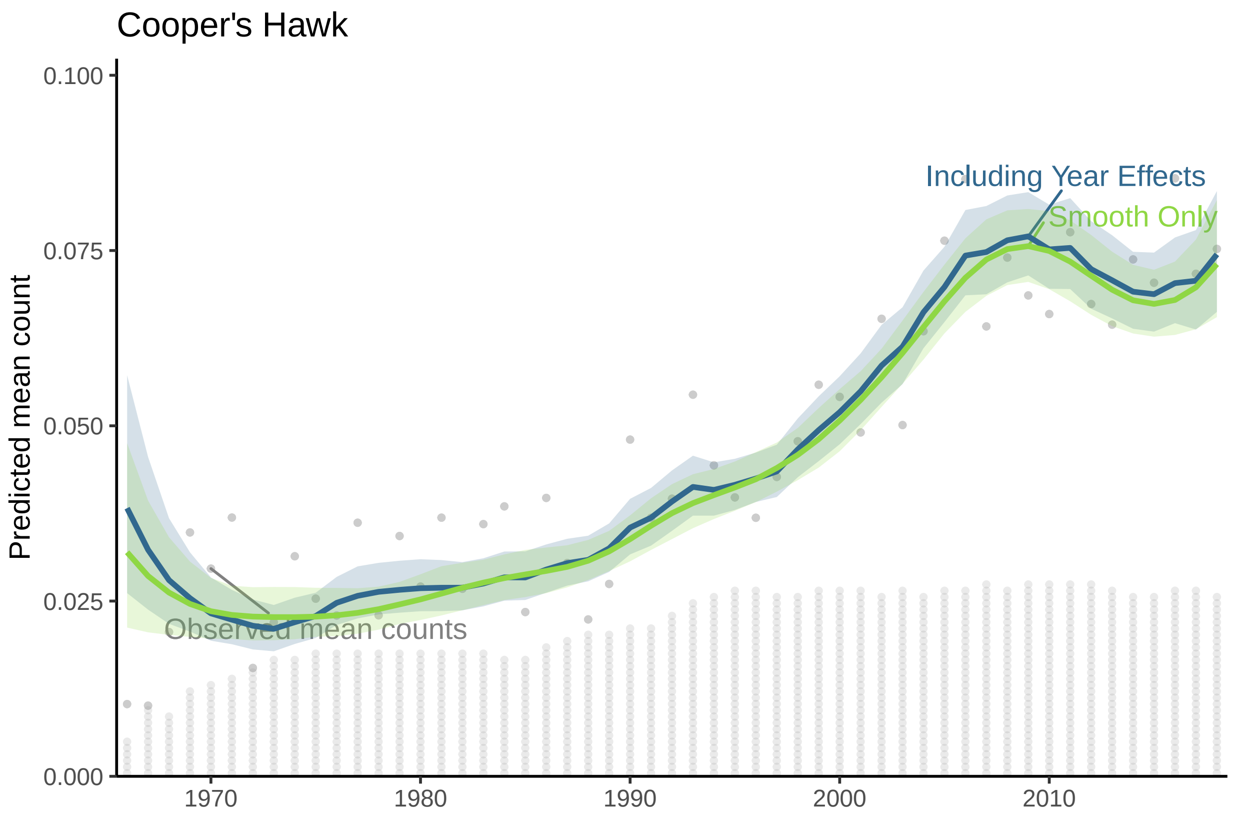


Figure 17: S5.G: Decomposition of the survey-wide population trajectory for Cooper's Hawk from the GAMYE, showing the full trajectory (Including Year Effects) and the isolated smooth component (Smooth Only), which can be used to estimate population trends that are less sensitive to the particular year in which they are estimated. The stacked dots along the x axis indicate the approximate number of BBS counts used in the model; each dot represents 50 counts.

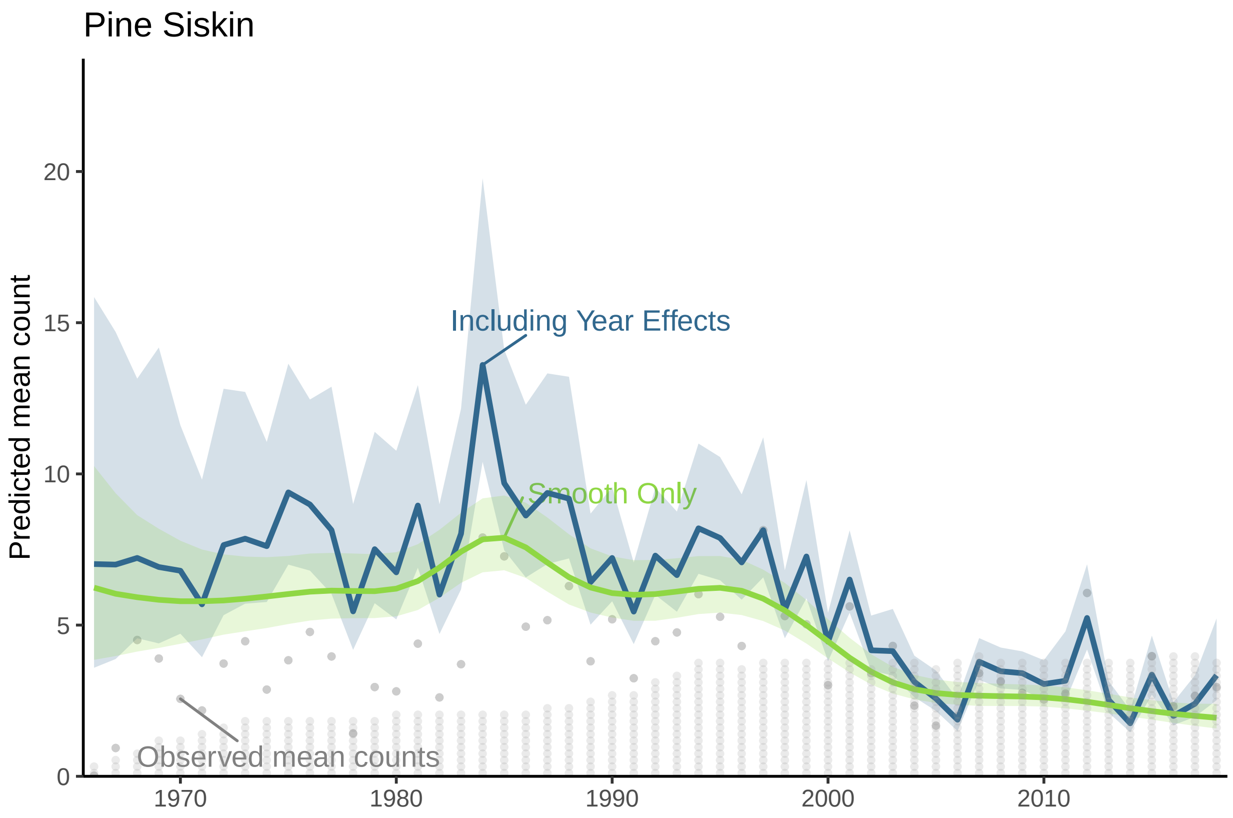


Figure 18: S5.H: Decomposition of the survey-wide population trajectory for Pine Siskin from the GAMYE, showing the full trajectory (Including Year Effects) and the isolated smooth component (Smooth Only), which can be used to estimate population trends that are less sensitive to the particular year in which they are estimated. The stacked dots along the x axis indicate the approximate number of BBS counts used in the model; each dot represents 50 counts.

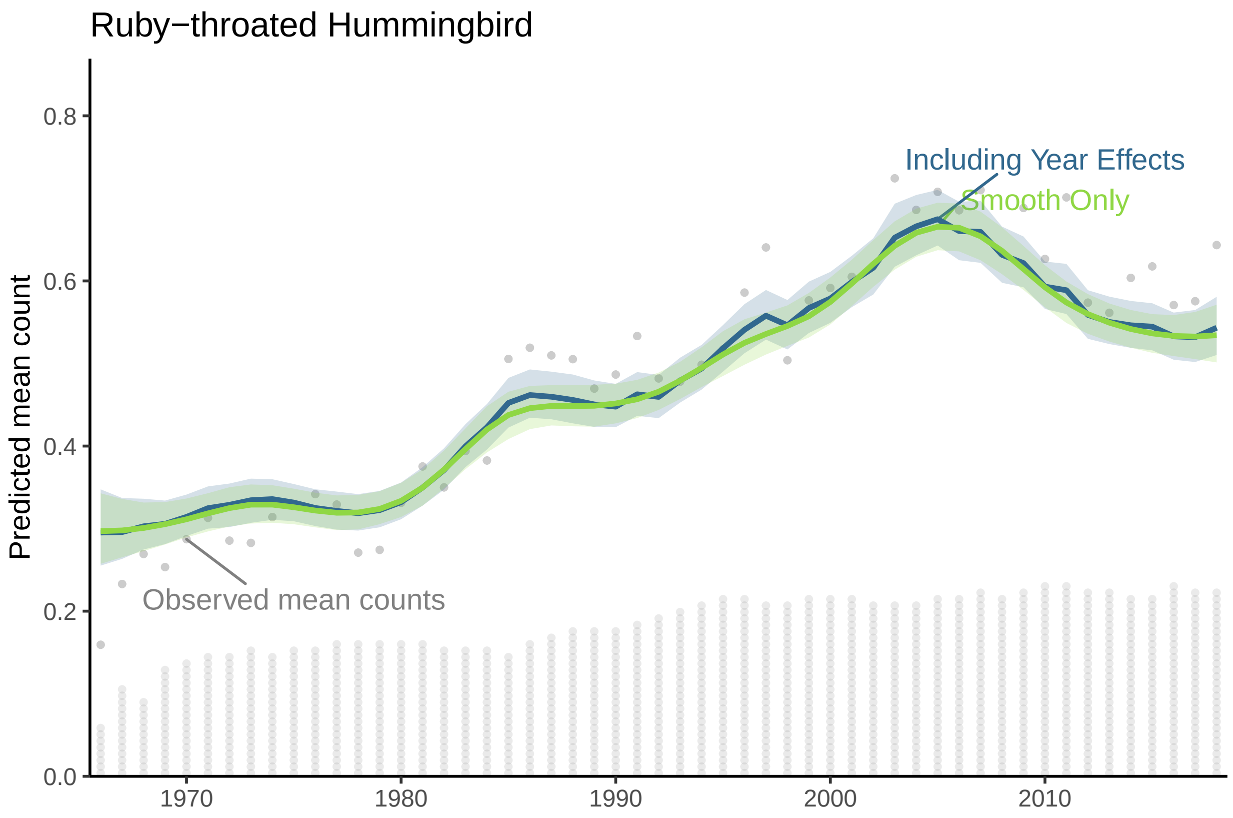


Figure 19: S5.I: Decomposition of the survey-wide population trajectory for Ruby-throated Hummingbird from the GAMYE, showing the full trajectory (Including Year Effects) and the isolated smooth component (Smooth Only), which can be used to estimate population trends that are less sensitive to the particular year in which they are estimated. The stacked dots along the x axis indicate the approximate number of BBS counts used in the model; each dot represents 50 counts.

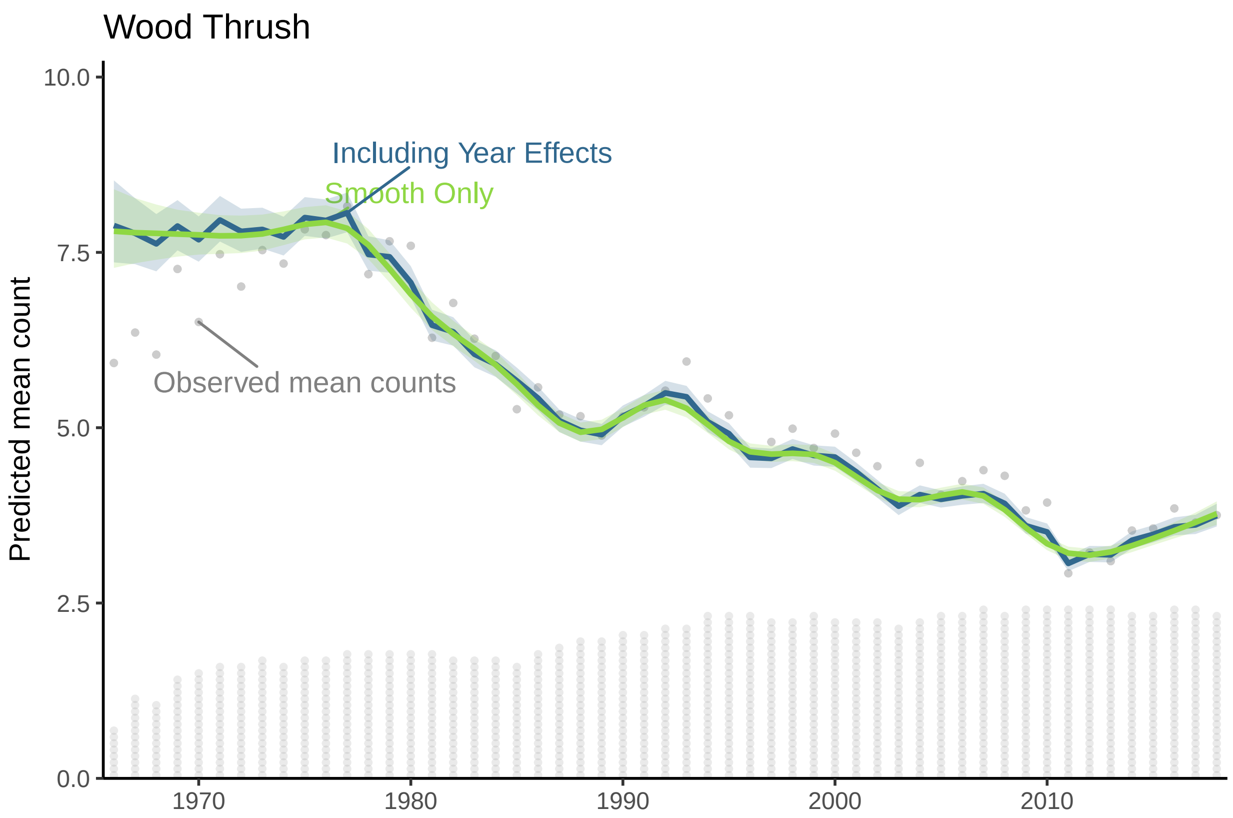
Wood Thrush
Predicted mean count
Including Year Effects
Smooth Only
Observed mean counts
10.0
7.5
5.0
2.5
0.0
1970
1980
1990
2000
2010

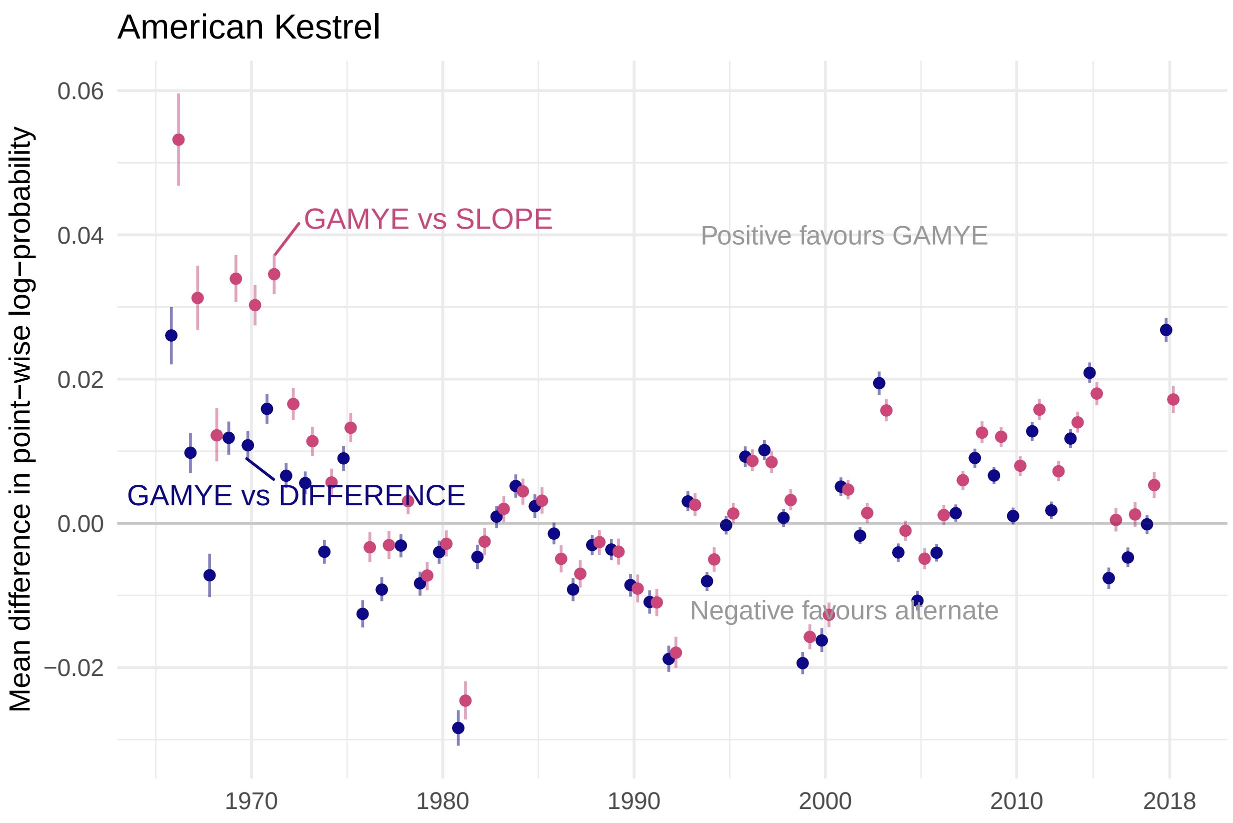


Figure 20: S6.A: Annual differences in predictive fit between the GAMYE and SLOPE (blue) and the GAMYE and DIFFERENCE model (red) for American Kestrel

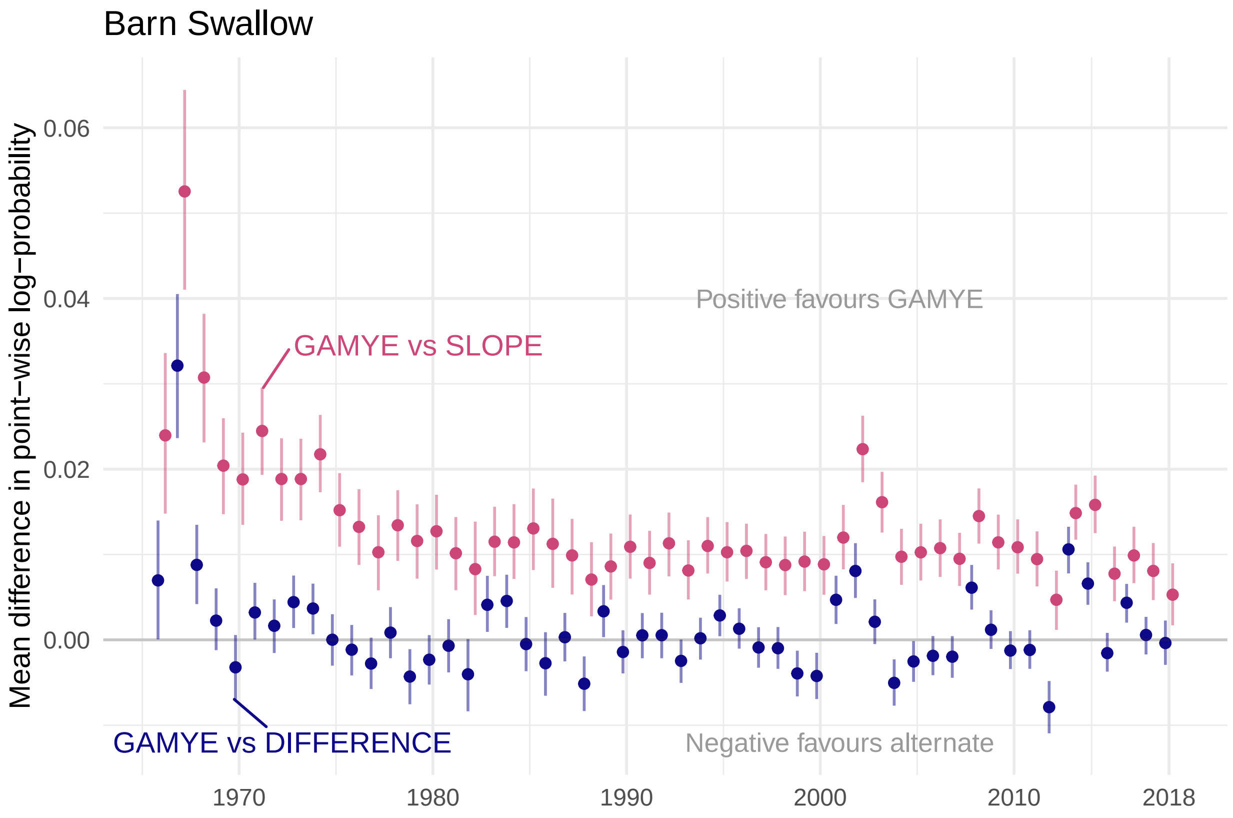


Figure 21: S6.B: Annual differences in predictive fit between the GAMYE and SLOPE (blue) and the GAMYE and DIFFERENCE model (red) for Barn Swallow

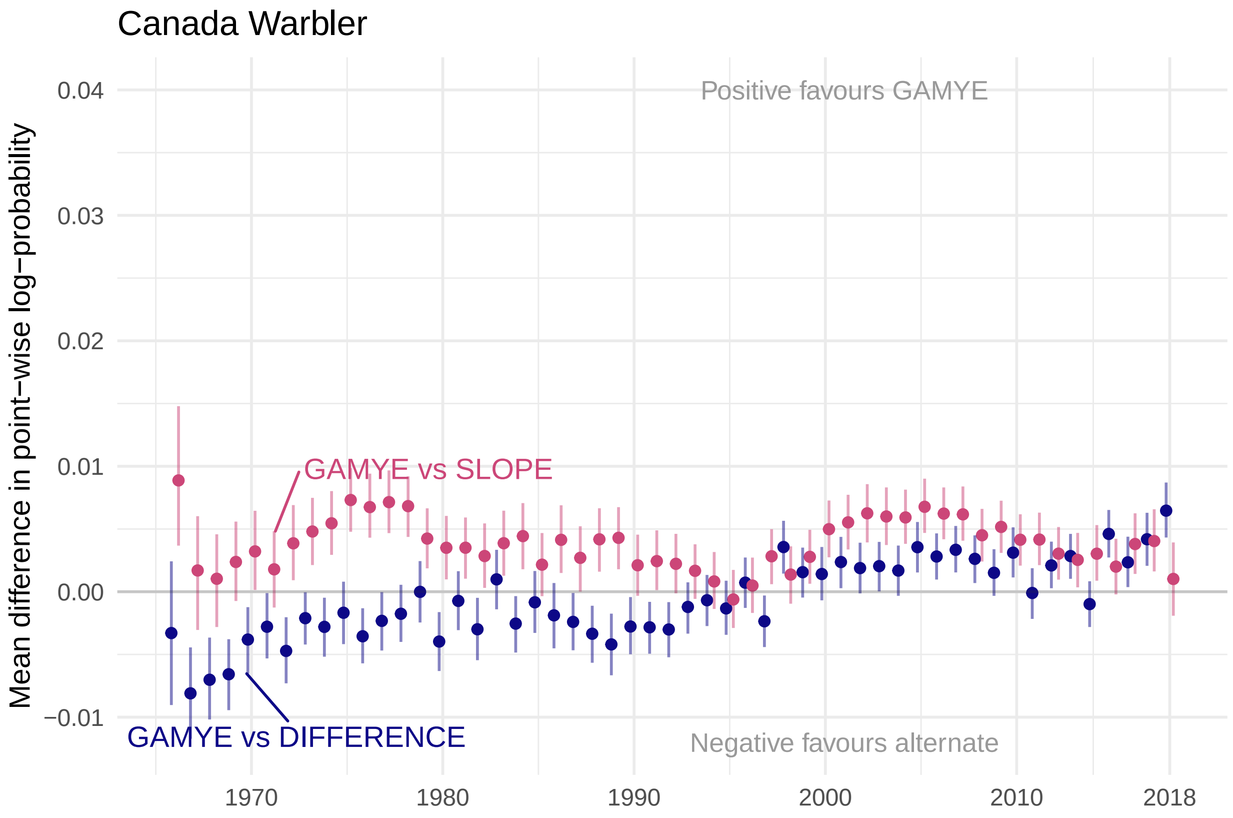


Figure 22: S6.C: Annual differences in predictive fit between the GAMYE and SLOPE (blue) and the GAMYE and DIFFERENCE model (red) for Canada Warbler

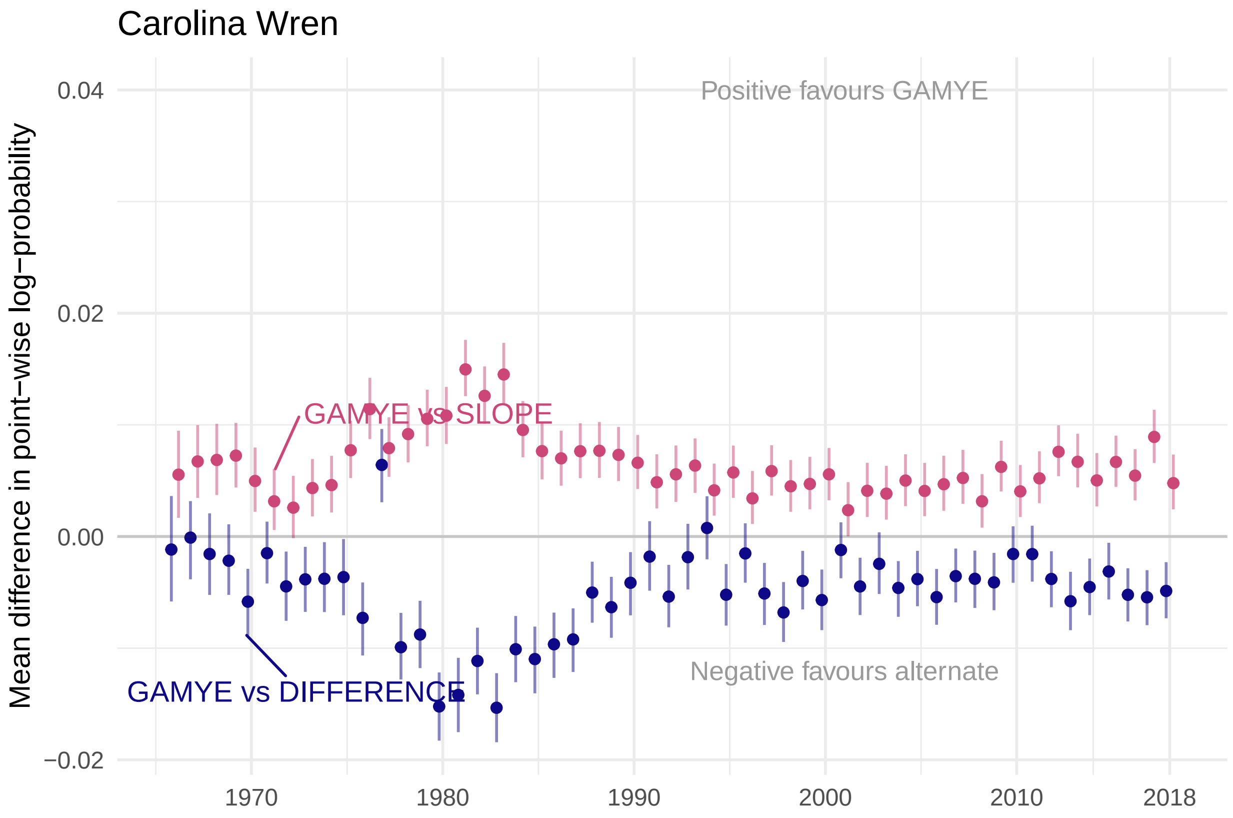


Figure 23: S6.D: Annual differences in predictive fit between the GAMYE and SLOPE (blue) and the GAMYE and DIFFERENCE model (red) for Carolina Wren

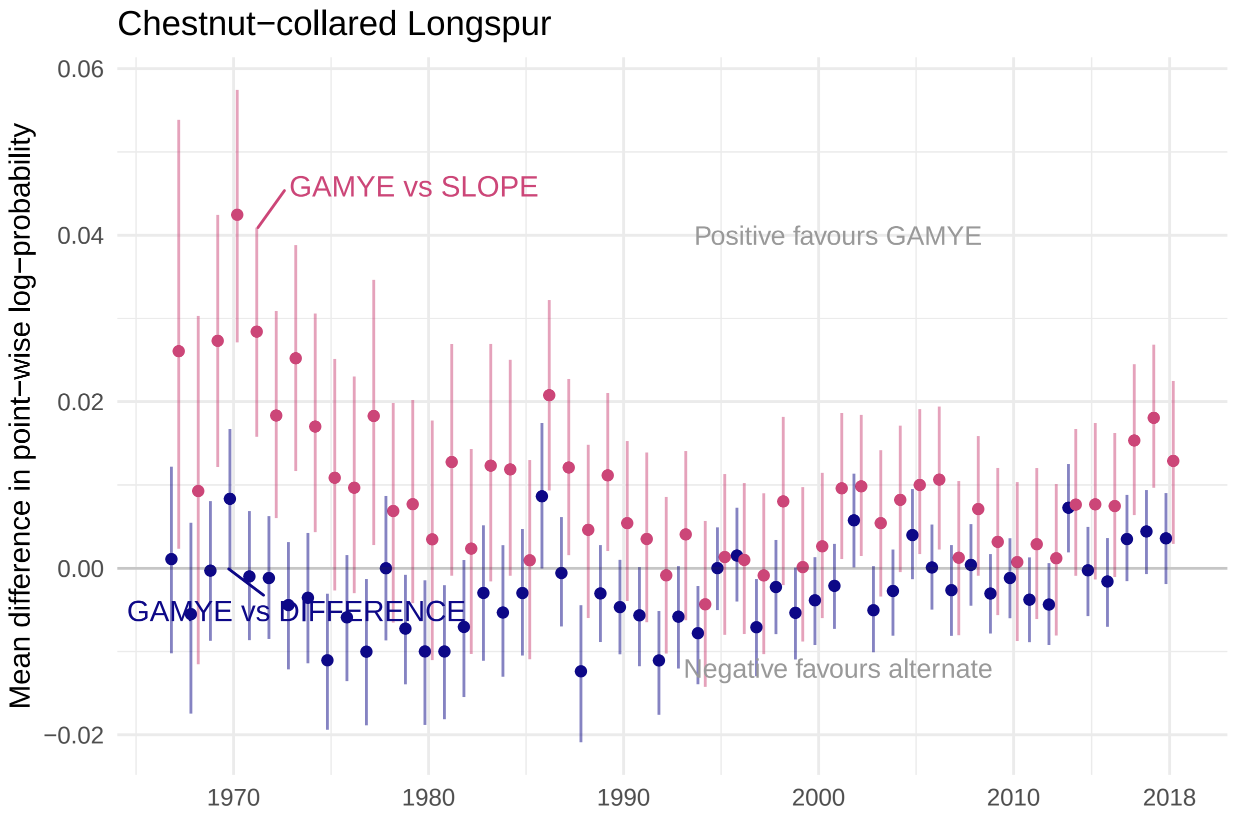


Figure 24: S6.E: Annual differences in predictive fit between the GAMYE and SLOPE (blue) and the GAMYE and DIFFERENCE model (red) for Chestnut-collared Longspur

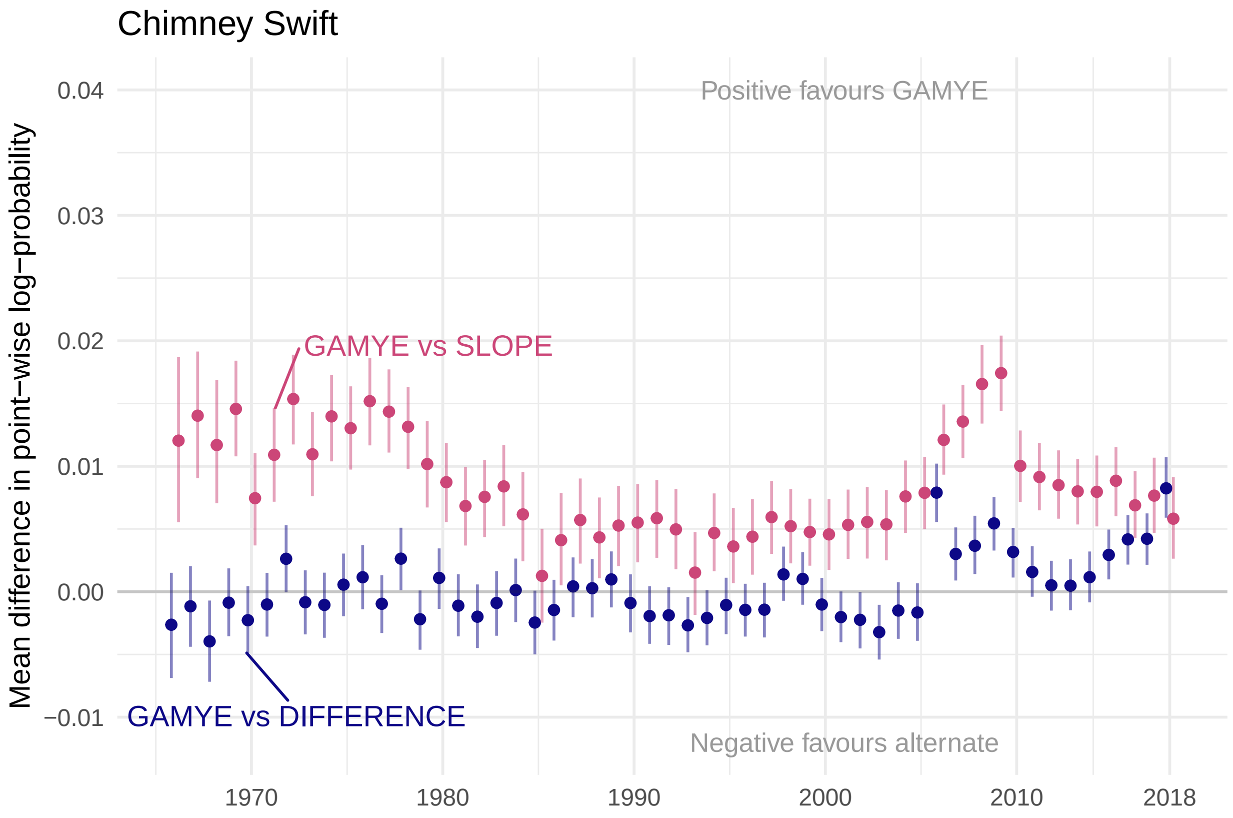


Figure 25: S6.F: Annual differences in predictive fit between the GAMYE and SLOPE (blue) and the GAMYE and DIFFERENCE model (red) for Chimney Swift

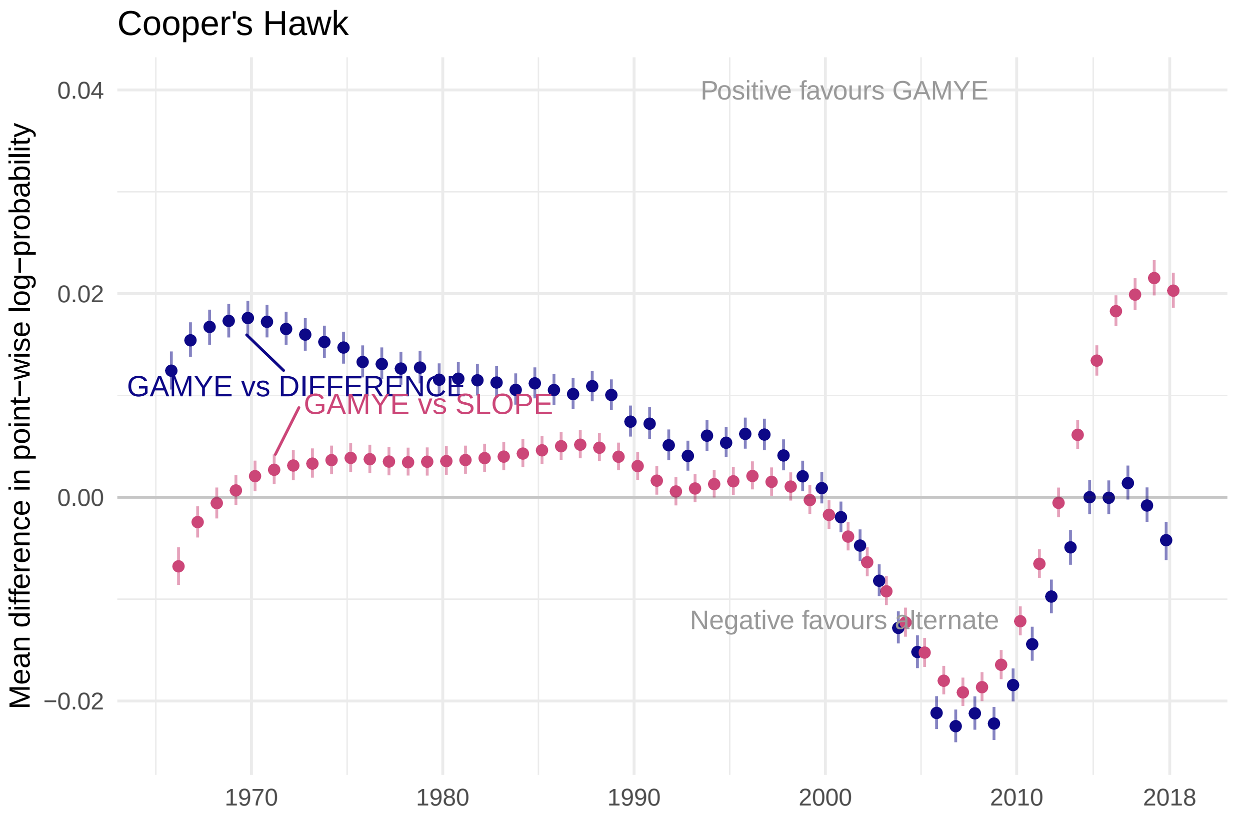


Figure 26: S6.G: Annual differences in predictive fit between the GAMYE and SLOPE (blue) and the GAMYE and DIFFERENCE model (red) for Cooper's Hawk

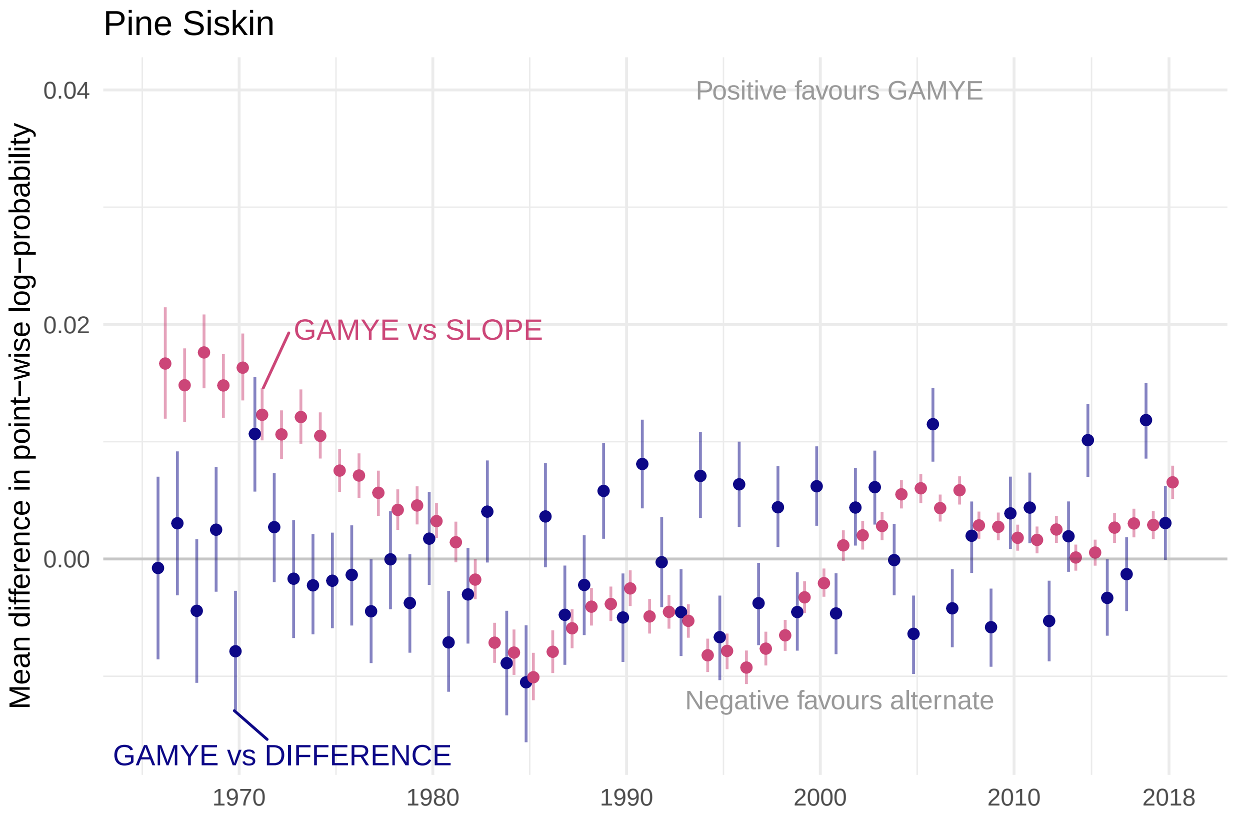


Figure 27: S6.H: Annual differences in predictive fit between the GAMYE and SLOPE (blue) and the GAMYE and DIFFERENCE model (red) for Pine Siskin

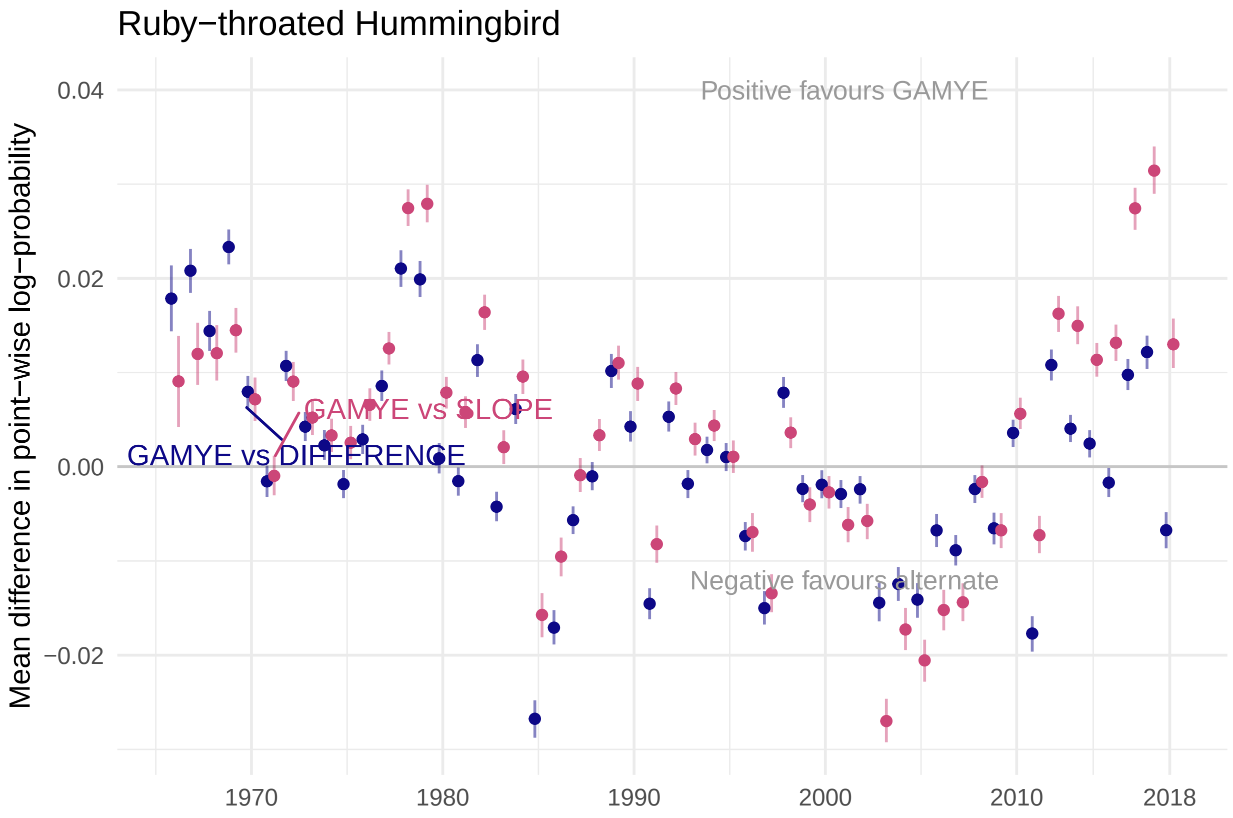


Figure 28: S6.I: Annual differences in predictive fit between the GAMYE and SLOPE (blue) and the GAMYE and DIFFERENCE model (red) for Ruby-throated Hummingbird

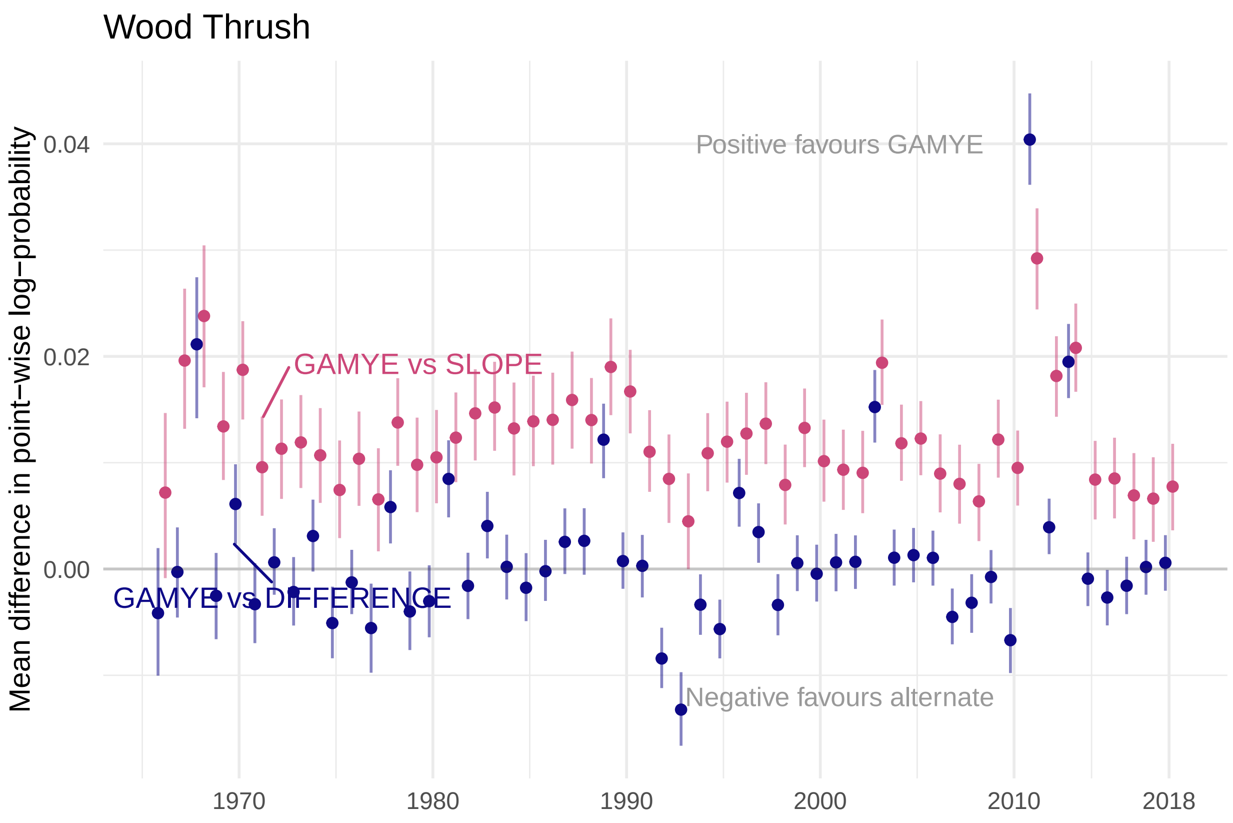
Wood Thrush
Mean difference in point−wise log−probability
0.04
0.02
0.00
Positive favours GAMYE
GAMYE vs SLOPE
GAMYE vs DIFFERENCE
Negative favours alternate
1970
1980
1990
2000
2010
2018

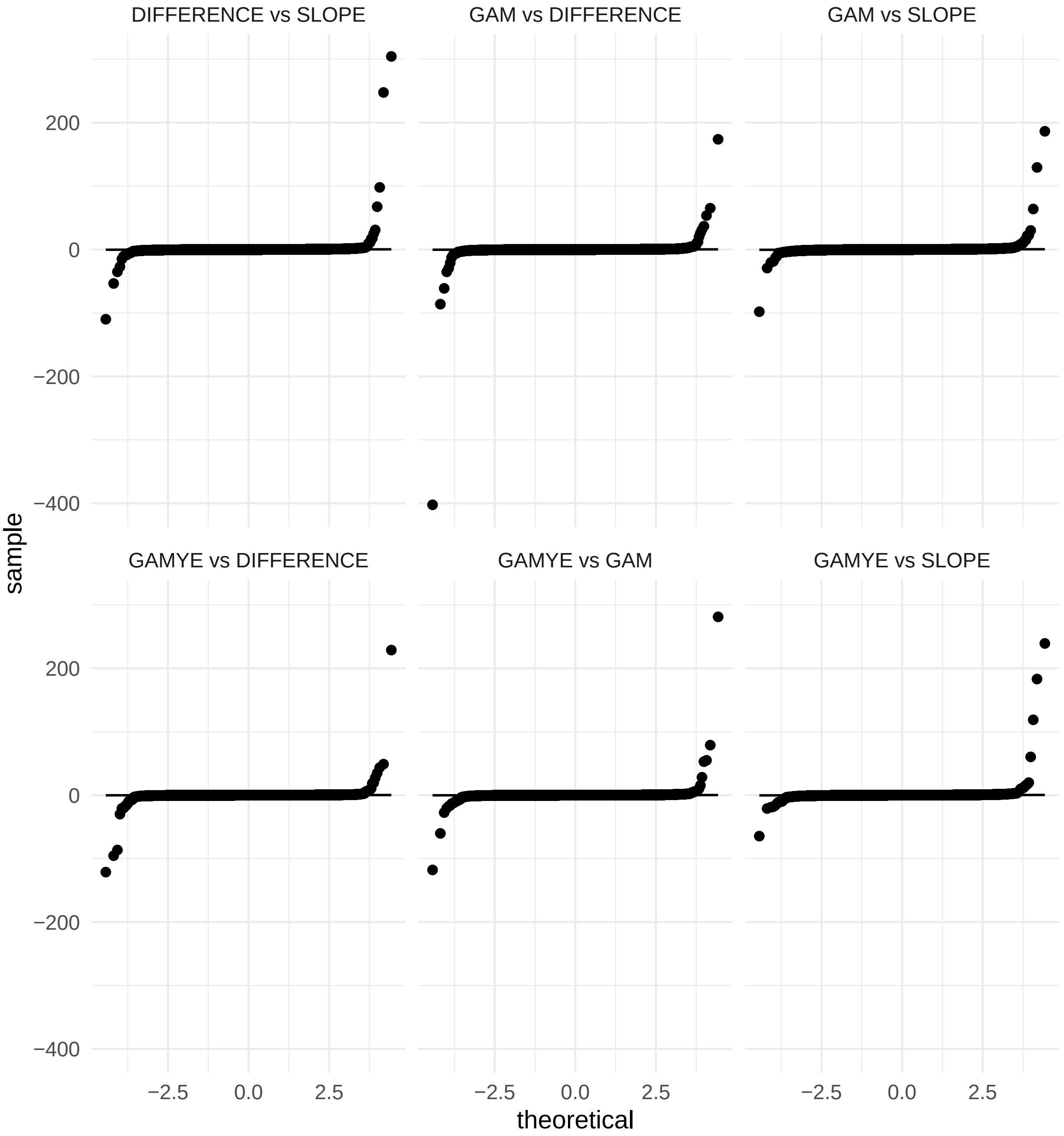


Figure 29: Figure S7: Normal qq plots for the differences in elpd between model pairs for Barn Swallow, demonstrating the non-normal distribution and the heavy tails better estimated using a t-distribution
