## Supplemental Figures S8-S9 for "North American Breeding Bird Survey status and trend estimates to inform a wide-range of conservation needs, using a flexible Bayesian hierarchical generalized additive model"

### Supplemental Material Figures S8 to S9, Smith and Edwards 2020, Improved status and trend estimates from the North American Breeding Bird Survey using a Bayesian hierarchical generalized additive model

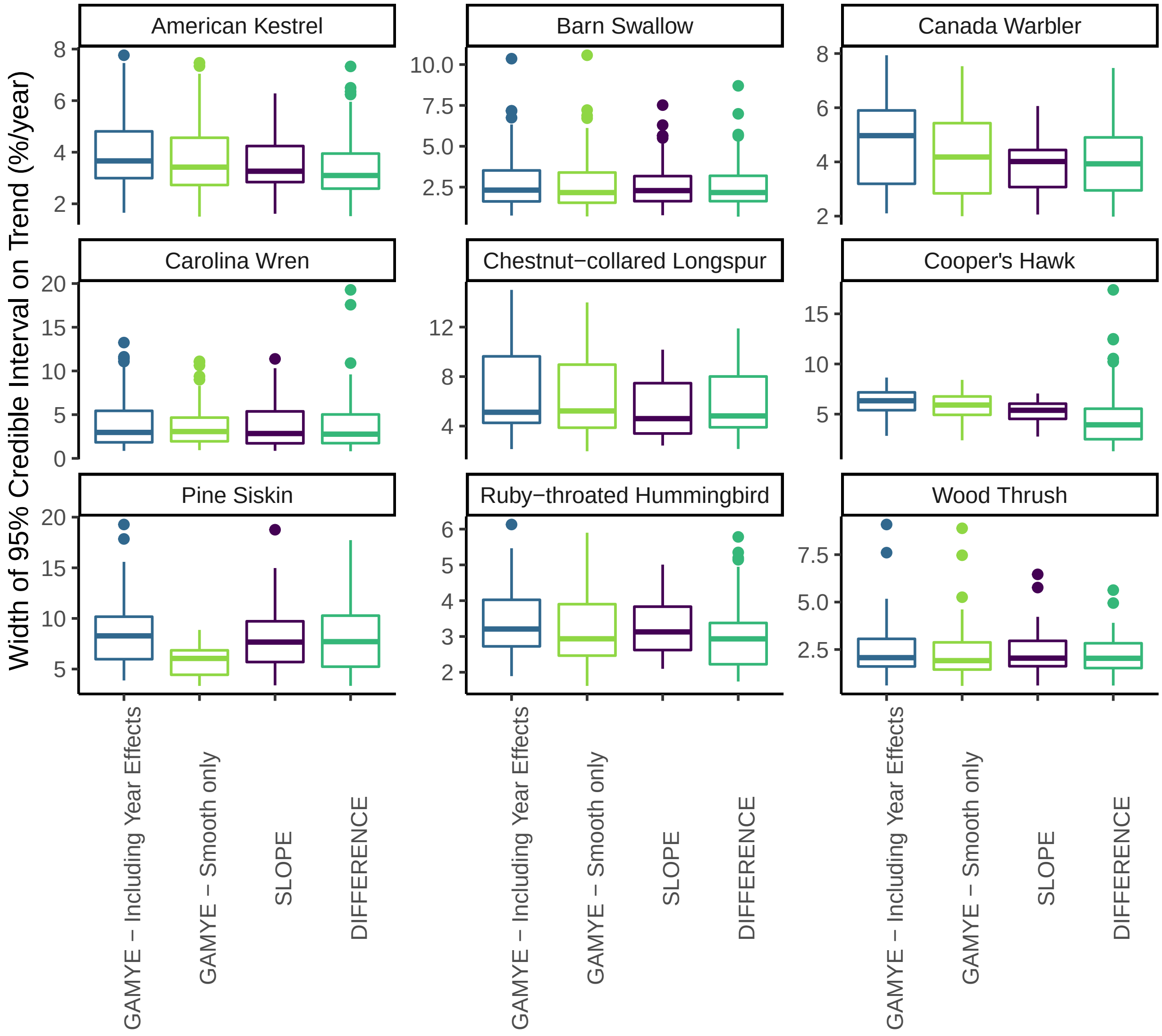


Figure S8. Comparison of the width of the credible intervals for trends (1970-2018) from the GAMYE either including the year-effects or using only the smooth component, compared to the same interval widths for trends from the SLOPE and DIFFERENCE models. The smooth-only trends from the GAMYE model tend to have slightly narrower credible intervals than the trends from the same model that include the year effects. However, the magnitude of the difference in precision is small, and in most cases the GAMYE trends with the smooth component only have CIs that are either similar in width or even larger than the estimates from the SLOPE and DIFFERENCE models. For one species, the width of the credible interval is smaller for the smooth only trends - Pine Siskin. This species' population trajectory is strongly dominated by extreme annual fluctuations and so explicitly modeling these annual fluctuations and removing their influence from the trend calculation results in a much more precise estimate of the average annual rate of change

#### Assessing the prior sensitivity of the GAM hyperparameters

The inverse gamma prior on the variance of the GAM hyperparameters follows the recommended priors in Crainiceanu et al. 2005 (~gamma(0.01,0.0001)). If transformed to the scale of the standard deviation, the 99% percentile of the prior distribution is > 10^100. This prior is intended to be uninformative because it is effectively flat across the entire range of plausible values (< approximately 5). However, this prior also includes values of the variance (or standard deviation) that are far beyond the range of plausible values. The posterior distribution of this variance parameter is largely responsible for the complexity penalty, controlling the degree of smoothing in the GAM, and so if the prior is unintentionally informative, it could result in under-restrictive smoothing penalties. If this "uninformative" prior is leading to under-restrictive smoothing, it would produce population trajectories that over-fit the data and appear more complex (i.e., "wiggly") than warranted. The figures below show that using a much more restrictive prior (~gamma(2,0.2)) that places > 99% of the posterior density for the standard deviation at values < 1.2. For most species, this is a regularizing prior because almost all of the prior density falls below the posterior estimates, which are generally > 2.0. So the two priors are very different, yet as the following figures demonstrate, the posterior estimates of the GAM trajectories are almost identical. The population inferences from the GAMs are almost entirely driven by the model-structures and the data.

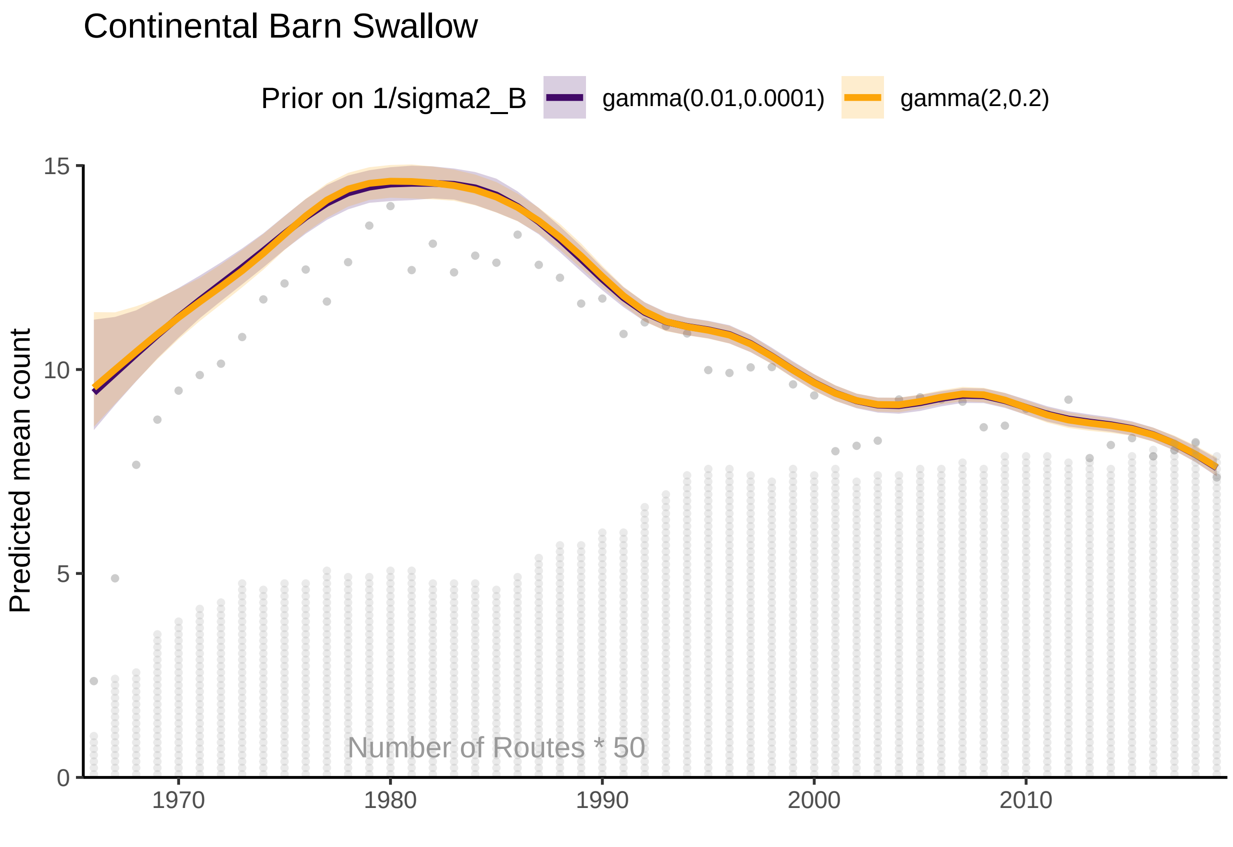


Figure 1: S9.A: Comparison of the continental population trajectories for Barn Swallow from a model for the BBS using a hierarchical GAM smooth with two alternative prior distributions on the variance parameter that provides the complexity penalty for the survey-wide GAM hyperparameters, i.e., the mean continental population trajectory towards which each stratum-level smooth is shrunk

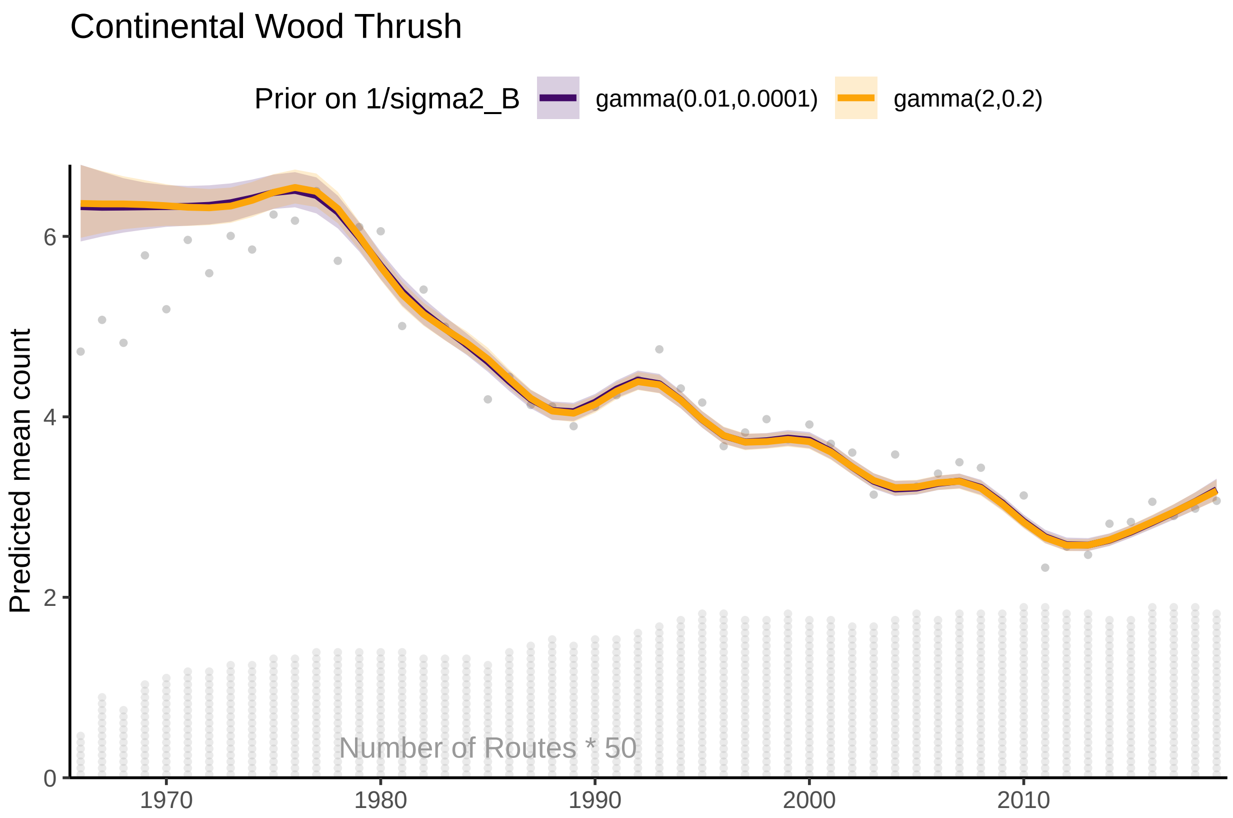


Figure 2: S9.B: Comparison of the continental population trajectories for Wood Thrush from a model for the BBS using a hierarchical GAM smooth with two alternative prior distributions on the variance parameter that provides the complexity penalty for the survey-wide GAM hyperparameters, i.e., the mean continental population trajectory towards which each stratum-level smooth is shrunk

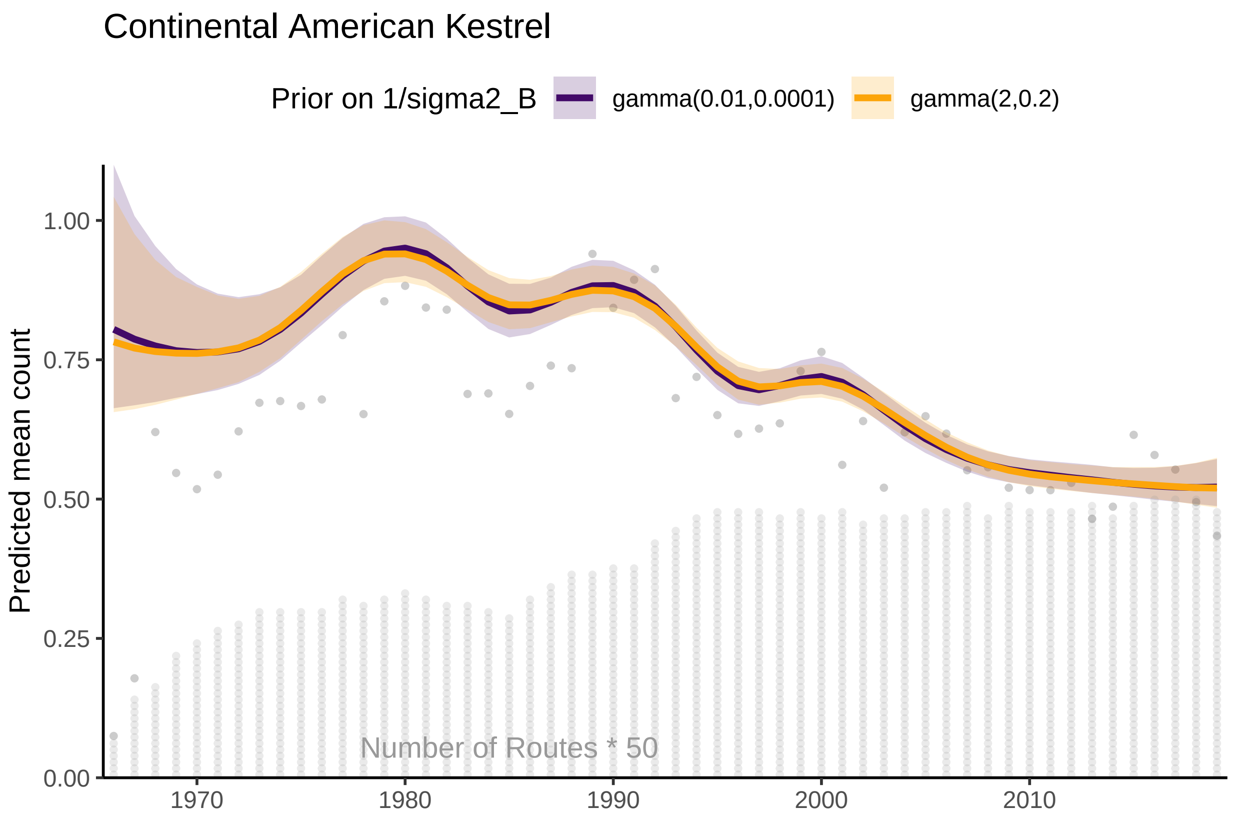


Figure 3: S9.C: Comparison of the continental population trajectories for American Kestrel from a model for the BBS using a hierarchical GAM smooth with two alternative prior distributions on the variance parameter that provides the complexity penalty for the survey-wide GAM hyperparameters, i.e., the mean continental population trajectory towards which each stratum-level smooth is shrunk

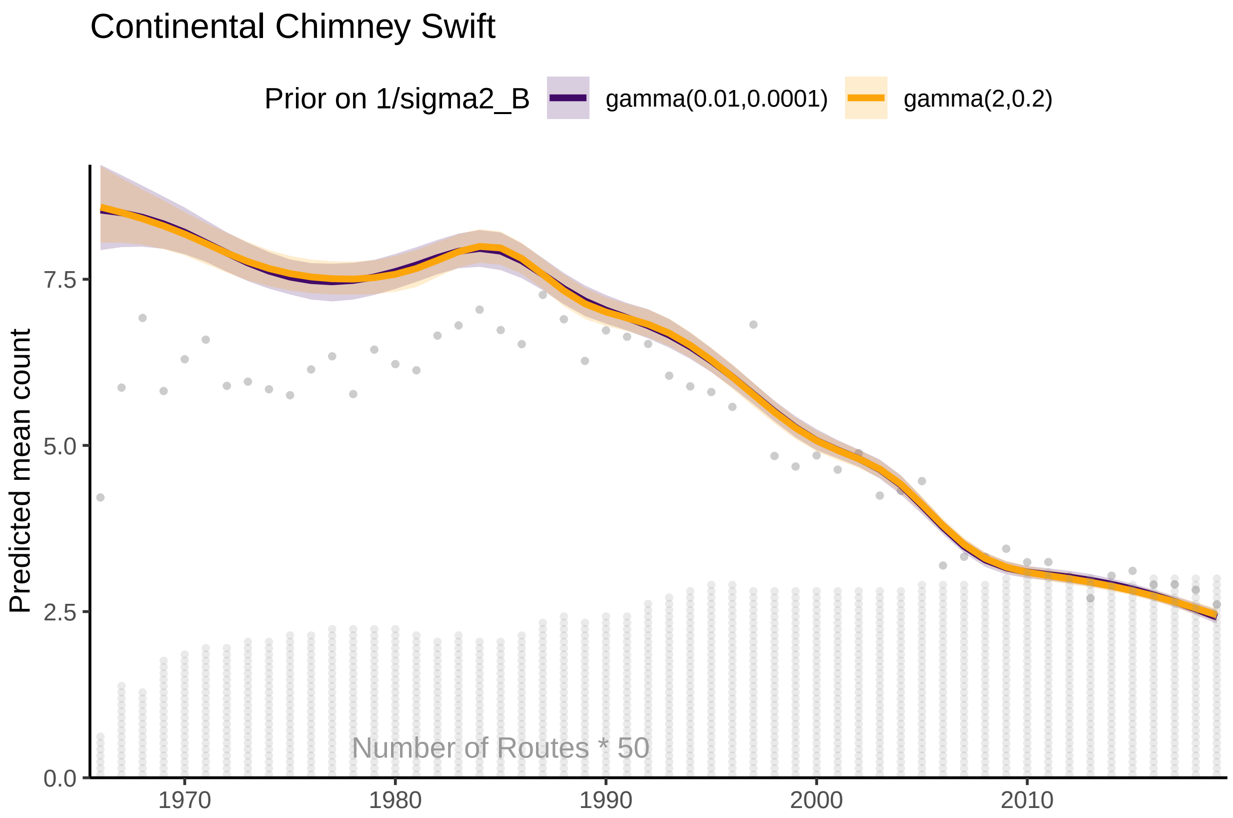


Figure 4: S9.D: Comparison of the continental population trajectories for Chimney Swift from a model for the BBS using a hierarchical GAM smooth with two alternative prior distributions on the variance parameter that provides the complexity penalty for the survey-wide GAM hyperparameters, i.e., the mean continental population trajectory towards which each stratum-level smooth is shrunk

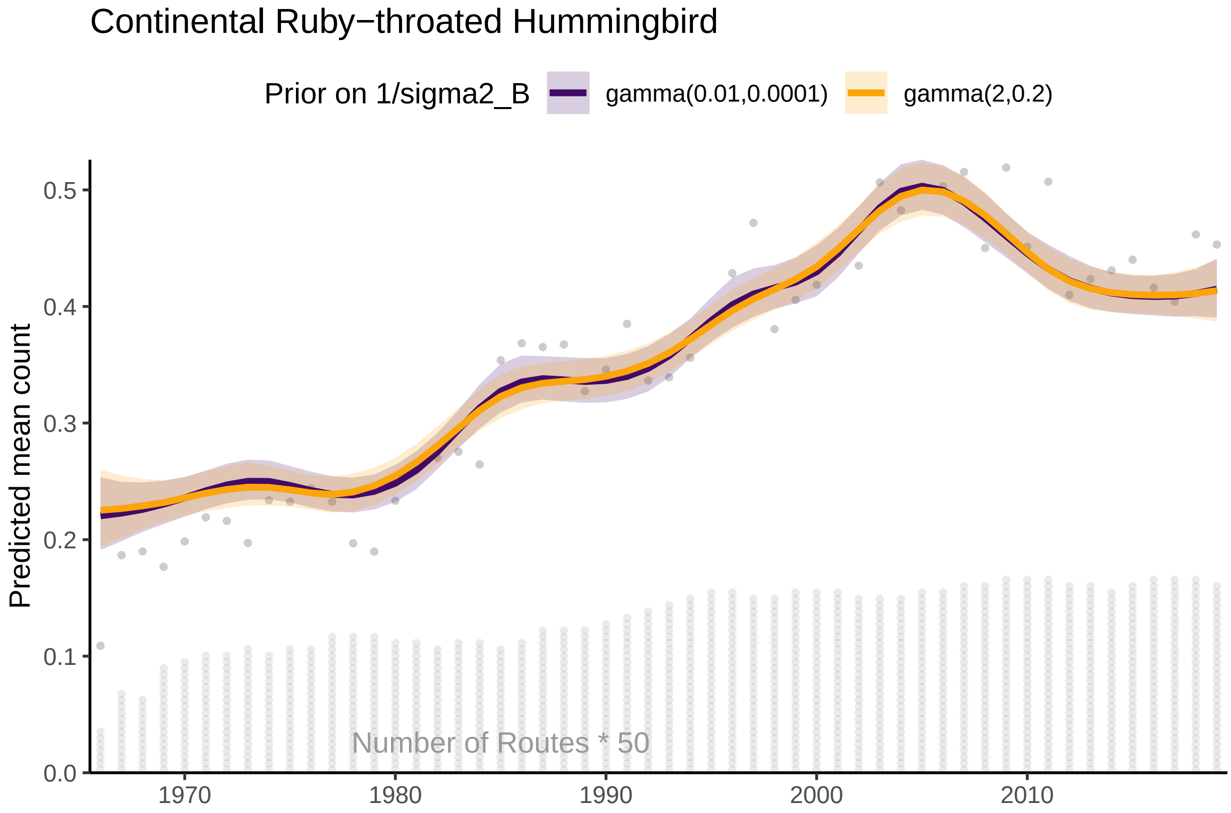


Figure 5: S9.E: Comparison of the continental population trajectories for Ruby-throated Hummingbird from a model for the BBS using a hierarchical GAM smooth with two alternative prior distributions on the variance parameter that provides the complexity penalty for the survey-wide GAM hyperparameters, i.e., the mean continental population trajectory towards which each stratum-level smooth is shrunk

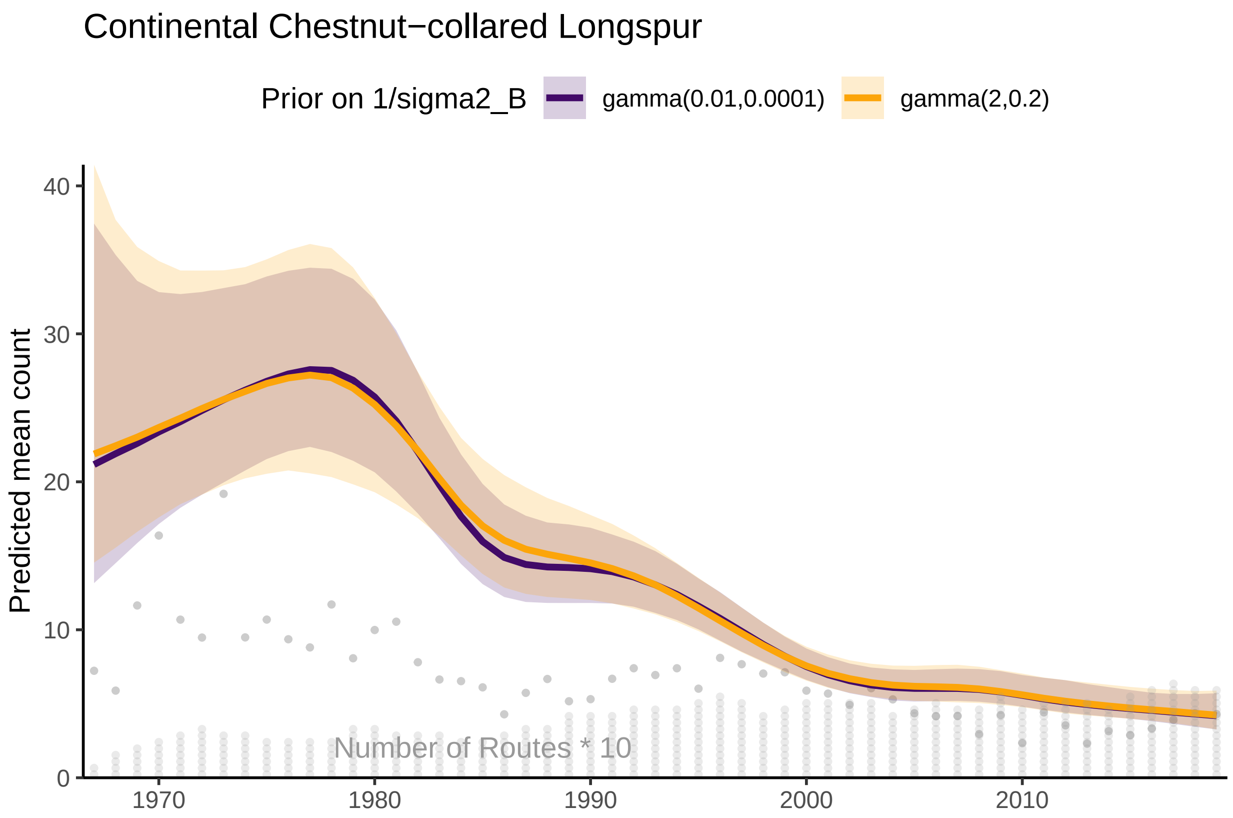


Figure 6: S9.F: Comparison of the continental population trajectories for Chestnut-collared Longspur from a model for the BBS using a hierarchical GAM smooth with two alternative prior distributions on the variance parameter that provides the complexity penalty for the survey-wide GAM hyperparameters, i.e., the mean continental population trajectory towards which each stratum-level smooth is shrunk

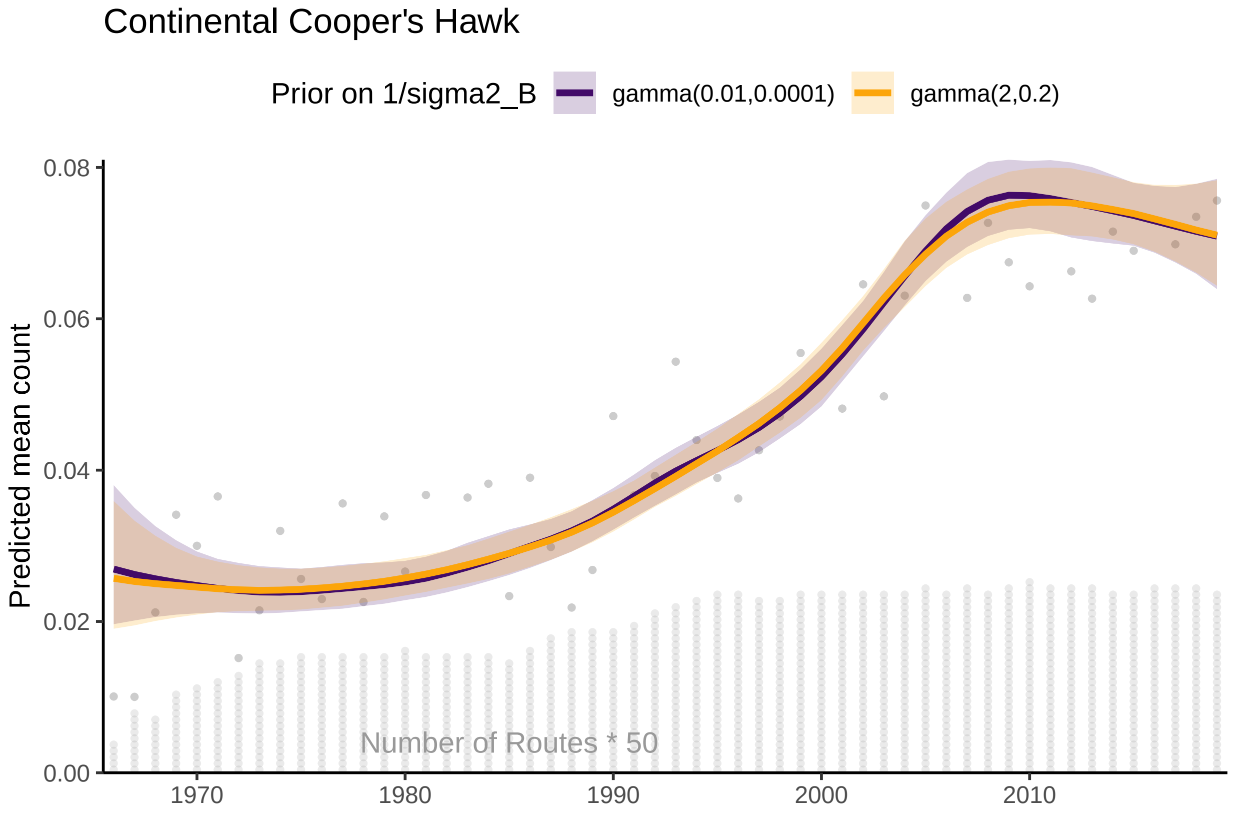


Figure 7: S9.G: Comparison of the continental population trajectories for Cooper's Hawk from a model for the BBS using a hierarchical GAM smooth with two alternative prior distributions on the variance parameter that provides the complexity penalty for the survey-wide GAM hyperparameters, i.e., the mean continental population trajectory towards which each stratum-level smooth is shrunk

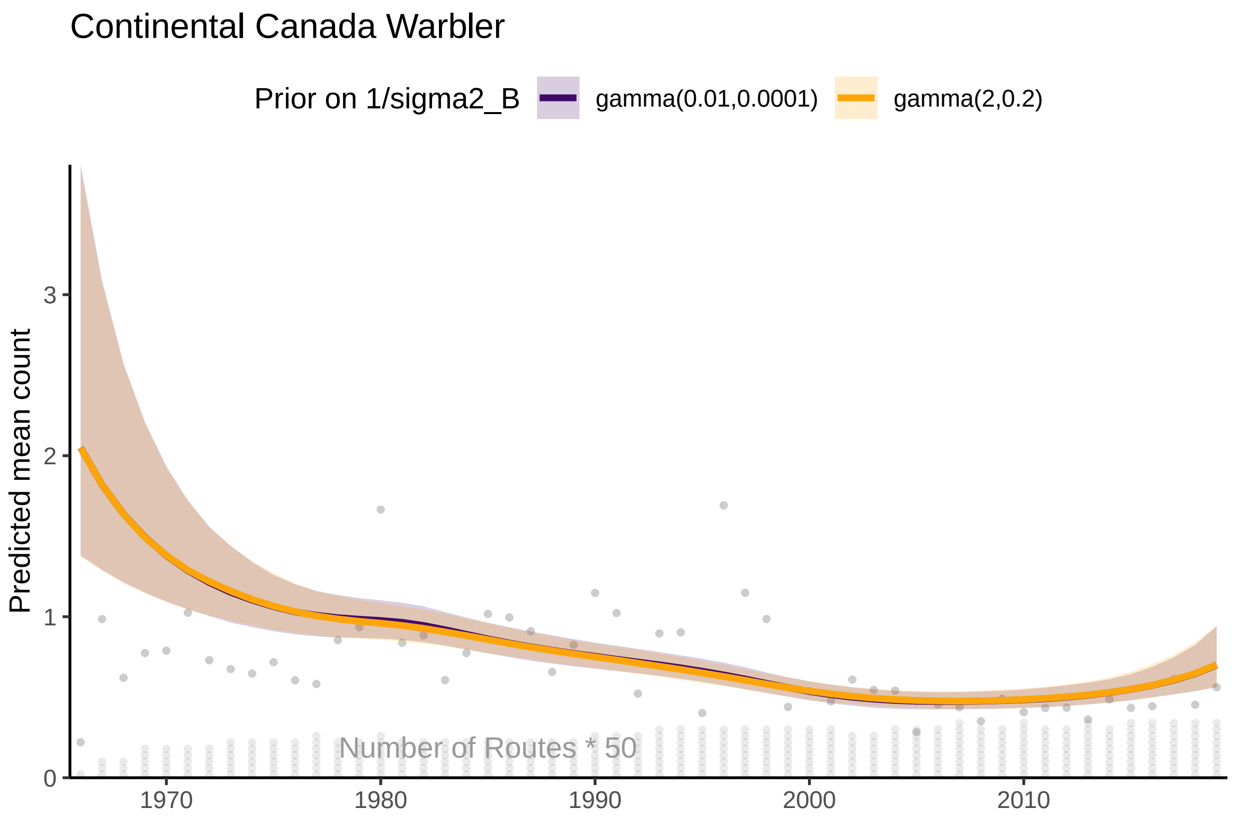


Figure 8: S9.H: Comparison of the continental population trajectories for Canada Warbler from a model for the BBS using a hierarchical GAM smooth with two alternative prior distributions on the variance parameter that provides the complexity penalty for the survey-wide GAM hyperparameters, i.e., the mean continental population trajectory towards which each stratum-level smooth is shrunk

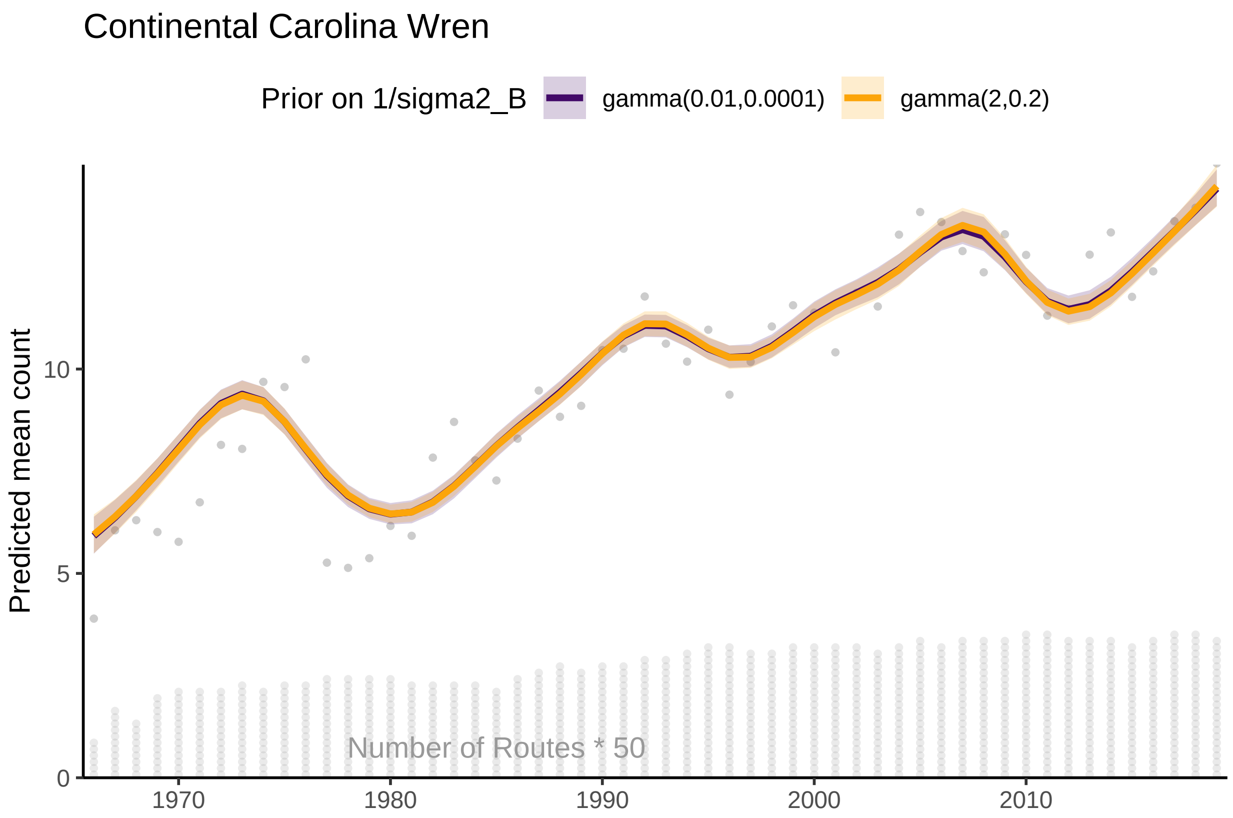


Figure 9: S9.I: Comparison of the continental population trajectories for Carolina Wren from a model for the BBS using a hierarchical GAM smooth with two alternative prior distributions on the variance parameter that provides the complexity penalty for the survey-wide GAM hyperparameters, i.e., the mean continental population trajectory towards which each stratum-level smooth is shrunk

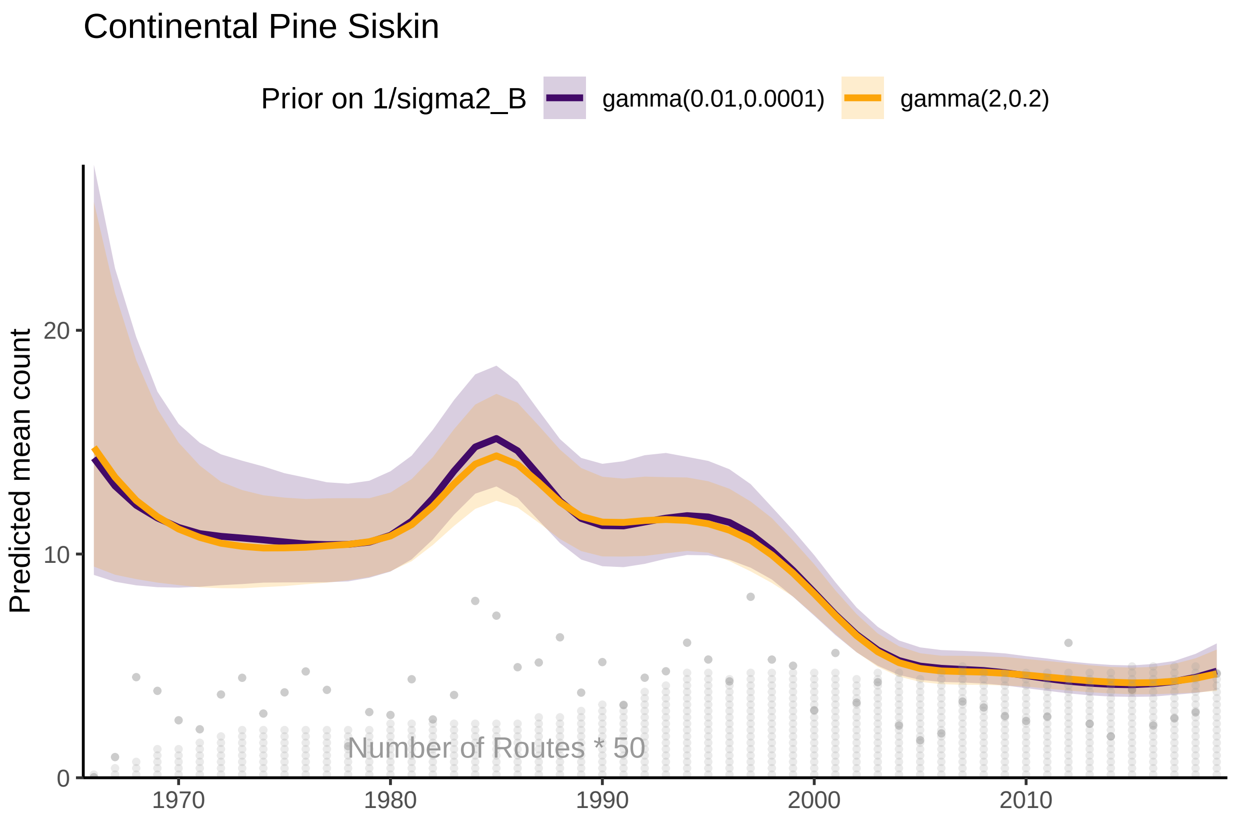


Figure 10: S9.J: Comparison of the continental population trajectories for Pine Siskin from a model for the BBS using a hierarchical GAM smooth with two alternative prior distributions on the variance parameter that provides the complexity penalty for the survey-wide GAM hyperparameters, i.e., the mean continental population trajectory towards which each stratum-level smooth is shrunk
